# Synergism of interferon-beta with antiviral drugs against SARS-CoV-2 variants

**DOI:** 10.1101/2022.07.22.501169

**Authors:** Denisa Bojkova, Richard Stack, Tamara Rothenburger, Joshua D Kandler, Sandra Ciesek, Mark N. Wass, Martin Michaelis, Jindrich Cinatl

## Abstract

Omicron BA.1 variant isolates were previously shown to replicate less effectively in interferon-competent cells and to be more sensitive to interferon treatment than a Delta isolate. Here, an Omicron BA.2 isolate displayed intermediate replication patterns in interferon-competent Caco-2-F03 cells when compared to BA.1 and Delta isolates. Moreover, BA.2 was less sensitive than BA.1 and similarly sensitive as Delta to betaferon treatment. Delta and BA.1 displayed similar sensitivity to the approved anti-SARS-CoV-2 drugs remdesivir, nirmatrelvir, EIDD-1931 (the active metabolite of molnupiravir) and the protease inhibitor aprotinin, whereas BA.2 was less sensitive than Delta and BA.1 to EIDD-1931, nirmatrelvir and aprotinin. Nirmatrelvir, EIDD-1931, and aprotinin (but not remdesivir) exerted synergistic antiviral activity in combination with betaferon, with some differences in the extent of synergism detected between the different SARS-CoV-2 variants. In conclusion, even closely related SARS-CoV-2 (sub)variants can differ in their biology and in their response to antiviral treatments. Betaferon combinations with nirmatrelvir and, in particular, with EIDD-1931 and aprotinin displayed high levels of synergism, which makes them strong candidates for clinical testing. Notably, effective antiviral combination therapies are desirable, as a higher efficacy is expected to reduce resistance formation.

## Introduction

SARS-CoV-2, the coronavirus that causes COVID-19, has caused a pandemic starting in December 2019 [Forchette et al., 2021]. This pandemic has been driven by different SARS-CoV-2 variants that subsequently replaced each other. The original Wuhan strain, was replaced by the Alpha variant (B.1.1.7), which was later replaced by the Beta and P.1 variants in some parts of the world, before the Delta became the dominant variant [Forchette et al., 2021]. Most recently, the Omicron (B.1.1.529, BA.1) variant took over from Delta, which keeps evolving into further subvariants such as BA.2, BA.2.12.1, BA.3, BA.4, and BA.5 [Kawaoka et al., 2022; Sullivan et al., 2022].

SARS-CoV-2 evolution is at least in part driven by the selection pressure induced by previous infections and vaccinations. In agreement, Omicron subvariants display the greatest propensity to infect individuals with pre-existing vaccine- or infection-mediated immunity [Bruel et al., 2022; Quandt et al., 2022]. Despite this immune evasion capacity, the available vaccines, which are based on the original Wuhan strain, still provide significant protection from severe COVID-19 [Accorsi et al., 2022: Andrews et al., 2022].

There is also concern that new SARS-CoV-2 variants may change in their susceptibility to antiviral drugs. We have previously shown that SARS-CoV-2 and the closely related SARS-CoV differ in their drug sensitivity profiles [Bojkova et al., 2021]. However, different SARS-CoV-2 variants including Omicron BA.1 and BA.2 have so far displayed comparable sensitivity to the approved anti-SARS-CoV-2 drugs remdesivir (RNA-dependent RNA polymerase inhibitor), molnupiravir (induces ‘lethal mutagenesis’ during virus replication), and nirmatrelvir (inhibitor of the SARS-CoV-2 main/ 3CL protease) [Bojkova et al., 2022, Kawaoka et al., 2022; Takashita et al., 2022; Takashita et al., 2022b; Vangeel et al., 2022].

Host cell interferon signalling is crucial for the control of SARS-CoV-2 replication and avoiding severe COVID-19, as indicated by the high vulnerability of individuals with defects in this innate immune response mechanism [Bastard et al., 2020; Hadjadj et al., 2020; Zhang et al., 2020]. Despite this importance for SARS-CoV-2 pathogenicity, interferons were not effective in initial clinical trials for the treatment of COVID-19 [Bhushan et al., 2021; Li et al., 2021; Monk et al., 2021; WHO Solidarity Trial Consortium, 2021]. However, we found that Omicron variant BA.1 isolates were substantially more sensitive to interferon treatment than a Delta isolate [Bojkova et al., 2022a].

Based on these findings, we here systematically compared the sensitivity of Delta, BA.1, and BA.2 isolates to betaferon (a clinically approved interferon-β preparation) alone or in combination with the approved anti-SARS-CoV-2 drugs remdesivir, molnupiravir, and nirmatrelvir [Ho et al., 2022]. Moreover, we included aprotinin in this study, a protease inhibitor that we have shown to inhibit SARS-CoV-2 replication at least in part by interfering with the cleavage and activation of the viral spike (S) protein by host cell proteases [Bojkova et al., 2020; Bojkova et al., 2022] and that was recently reported to be effective in COVID-19 patients in a clinical trial [Redondo-Calvo et al., 2022].

## Methods

### Analysis of sequence variants

Amino acid sequences of the SARS-CoV-2 isolates FFM-SIM0550 (Omicron BA.1, GenBank ID: OL800702), FFM-BA.2-3833 (Omicron BA.2, GenBank ID: OM617939), and FFM-IND8424 (Delta/ B.1.617.2, GenBank ID: MZ315141) were obtained and aligned using the NCBI Virus tool (https://www.ncbi.nlm.nih.gov/labs/virus/). The mutation prevalence in these isolates was compared to their prevalence across the lineage using https://www.outbreak.info. The potential significance of individual mutations was assessed relative to the Wuhan reference strain by determining Blosum80 scores (https://www.rdocumentation.org/packages/peptider/versions/0.2.2/topics/BLOSUM80), evolutionary conservation using ConSurf [Ashkenazy et al., 2016], and the potential impact of mutation on protein stability using the mCSM-PPI2 server [Rodrigues et al., 2019].

Changes to residue bonding were visualised using Covid-3D [Portelli et al., 2020] and Pymol (https://pymol.org/2/). Annotated protein structures were created from existing structures obtained from the Protein Databank in Europe (PDBe) [PDBe-KB consortium, 2022] Covid-19 data portal (https://www.ebi.ac.uk/pdbe/covid-19) or modelled using AlphaFold [Jumper et al., 2021].

### Cell culture

The Caco-2 subline Caco-2-F03 [Cinatl et al., 2004; Hoehl et al., 2020; Bojkova et al., 2021; Bojkova et al., 2022b] (derived from the Resistant Cancer Cell Line (RCCL) collection [Michaelis et al., 2019]), Vero (DSMZ, Braunschweig, Germany), Calu-3 (ATCC, Manassas, VA, US) were grown at 37 °C in minimal essential medium (MEM) supplemented with 10% fetal bovine serum (FBS), 100 IU/mL of penicillin, and 100 μg/mL of streptomycin. All culture reagents were purchased from Sigma-Aldrich.

### Virus preparation

The SARS-CoV-2 isolates Omicron BA.1 (B.1.1.529: FFM-SIM0550/2021, EPI_ISL_6959871, GenBank ID OL800702), Omicron BA.2 (B.1.1.529.2: FFM-BA.2-3833, GenBank ID OM617939), and Delta (B.1.167.2: FFM-IND8424/2021, GenBank ID MZ315141) were cultivated in Caco-2 cells as previously described [Cinatl et al., 2004; Hoehl et al., 2020; Bojkova et al., 2021] and stored at –80°C.

### Determination of infectious titres

Caco-2-F03 cells were infected with SARS-CoV-2 variants at MOI of 1 for 1h. After the incubation period, the infectious inoculum was removed, cells were washed with PBS and supplemented with fresh medium. One day later, supernatants were collected and stored at -80°C upon titration. Infectious titres were determined by serial dilutions of cell culture supernatants on confluent layers of Caco-2 cells in 96-well plates and expressed as TCID50/ml.

### Immunofluorescence staining

The cells were fixed at indicated times with 3% PFA permeabilized with 0.1 % Triton X-100. Prior to primary antibody labeling, cells were blocked with 5% donkey serum in PBS or 1% BSA and 2% goat serum in PBS for 30 minutes at RT. Spike protein was detected by primary antibody (1:1500, Sinobiological) followed by Alexa Fluor 647 anti-rabbit secondary antibody (1:1000, Invitrogen). The nucleus was labelled using DAPI (1:1000, Thermo Scientific). The images were taken by Spark^®^ Mulitmode microplate reader (TECAN) at 4x magnification.

### Immunostaining

Cells were fixed with acetone:methanol (40:60) solution and immunostaining was performed using a monoclonal antibody directed against the spike protein of SARS-CoV-2 (1:1500, Sinobiological), which was detected with a peroxidase-conjugated anti-rabbit secondary antibody (1:1000, Dianova), followed by addition of AEC substrate. The spike positive area was scanned and quantified by the Bioreader® 7000-F-Z-I microplate reader (Biosys). The results are expressed as percentage of inhibition relative to virus control which received no drug.

### Antiviral assay

Confluent layers of cells in 96-well plates were treated with decreasing concentrations of interferon-β (betaferon, Bayer), remdesivir (Selleckchem), nirmatrelvir (Selleckchem), molnupiravir (Selleckchem) and/ or aprotinin (Sigma-Aldrich) subsequently infected with SARS-CoV-2 at an MOI of 0.01. In experiments with remdesivir and nirmatrelvir, 1 µM of the ABCB1 inhibitor Zosuquidar (Selleckchem) was added. Antiviral effects were determined by immunostaining for the SARS-CoV-2 spike (S) protein 24 h post infection.

To evaluate antiviral activity of interferon-β in a combination with remdesivir, nirmatrelvir, EIDD-1931 (Selleckchem), or aprotinin the agents were tested alone or in fixed combinations at 1:2 dilutions using monolayers of Caco-2 cells infected SARS-CoV-2 isolates at MOI 1 24 h post infection. The calculation of IC_50_, IC_75_, IC_90_ and IC_95_ for single drugs and their combinations as well as combination indexes (CIs) was performed using the software CalcuSyn (Biosoft) based on the method of Chou and Talalay [Chou, 2006]. The weighted average CI value (CI_wt_) was calculated according to the formula: CI_wt_ [CI_50_ + 2CI_75_ + 3CI_90_ + 4CI_95_]/10. CI_wt_ values were calculated for mutually exclusive interactions where CI_wt_ <1 indicates synergism, CI_wt_ =1 indicates additive effects, and CI_wt_ ˃1 suggest antagonism.

### Quantification of SARS-CoV-2 RNA

Quantification of SARS-CoV-2 RNA was performed as previously described [Toptan et al., 2020]. SARS-CoV-2 RNA from cell culture supernatant samples was isolated using AVL buffer and the QIAamp Viral RNA Kit (Qiagen) according to the manufacturer’s instructions. Intracellular RNAs were isolated using the RNeasy Mini Kit (Qiagen) as described by the manufacturer. RNA was subjected to OneStep qRT-PCR analysis using the Luna Universal One-Step RT-qPCR Kit (New England Biolabs) or Luna Universal Probe One-Step RT-qPCR Kit (New England Biolabs) or LightCycler® Multiplex RNA Virus Master (Roche) using the CFX96 Real-Time System, C1000 Touch Thermal Cycler. The primer pairs for the E-, S- and M-gene-specific PCRs were used in equimolar concentrations (0.4 µM each per reaction). The RdRP primer pairs were used according to Corman et al. [Corman et al., 2020] with 0.6 µM and 0.8 µM concentrations of the forward and reverse primers, respectively.

The cycling conditions were used according to the manufacturer’s instructions. Briefly, for SYBR green-and probe-based Luna Universal One-Step RT-qPCR Kits, 2 µL of RNA was subjected to a reverse transcription reaction in a reaction volume of 20 µL, performed at 55 °C for 10 min. Initial denaturation was performed for 1 min at 95 °C, followed by 45 cycles of denaturation for 10 s and extension for 30 s at 60 °C. Melt curve analysis (SYBR green) was performed from 65–95 °C with an increment of 0.5 °C each 5 s. For the IVD-approved LightCycler® Multiplex RNA Virus Master (Roche), 5 µL of template RNA in a total reaction volume of 20 µL was used. Reverse transcription was performed at 55 °C for 10 min. Initial denaturation was induced for 30 s at 95 °C, followed by 45 cycles of denaturation for 5 s at 95 °C, and extension for 30 s at 60 °C, and a final cool-down to 40 °C for 30 s. The PCR runs were analysed with the Bio-Rad CFX Manager software, version 3.1 (Bio Rad Laboratories).

### Cell viability

Cell viability was determined using the CellTiter-Glo^®^ Luminescent Cell Viability Assay (Promega) according to the manufacturer’s instructions.

### Statistics

The results are expressed as the mean ± standard deviation (SD) of the number of biological replicates indicated in figure legends. The statistical significance is depicted directly in graphs and the statistical test used for calculation of p values is indicated in figure legends. GraphPad Prism 9 was used to determine IC50 values.

## Results

### Sequence differences in the interferon antagonists between Delta, Omicron BA.1, and Omicron BA.2

We have previously shown that Omicron BA.1 virus isolates display higher sensitivity to interferons than a Delta isolate [Bojkova et al., 2022b]. Sequence differences in a range of putative viral interferon antagonists may be responsible for this [Bojkova et al., 2022b].

Here, a comparison of sequence variants in a Delta, an Omicron BA.1, and an Omicron BA.2 virus isolate identified 96 sequence variants in putative interferon antagonists that differed from the reference genome of the original Wuhan strain (Suppl. Table 1).

Only three sequence variants were shared between all three isolates (Figure 1A). The overlap in sequence variants between BA.1 and BA.2 was larger (49) than between Delta and BA.1 (21) and Delta and BA.2 (18). Moreover, Delta displayed more unique sequence variants (54) in the putative interferon antagonists than BA.1 (23) or BA.2 (26) (Figure 1A).

**Figure 1.**
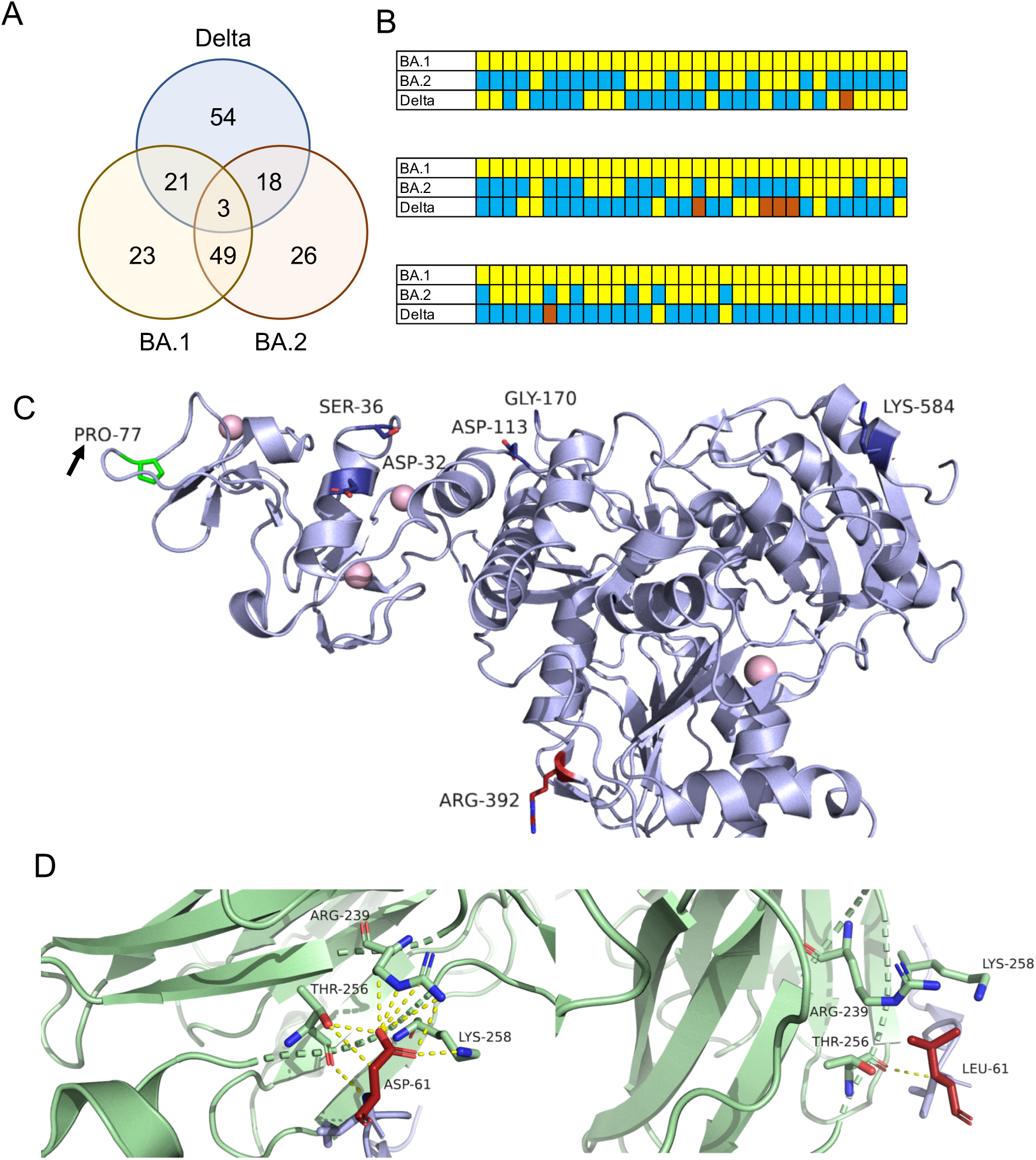
Sequence variants that may contribute to differences in the response to interferon treatment Omicron BA.1, Omicron BA.2, and Delta isolates. A) Overlaps between sequence variants in putative SARS-CoV-2 interferon antagonists determined in BA.1 (FFM-SIM0550/2021, GenBank ID: OL800702), BA.2 (FFM-BA.2-3833, GenBank ID: OM617939), and Delta (FFM-IND8424/2021, GenBank ID: MZ315141) isolates. B) A heatmap illustrating the differences in amino acid residues in SARS-CoV-2 proteins anticipated to be of potential relevance for interferon signalling between the SARS-CoV-2 isolates (differences in colour indicate different residues). C) Key residues of NSP13 thought to antagonise interferon signalling via interaction with TBK1. Only the Delta isolate is harbouring a P77L (proline/ Pro to leucine/ Leu) change (highlighted by an arrow), which has been proposed to affect the interaction of NSP13 and TBK1 [Rashid et al., 2021]. Source PDB structure 7re2. D) ORF6 antagonises the cellular interferon response by direct interaction of its C-terminal domain with the RNA binding pocket of the Nup98-Rae1 complex [Miorin et al., 2020; Kato et al., 2021]. While BA.1 and Delta harbour leucine (L/ Leu) in position 61, BA.2 harbours an aspartate (D/ Asp) in this position. The number of hydrogen bonds between ORF6 and Rae1 is anticipated to be strongly reduced when aspartate (left image) is replaced by leucine (right image). The resulting reduced complex stability is likely to modify the capacity of ORF6 to suppress the cellular interferon response. Source PDB structure 7vph.

These findings appear to reflect the closer relatedness of BA.1 and BA.2 relative to Delta. However, the variant overlaps are complex (Figure 1B), and it is not clear, which of them drive the virus response to interferons.

45 of the 96 sequence variants could be modelled on protein structures or models (Suppl. File 1). However, it was difficult to draw reliable conclusions. Many of the mutations were in the Spike (S) protein, for which detailed information on its role as interferon antagonist is lacking (Suppl. Table 1, Suppl. File 1).

There are only two sequence variants that are likely to modify interferon signalling. NSP13 inhibits the activation of an interferon response by physically interacting with the TANK binding kinase 1 (TBK1), which, prevents the phosphorylation, dimerisation, and nuclear translocation of the interferon regulatory factors 3 and 7 (IRF3, IRF7). In contrast to BA.1 and BA.2, the Delta isolate harbours a P77L change in NSP13 (Figure 1C, Suppl. Table 1, Suppl. File 1). This proline (P) to leucine (L) change is likely to have an impact, as a proline at this position has been proposed to be crucial for the NSP13-TBK1 interaction [Rashid et al., 2021].

Moreover, BA.2 harbours in contrast to BA.1 and Delta a D61L sequence variant in ORF6 (Figure 1C, Suppl. Table 1, Suppl. File 1), which has been shown to antagonise the cellular interferon response by inhibiting the nuclear translocation of STAT1 and STAT2 through direct interaction of its C-terminal domain with the RNA binding pocket of the Nup98-Rae1 complex [Miorin et al., 2020; Kato et al., 2021].

This aspartic acid (D) to leucine (L) change at position 61 in the C-terminal domain has the potential to be significant, as the number of hydrogen bonds formed between ORF6 and Rae1 is predicted to be strongly reduced when the aspartate is replaced by a leucine (Figure 1D).

Taken together, sequence differences between the SARS-CoV-2 interferon antagonists in BA.1, BA.2, and Delta warrant the further comparison of these three SARS-CoV-2 variants for their responses to interferon treatment.

### Replication kinetics of Delta, Omicron BA.1, and Omicron BA.2 in Caco-2 cells

Previously, we have shown that a Delta isolate infects a higher proportion of cells and replicates to higher titres in Caco-2-F03 cells (a Caco-2 subline that is highly susceptible to SARS-CoV-2 infection [Bojova et al., 2022b]) than two Omicron BA.1 isolates [Bojkova et al., 2022; Bojkova et al., 2022a]. Here, these results were confirmed (Figure 2A-C). BA.2 (GenBank ID OM617939) replicated more effectively than BA.1 but less effectively than Delta in Caco-2 cells (Figure 2-C).

**Figure 2.**
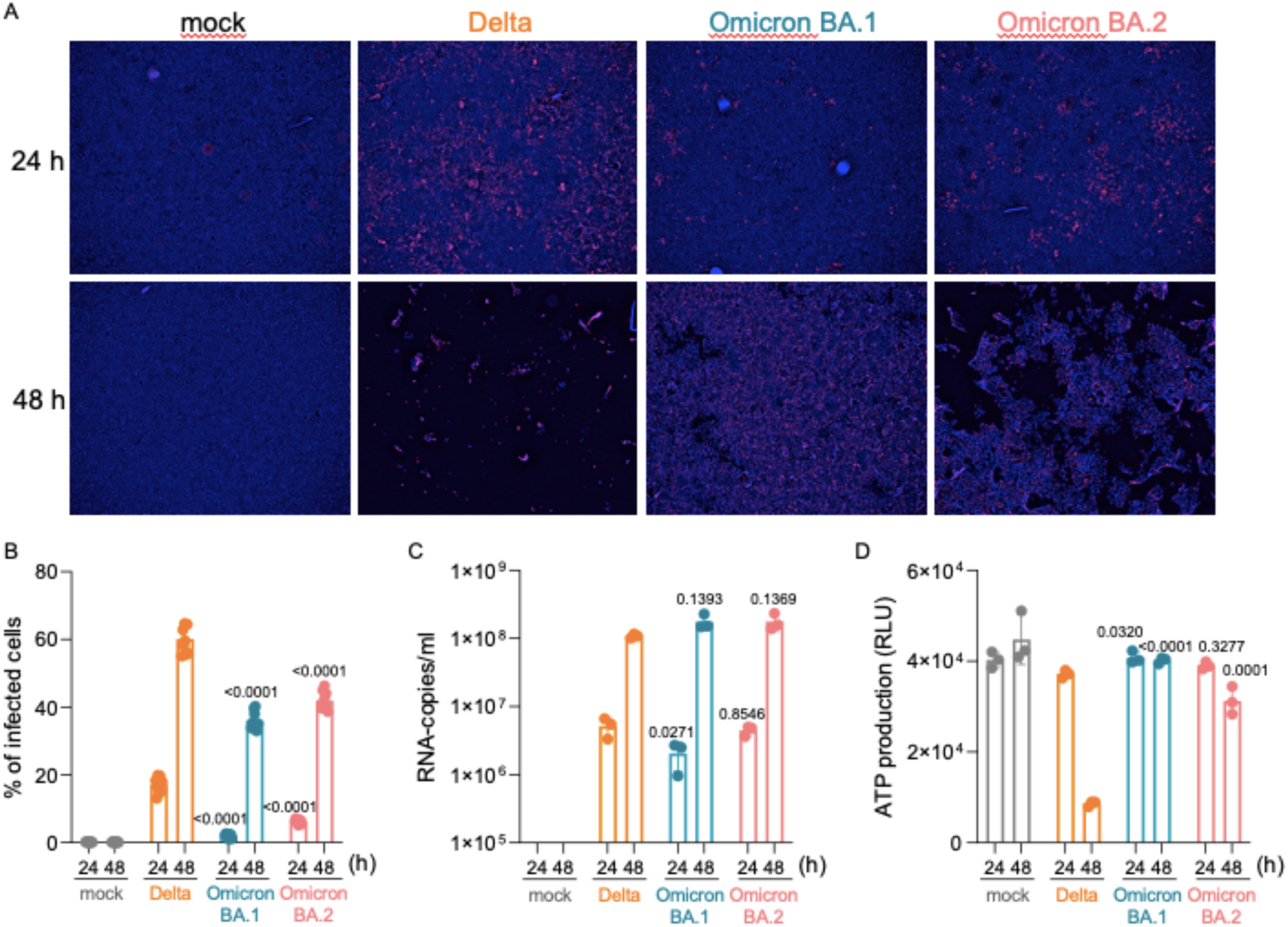
Replication kinetics of SARS-CoV-2 Delta, Omicron BA.1, and Omicron BA.2 isolates in Caco-2 cells. A) Representative immunofluorescence images indicating the number of Spike (S) protein-positive Caco-2-F03 cells 24h and 48h post infection with Delta, BA.1, and BA.2 at an MOI of 1. B) Quantification of S protein-positive Caco-2-F03 cells 24h and 48h post infection with Delta, BA.1, and BA.2 at an MOI of 1. C) Genomic RNA copy numbers determined by qPCR 24h and 48h post infection of Caco-2 cells with Delta, BA.1, and BA.2 at an MOI of 1. D) Cell viability in Caco-2-F03 cells 24h and 48h post infection as determined by CellTiter-Glo^®^ Luminescent Cell Viability Assay (Promega). Values represent mean ± S.D. of three independent experiments. P-values represent statistical differences between Delta and BA.1 or BA.2 calculated by one-way ANOVA and Tukey’s test. These differences in the replication kinetics (Delta > BA.2 > BA.1) were also reflected in cytopathogenic effect (CPE) formation (Figure 2A) and cell viability measurements (Figure 2D).

### Delta, Omicron BA.1, and BA.2 sensitivity to approved antiviral drugs

Next, we tested the effects of the approved anti-SARS-CoV-2 drugs remdesivir, EIDD-1931 (the active metabolite of the prodrug molnupiravir), and nirmatrelvir (the antivirally active agent in Paxlovid) on Delta, BA.1, and BA.2 replication. All three isolates displayed similar sensitivity to all three drugs (Figure 3A, 3B).

**Figure 3.**
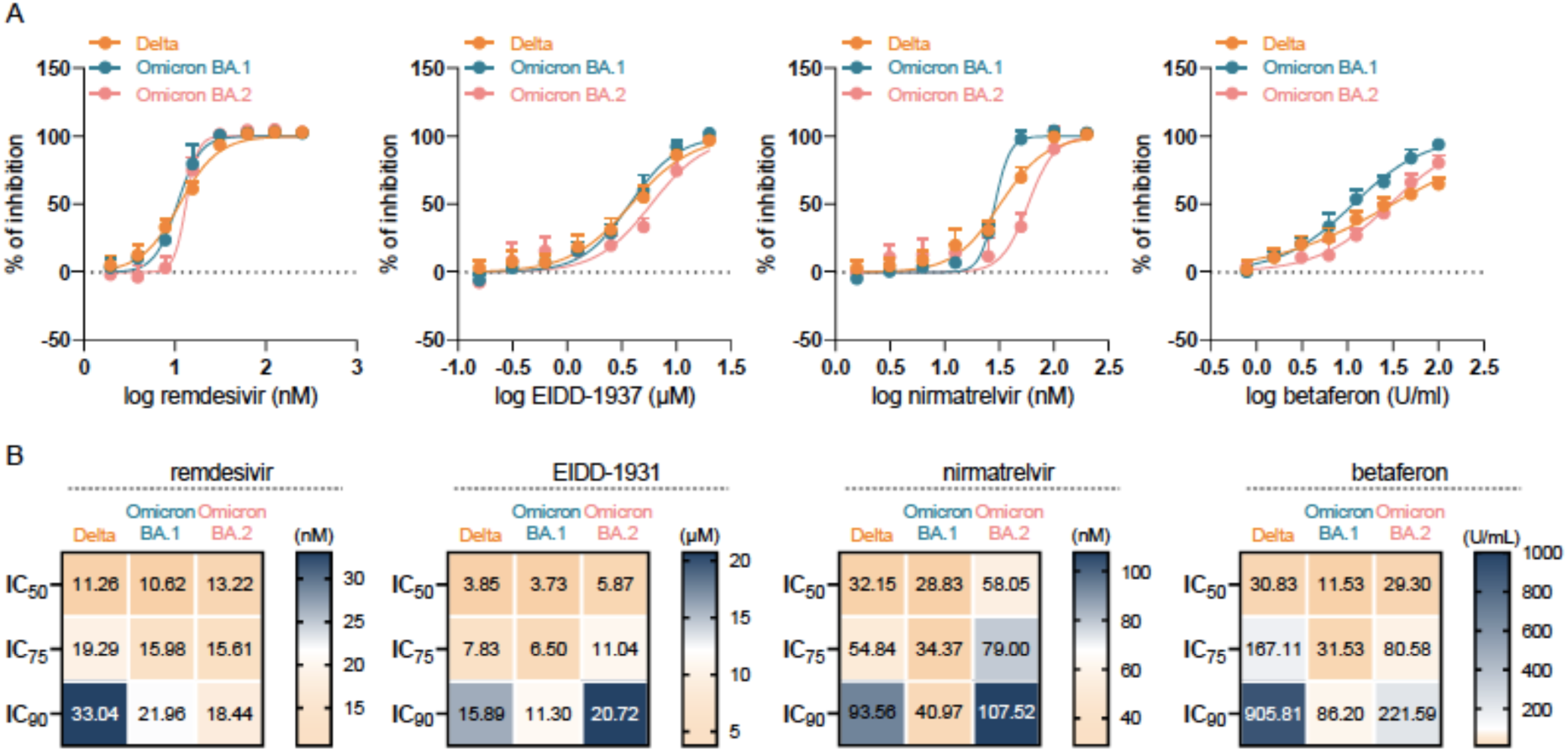
Delta, Omicron BA.1, and Omicron BA.2 isolate sensitivity to antiviral drugs. A) Dose response curves and B) concentrations that reduce the number of Spike (S)-protein positive cells by 50% (IC_50_), 75% (IC_75_), and 90% (IC_90_) in Caco-2-F03 cells infected with the different SARS-CoV-2 isolates at an MOI 1 24 h post infection, as determined by immunostaining.

We had previously shown that BA.1 is more sensitive to interferon-β than Delta [Bojkova et al., 2022b]. This time, we used the clinically approved interferon-β preparation betaferon (Bayer) for our experiments. In agreement with the previous data, betaferon was more effective against BA.1 than against Delta (Figure 3A, 3B). Interestingly and perhaps unexpectedly, the betaferon response of BA.2 more closely resembled that of Delta and not that of the more closely related BA.1 (Figure 3A, 3B). This confirmed our previous findings (Figure 1) that the impact of amino acid sequence differences in different SARS-CoV-2 isolates on the viral interferon response is not easily predictable and can differ even between closely related virus variants.

### Effects of betaferon in combination with approved anti-SARS-CoV-2 drugs

Since our previous findings had shown that interferon-β displayed different levels of synergism with remdesivir, EIDD-1931, and nirmatrelvir [Bojkova et al., 2022a], we further tested betaferon in combination with these drugs.

Results were comparable to the previous findings [Bojkova et al., 2022b]. Remdesivir displayed additive to moderately synergistic effects in combination with betaferon against all three variants (Figure 4). While EIDD-1931 and nirmatrelvir treatment resulted in similar levels of synergism with betaferon against Delta, combined EIDD-1931 and interferon treatment was associated with a more pronounced synergism against BA.1 and BA.2 than the combination of nirmatrelvir and betaferon (Figure 4).

**Figure 4.**
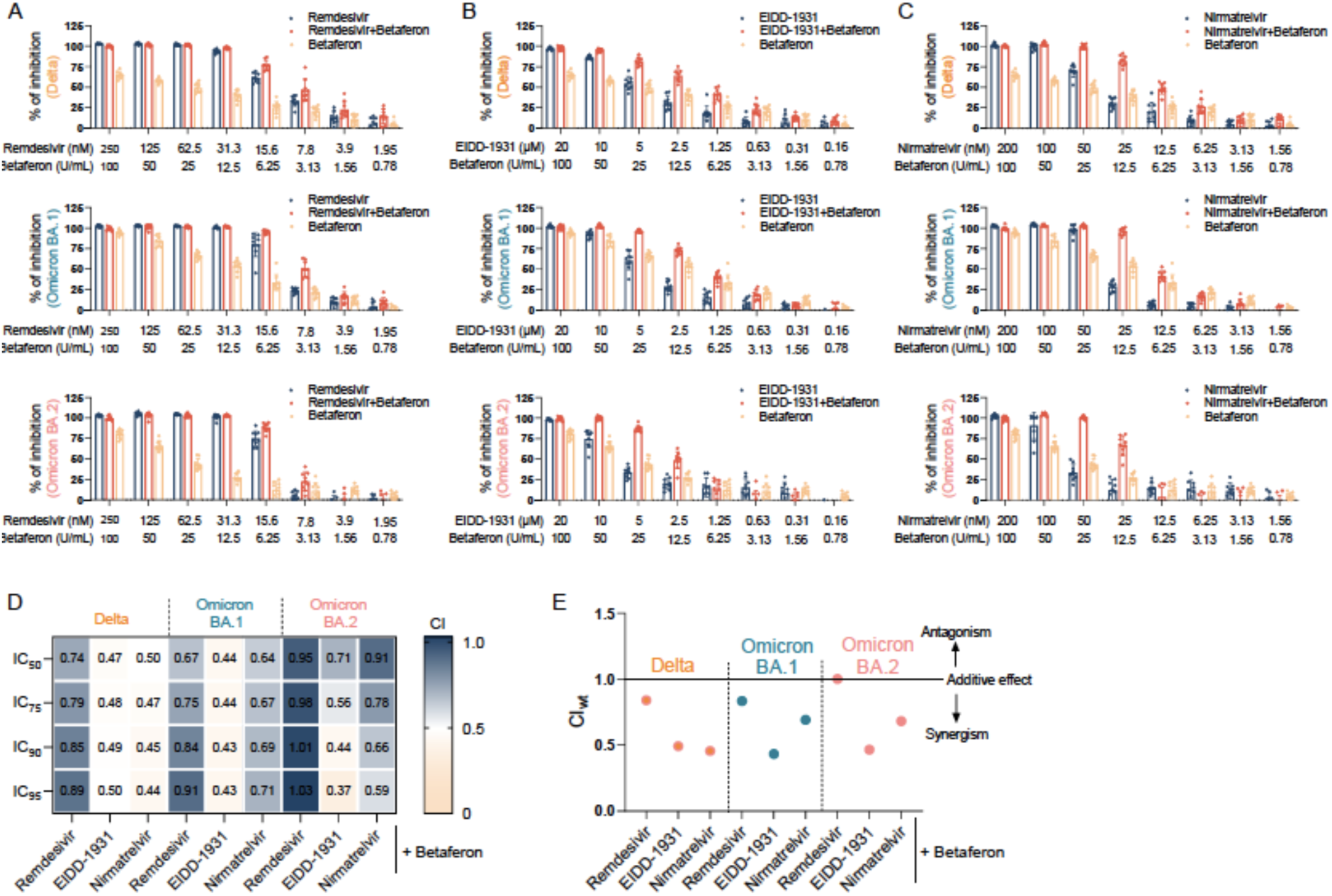
Antiviral effects of approved anti-SARS-CoV-2 drugs in combination with interferon-β (betaferon) against Delta, Omicron BA.1, and Omicron BA.2 isolates. Betaferon was tested in fixed combinations combination with remdesivir (A), EIDD-1931 (B), or nirmatrelvir (C) in SARS-CoV-2 (MOI 0.01)-infected Caco-2-F03 cells. Values represent mean ± S.D. of three independent experiments. D) Combination indices were calculated at the IC_50_, IC_75_, IC_90_, and IC_95_ levels following the method of Chou and Talalay [Chou, 2006]. E) The weighted average CI value (CI_wt_) was calculated according to the formula: CI_wt_ [CI_50_ + 2CI_75_ + 3CI_90_ + 4CI_95_]/10. A CI_wt_ <1 indicates synergism, a CI_wt_ =1 indicates additive effects, and a CIwt ˃1 suggest antagonism.

### Effects of betaferon in combination with the antiviral protease inhibitor aprotinin

Previously, we have shown that the protease inhibitor aprotinin inhibits replication of the SARS-CoV-2 original Wuhan strain at least in part by inhibition of the cleavage and activation of the viral spike (S) protein by host cell proteases [Bojkova et al., 2020]. Based on these findings, a clinical trial was initiated that reported improved outcomes of COVID-19 patients treated with an aprotinin aerosol [Redondo-Calvo et al., 2022]. Among other improvements, aprotinin treatment reduced the length of hospital stays by five days [Redondo-Calvo et al., 2022].

Here, we show that aprotinin inhibits Delta (IC50: 0.66µM) and BA.1 (IC50: 0.64µM) in a similar concentration range as the original Wuhan strain isolates [Bojkova et al., 2020] (Suppl. Figure 1). Effects against BA.2 were less pronounced (IC50: 1.95µM) but still in the range of clinically achievable plasma concentrations after systemic administration, which have been described to reach 11.8µM [Levy et al., 1994; Bojkova et al., 2020]. Moreover, aerosol preparations like the one used in the clinical trial that demonstrated therapeutic efficacy of aprotinin against COVID-19 [Redondo-Calvo et al., 2022] are expected to result in substantially higher aprotinin concentrations locally in the lungs.

Interestingly, aprotinin displayed a strong synergism with betaferon against Delta and an even much stronger synergism against BA.1 and BA.2 (Figure 5).

**Figure 5.**
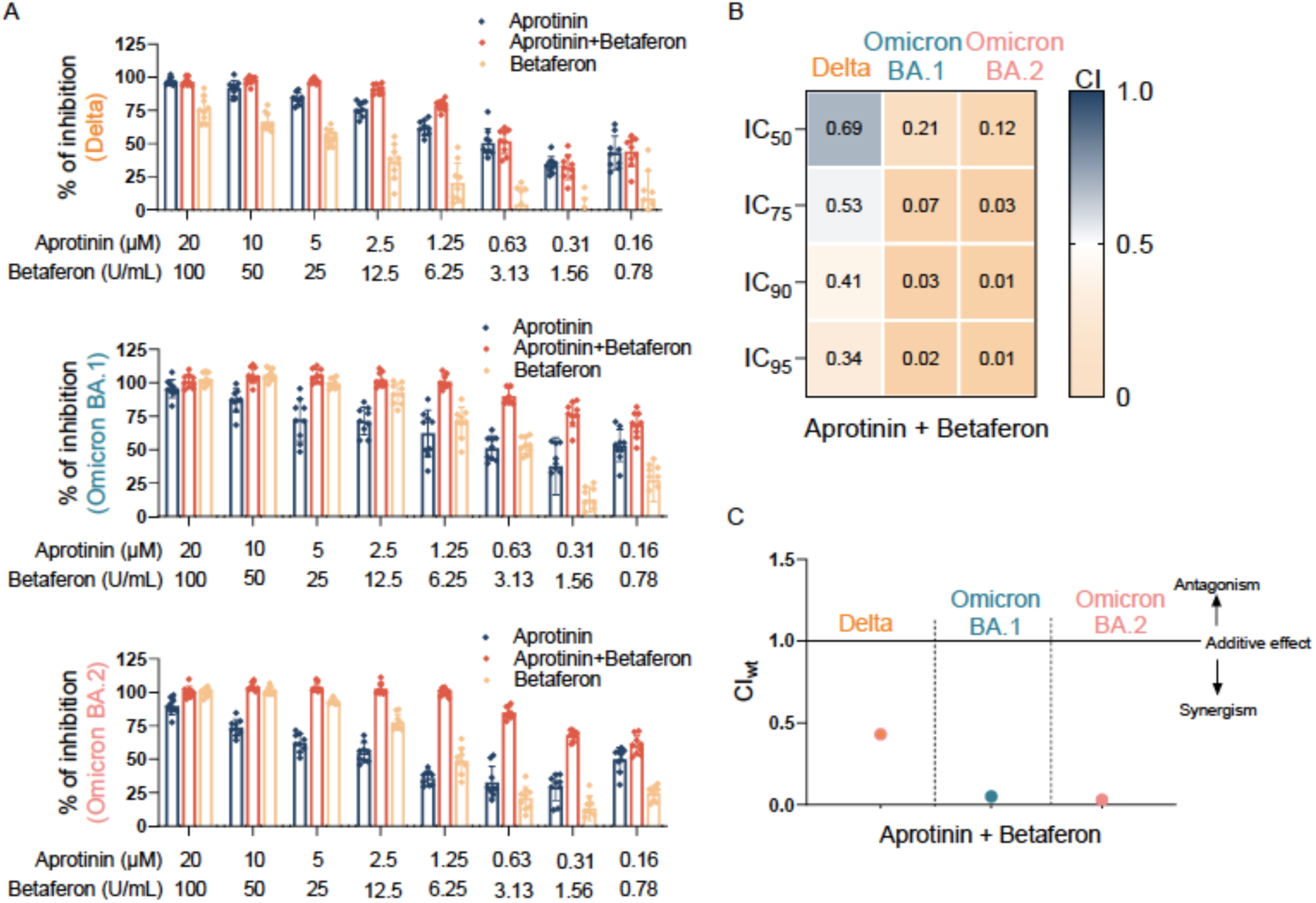
Antiviral effects of aprotinin in combination with interferon-β (betaferon) against Delta, Omicron BA.1, and Omicron BA.2 isolates. Betaferon was tested in a fixed combination with aprotinin in SARS-CoV-2 (MOI 0.01)-infected Caco-2-F03 cells. Values represent mean ± S.D. of three independent experiments. B) Combination indices were calculated at the IC_50_, IC_75_, IC_90_, and IC_95_ levels following the method of Chou and Talalay [Chou, 2006]. C) The weighted average CI value (CI_wt_) was calculated according to the formula: CI_wt_ [CI_50_ + 2CI_75_ + 3CI_90_ + 4CI_95_]/10. A CI_wt_ <1 indicates synergism, a CI_wt_ =1 indicates additive effects, and a CI_wt_ ˃1 suggest antagonism.

## Discussion

Previously, we found that Omicron BA.1 variant isolates induce a stronger interferon response and replicate less effectively in interferon-competent cells than a Delta isolate. Moreover, BA.1 isolates were more sensitive to interferon treatment than a Delta isolate [Bojkova et al., 2022; Bojkova et al., 2022a]. Here, we show that an Omicron BA.2 isolate displays intermediate replication patterns in interferon-competent Caco-2-F03 cells when compared to BA.1 and Delta isolates. Moreover, BA.2 is less sensitive than BA.1 and similarly sensitive as Delta to betaferon treatment.

The reasons for these differences are not obvious. The sequence differences in the putative viral interferon antagonists are complex. There are two sequence variants for which there is plausible evidence that they may impact on the viral interferon sensitivity based on an *in silico* structural analysis. The Delta isolate harbours a P77L change in NSP13 that is likely to have an impact on TBK1-mediated interferon signalling [Rashid et al., 2021]. Additionally, BA.2 harbours in contrast to BA.1 and Delta a D61L sequence variant in ORF6 that may modify the potential of ORF6 to antagonise the cellular interferon response [Miorin et al., 2020; Kato et al., 2021]. However, these changes are probably just small pieces in a large puzzle of virus protein interactions with the complex regulatory networks that determine the cellular interferon response [Blalock, 2021].

Delta and BA.2 displayed similar sensitivity to the approved anti-SARS-CoV-2 drugs remdesivir, nirmatrelvir, and EIDD-1931 (the active metabolite of molnupiravir), whereas BA.2 was less sensitive to EIDD-1931 than Delta and BA.1. Moreover, BA.2 was less sensitive than BA.1 and Delta to aprotinin, a protease inhibitor that was previously shown to inhibit the original SARS-CoV-2 Wuhan strain and demonstrated clinical efficacy in COVID-19 patients [Bojkova et al., 2020; Redondo-Calvo et al., 2022].

When we investigated these four drugs in combination with betaferon, only betaferon combinations with nirmatrelvir, EIDD-1931, and aprotinin resulted in synergistic activity. We also detected variant-specific differences. While nirmatrelvir and EIDD-1931 showed similar synergy with betaferon against Delta, the betaferon/ EIDD-1931 synergism was more pronounced than the betaferon/ nirmatrelvir synergism against BA.1 and BA.2. Aprotinin displayed the strongest synergism with betaferon against BA.1 and BA.2 among all tested drugs. Against Delta, the level of synergism of aprotinin/ betaferon was similar to that of EIDD-1931/ betaferon. Given the differences between the SARS-CoV-2 (sub)variants, our data suggest that an improved understanding of the combined effects of antiviral drugs on certain SARS-CoV-2 variants can inform the design of optimised combination therapies.

Effective antiviral combination therapies are anticipated to be of crucial importance for the control of virus outbreaks, as a higher efficacy is expected to decrease or even prevent resistance formation [White et al., 2021]. So far, clinical studies reported mixed outcomes in patients treated with remdesivir/ interferon combinations [Kalil et al., 2021; Tam et al., 2022]. This may not be too surprising in the light of our current findings, suggesting that combining betaferon with nirmatrelvir, molnupiravir, and aprotinin is more promising than with remdesivir.

In conclusion, even closely related SARS-CoV-2 (sub)variants can differ in their biology, as indicated by different BA.1 and BA.2 replication kinetics, and in their response to antiviral treatments, as indicated by differences in the virus responses to betaferon, EIDD-1931/ molnupiravir, and aprotinin and differing levels of synergism of betaferon combinations with other antiviral drugs. Betaferon combinations with nirmatrelvir and, in particular, with EIDD-1931 and aprotinin displayed high levels of synergism, which makes them strong candidates for clinical testing.

## Acknowledgements

We thank Lena Stegman, Kerstin Euler, and Sebastian Grothe for their technical assistance.

## Suppl. File 1

**Suppl. Figure 1.**
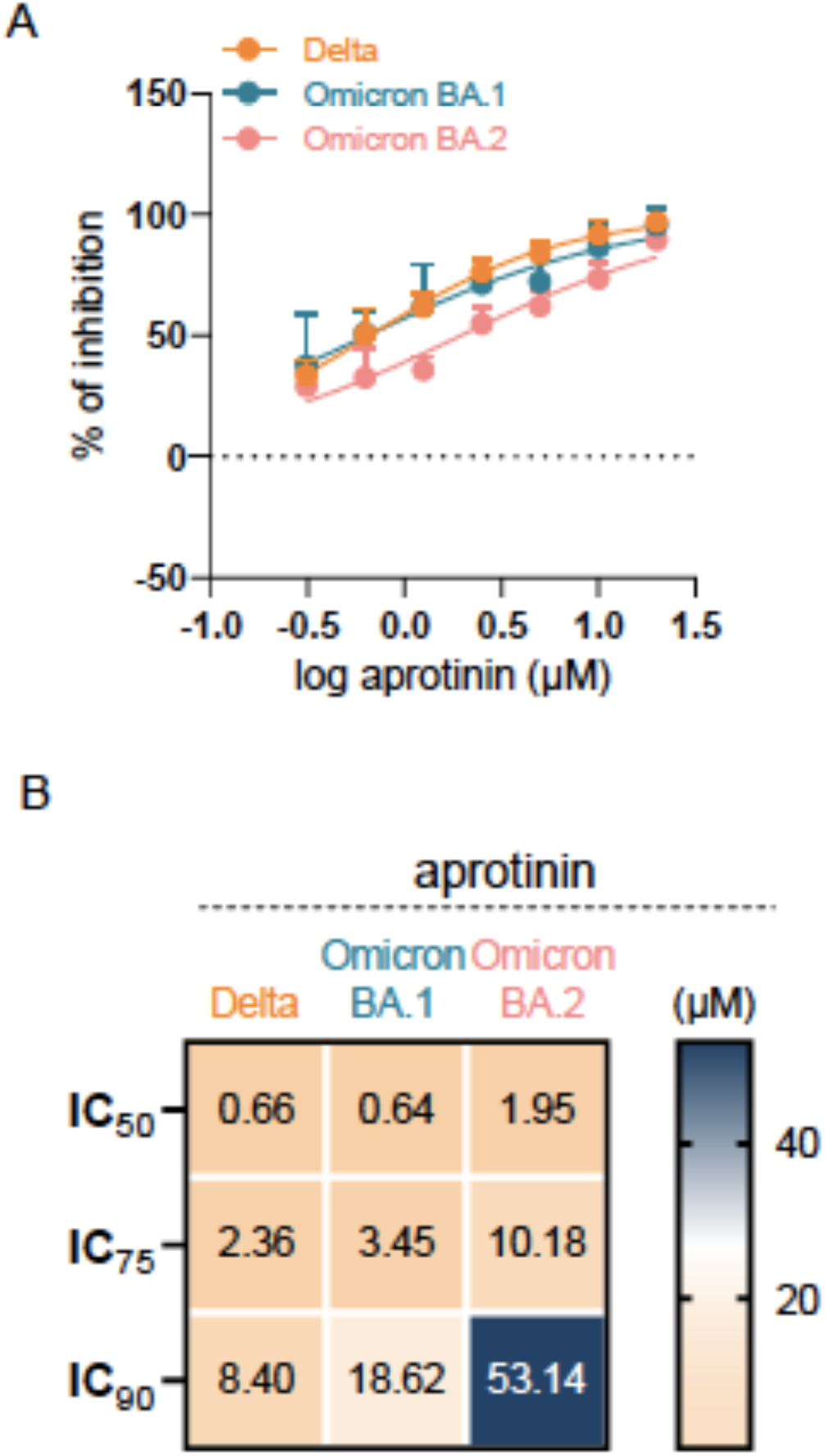
Anti-SARS-CoV-2 effects of aprotinin against SARS-CoV-2 isolates. A) Dose response curves and B) concentrations that reduce the number of Spike (S)-protein positive cells by 50% (IC_50_), 75% (IC_75_), and 90% (IC_90_) in Caco-2 cells infected with the different SARS-CoV-2 isolates at an MOI 1 24 h post infection, as determined by immunostaining.

**Suppl. Table 1.**
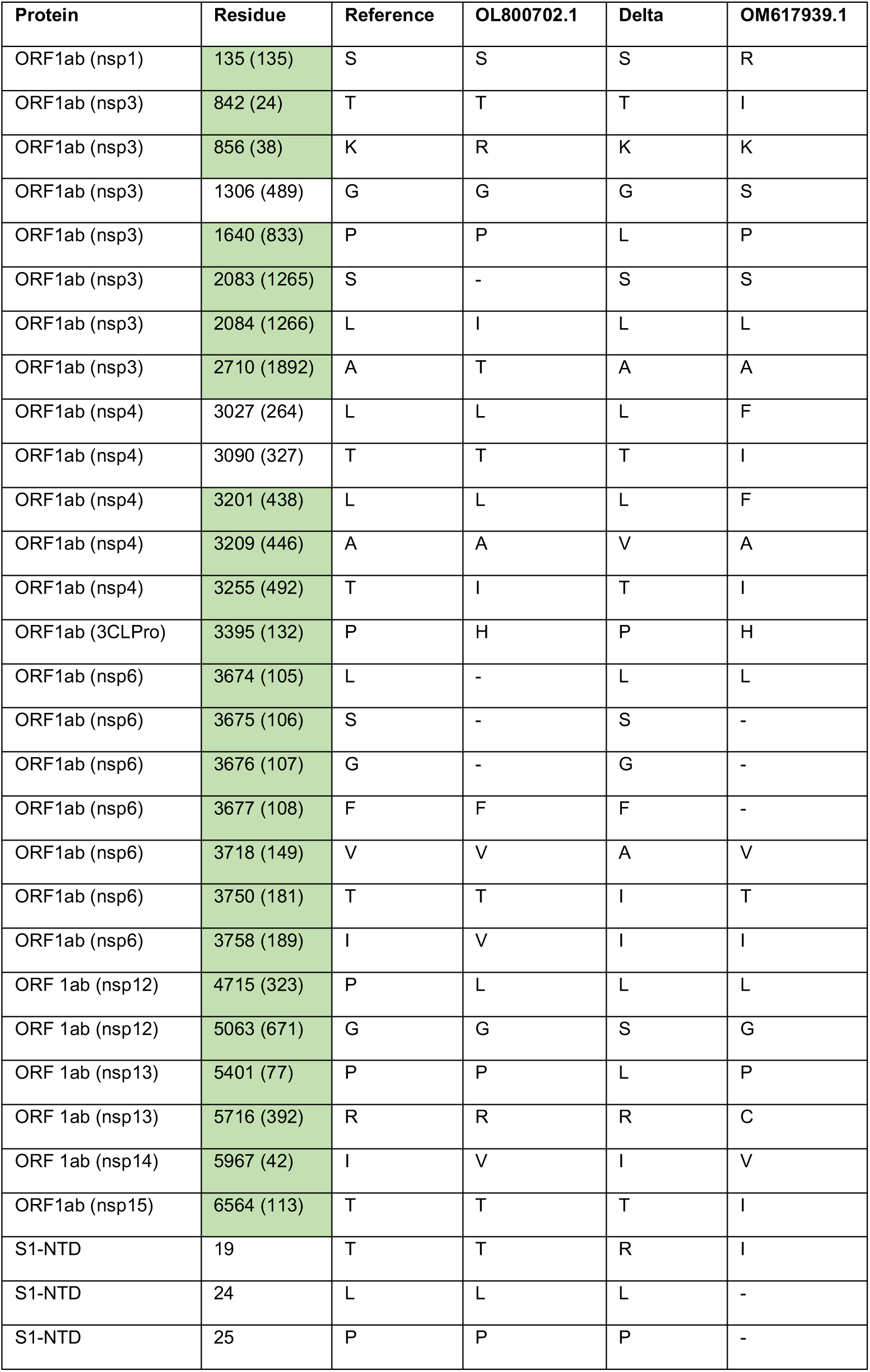
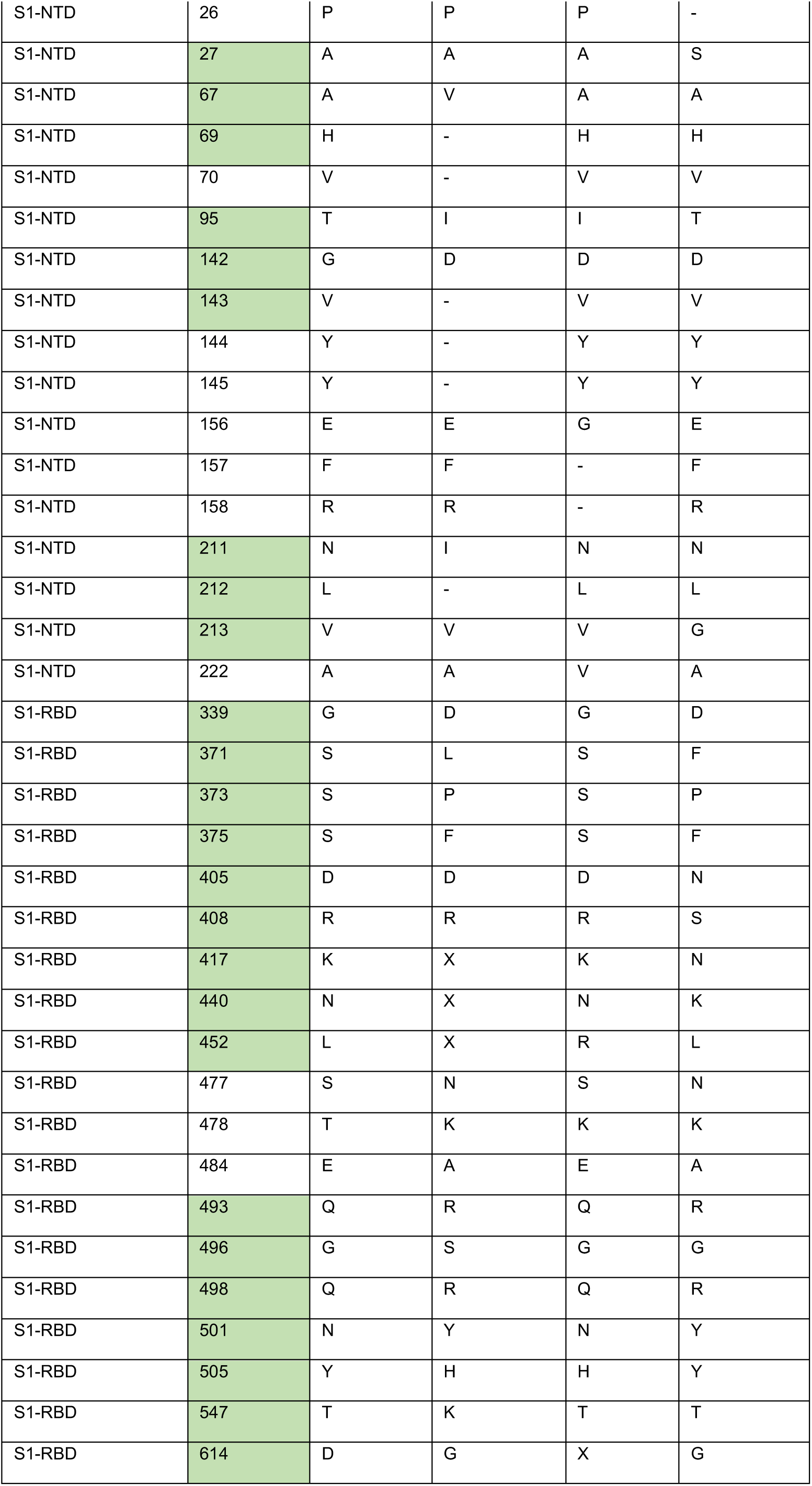
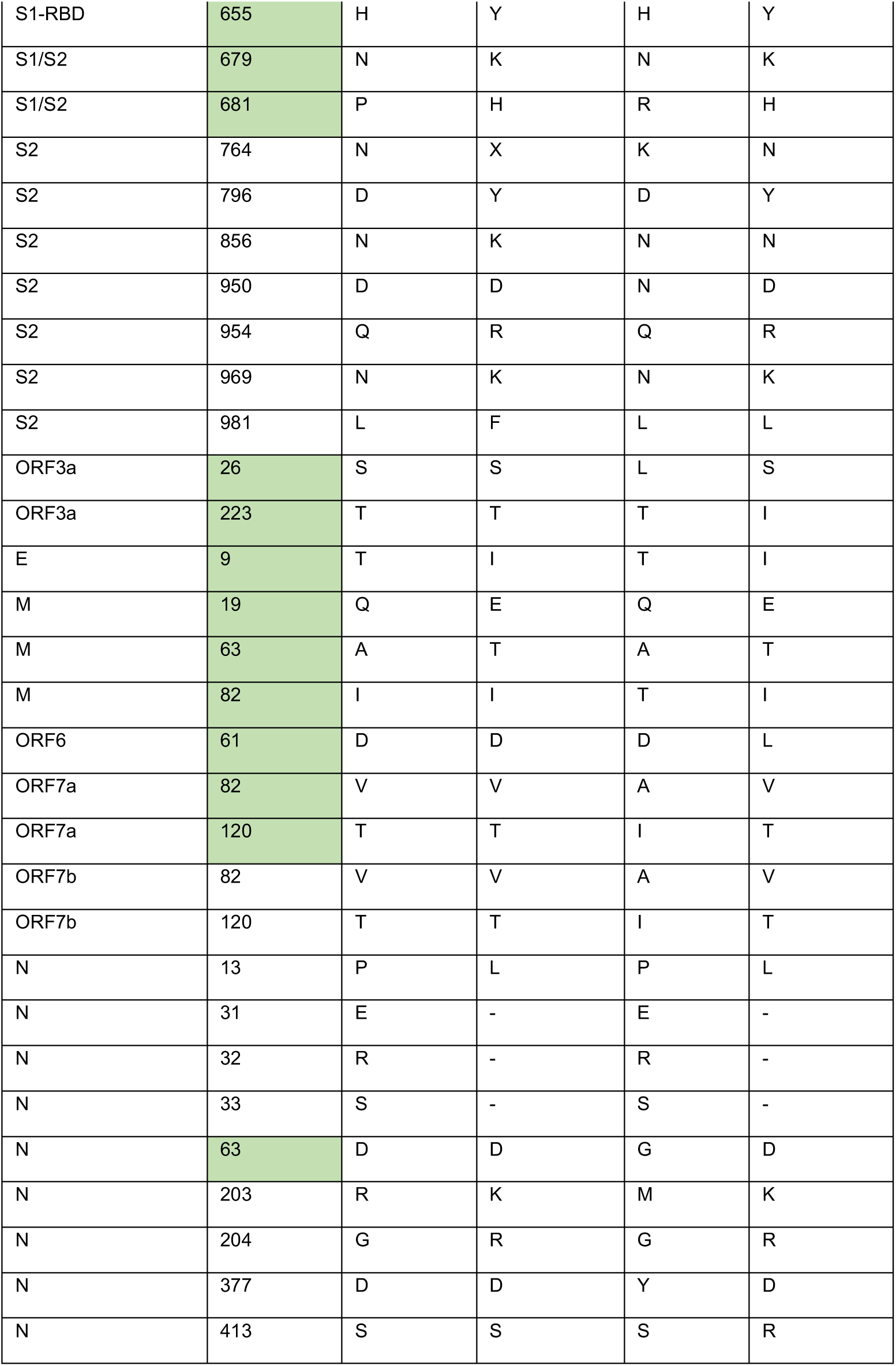
SARS-CoV-2 Omicron-associated sequence variants in proteins described to possess interferon-antagonising activity [Abbas et al., 2022]. Residue numbers are based on the Wuhan reference sequence with residue numbers in brackets indicating the position in the individual proteins within ORF1ab. The residue present in the reference sequence is shown in the reference column. – indicates a deletion.

## Suppl. File 1. Notes on the sequence variants from Suppl. Table 1 that could be modelled on protein structures and models

### ORF1ab polyprotein

*Nsp1.*

Model shown is from COVID-3D as no PDB structures exist that contain the residue of interest.

BA.2 mutation S135R. Serine to Arginine.

Blosum score of -1. Polar uncharged to positively charged.

ConSurf score of 1. Highly variable residue.

ΔΔG^stability^ mCSM: -0.58 kcal.mol^-1^ (destabilising).

This mutation is present in 98.1% of all BA.2 sequences.

Nsp1 has a globular N-terminal domain, a short linking domain, and a C-terminal helix-turn-helix motif. Its C-terminal binds to the mRNA entry channel of 40S ribosomal subunits, blocking host translation including interferons and ISGs (1).

Ser-135 is located in the linking domain and does not form bonds with residues within the C-terminal domain. There is one polar bond connecting Ser-135 to the globular domain which is unaltered upon mutation, and it does not appear to be providing stability to the C-terminal domain. Given the high variability of this residue it is likely that the mutation is inconsequential.

**Fig 1:**
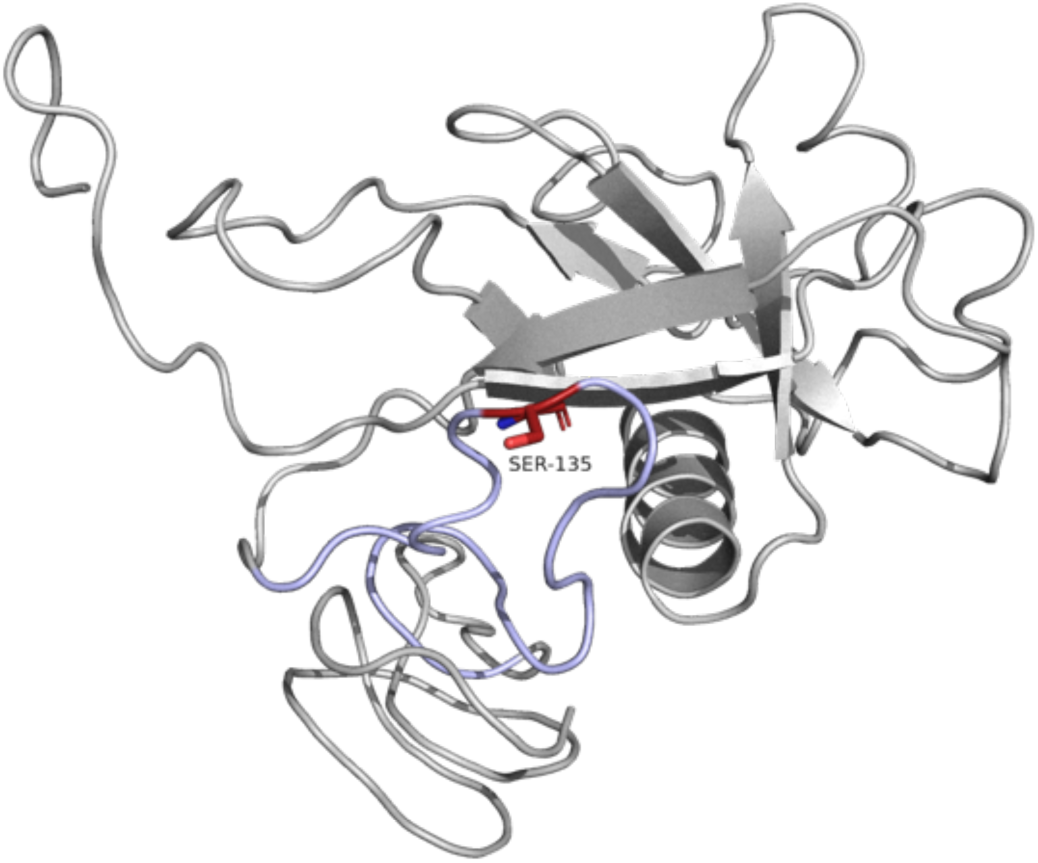
Cartoon of nsp1. The N-terminal globular domain is in the upper part of the image. Linking domain is highlighted in light blue, Ser-135 in red. The C-terminal domain is in the lower part of the image. Source structure 3-D Covid (2).

*Nsp3: PL-pro.*

PDB structure 6xa9.

Delta Mutation. P77L. Proline to Leucine.

Blosum score of -3. Special case to hydrophobic.

ConSurf score of 3. Moderately variable residue.

ΔΔG^stability^ mCSM: -0.24 kcal/mol^-1^ (Destabilising).

This mutation is only present in 9.1% of all Delta sequences.

PLpro is a domain of nsp3 with protease activity. This PDB structure has assembled PLpro as a homotrimer, but it is shown below as a monomer in association with ISG15. Pro-77 is located at the end of an alpha helix, on a turn. It is likely to be providing rigidity to this part of the structure. mCSM predicts this is a destabilising mutation, and that Pro-77 loses a polar and a hydrogen bond to surrounding residues on mutation. However, Pro-77 is located some distance from two binding areas, described as fingers and thumb (mainly composed of sections of beta-sheets).

**Figure 2.**
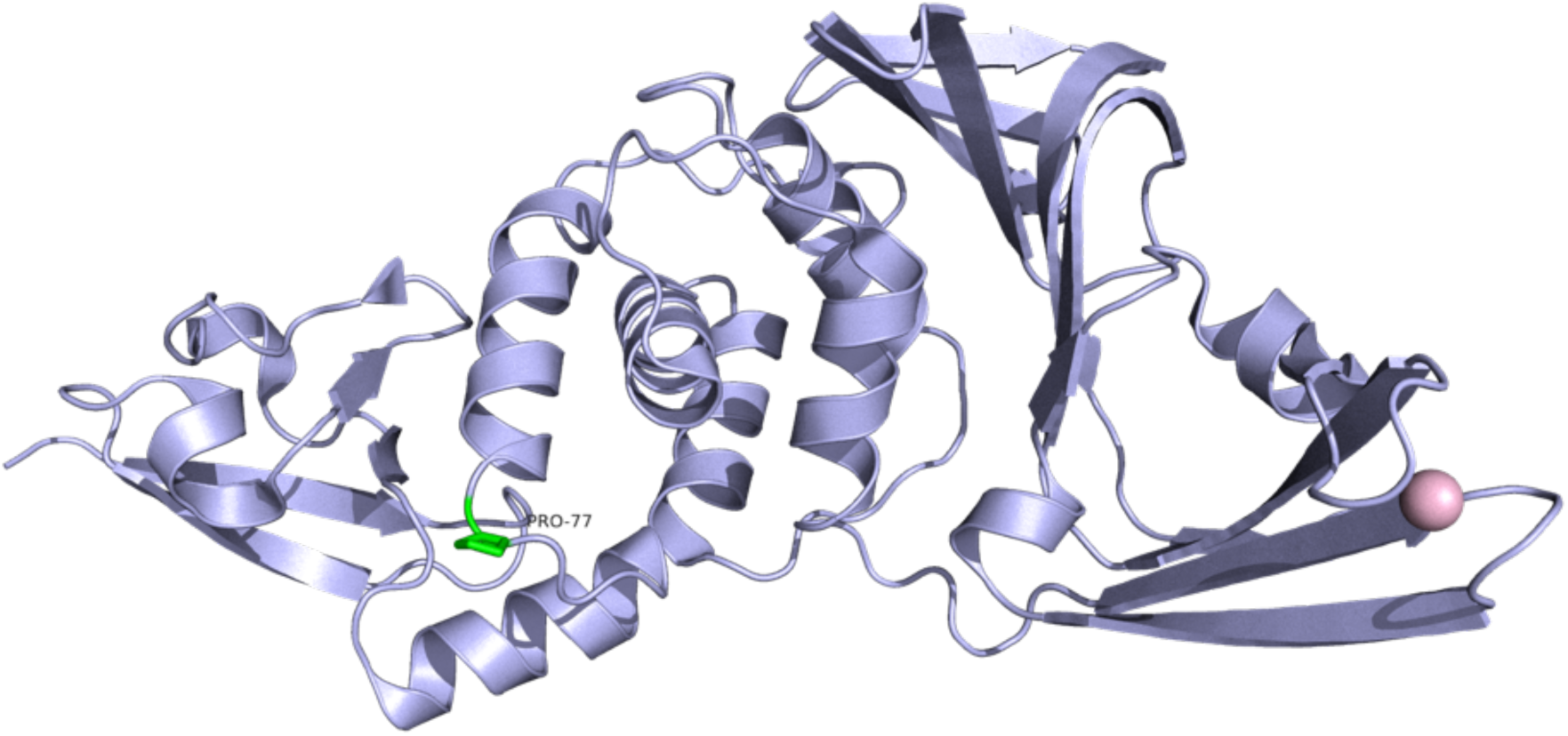
Cartoon representation of PLpro, monomeric form. Pro-77 is highlighted in green. Source PDB structure 7cjm.

PLpro has been shown to cleave ISG15, and its enzymatic activity appears to be essential to its inhibitory activity against Type I IFN responses (3). Two structures showing alternative bindings of PLpro with ISG15 are shown below. mCSM-PPI2 predicted no change in affinity between the two proteins on mutation. However, since mCSM predicted a decrease in structural stability, there could be an implication on binding with ISG15, either increasing or decreasing affinity between the two proteins.

**Figure 3.**
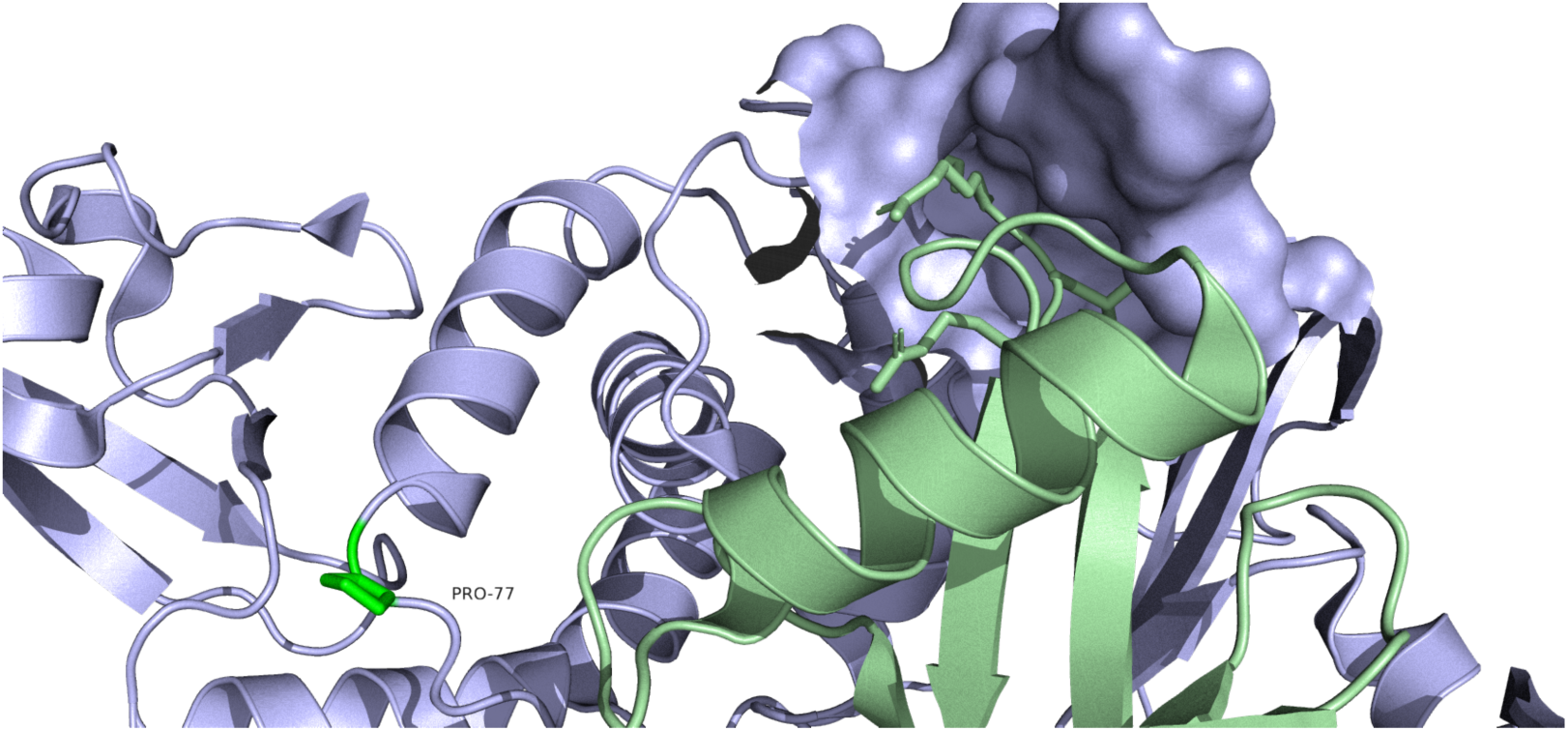
PLpro in association with ISG15. ISG15 (light green chain) associates with the thumb area of PLpro, which is the same region where small molecule inhibitors associate. In the image above, the binding pocket has been rendered with a surface to show distance from Pro-77 (highlighted in green). Source PDB structure 6xa9.

**Figure 4.**
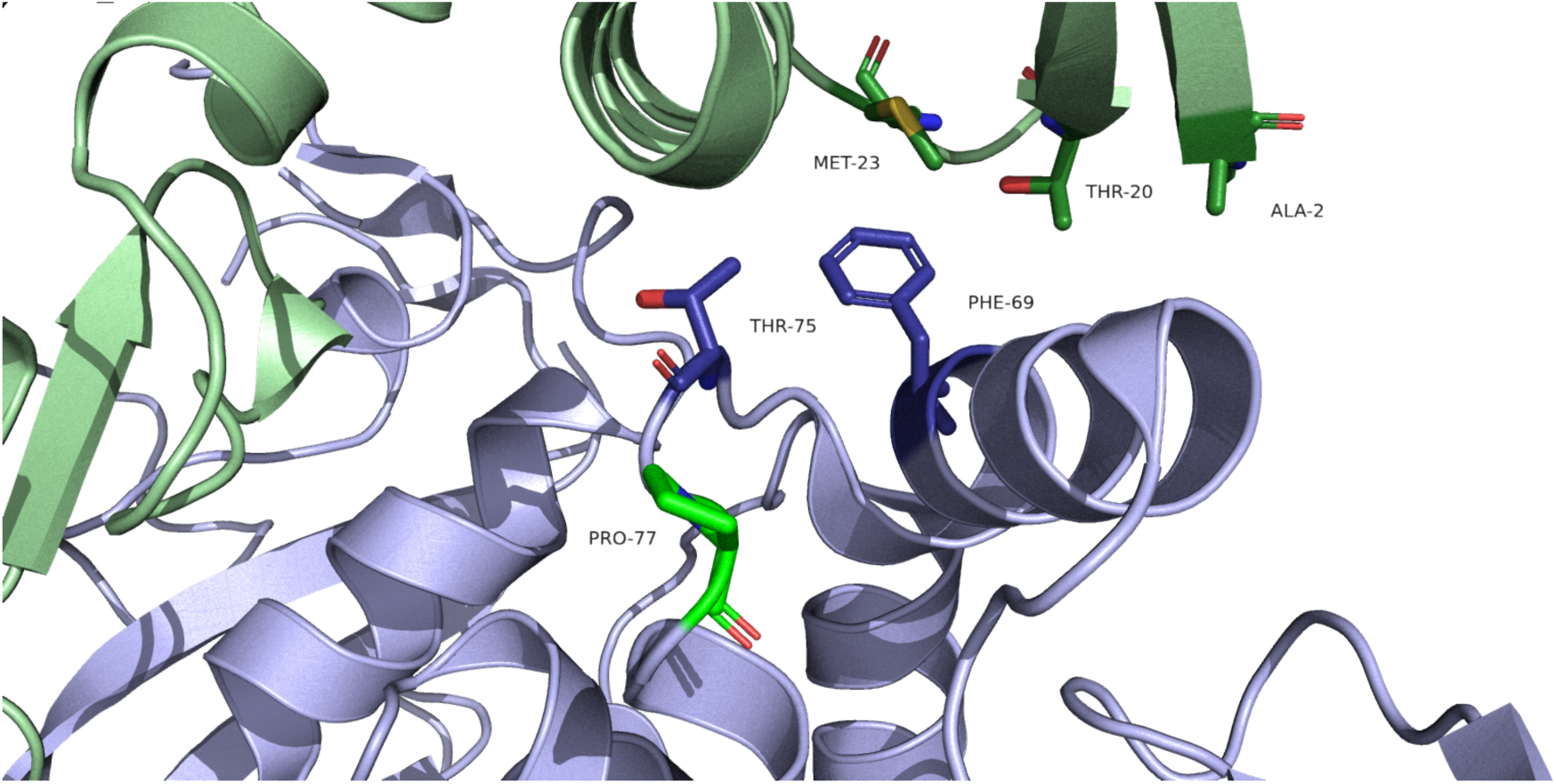
PLpro in association with ISG15, alternative binding. Residues Ala-2, Thr-20 and Met-23 of ISG15 (light green) form a hydrophobic patch, which mediates the association with PLpro. The key residues on PL-pro, where mutation to Alanine decreases binding to ISG15 and enzymatic activity are Thr-75 and Phe-69 (deep blue) . Pro-77 (bright green) is positioned a short distance along the loop from these residues, away from the hydrophobic patch. Source PDB structure 6yva.

*Nsp5: 3CLpro.*

PDB structure 7lmc.

BA.1/BA.2 Mutation. P132H. Proline to Histidine.

Blosum score of -2. Special case to positively charged.

ConSurf score of 3. Moderately variable residue.

ΔΔG^stability^ mCSM: -1.77 kcal/mol^-1^ (Destabilising).

This mutation is present in 98.1% of all BA.1 sequences and 99.4% of BA.2 sequences.

Numerous model structures exist with multiple ligands - mostly inhibitors. There is consensus that dimerisation of 3CLpro is essential to stabilise the conformation of the catalytic site.

Pro-132 is situated on a turn, and is located away from binding/catalytic sites (Fig 5-8), and opposite to the plane where the protomers associate with one another (Fig 5). Blosum score is low, but perhaps because both residues contain a cyclic compound, the effect of removing a proline from this loop position might be lessened. Modelling mutagenesis in Pymol shows that all 9 rotamers of Histidine create multiple clashes with surrounding residues, which may be why mCSM predicts this as a destabilising mutation.

The enzymatic (cysteine protease) activity of 3CLpro was described to be essential for the inhibition of interferon induction (4). Both RIG-I and NEMO are potential cleavage targets of 3CLpro (4,5). Since the protease activity of 3CLpro is so important to viral replication, it seems unlikely that this mutation has much significance.

**Figure 5.**
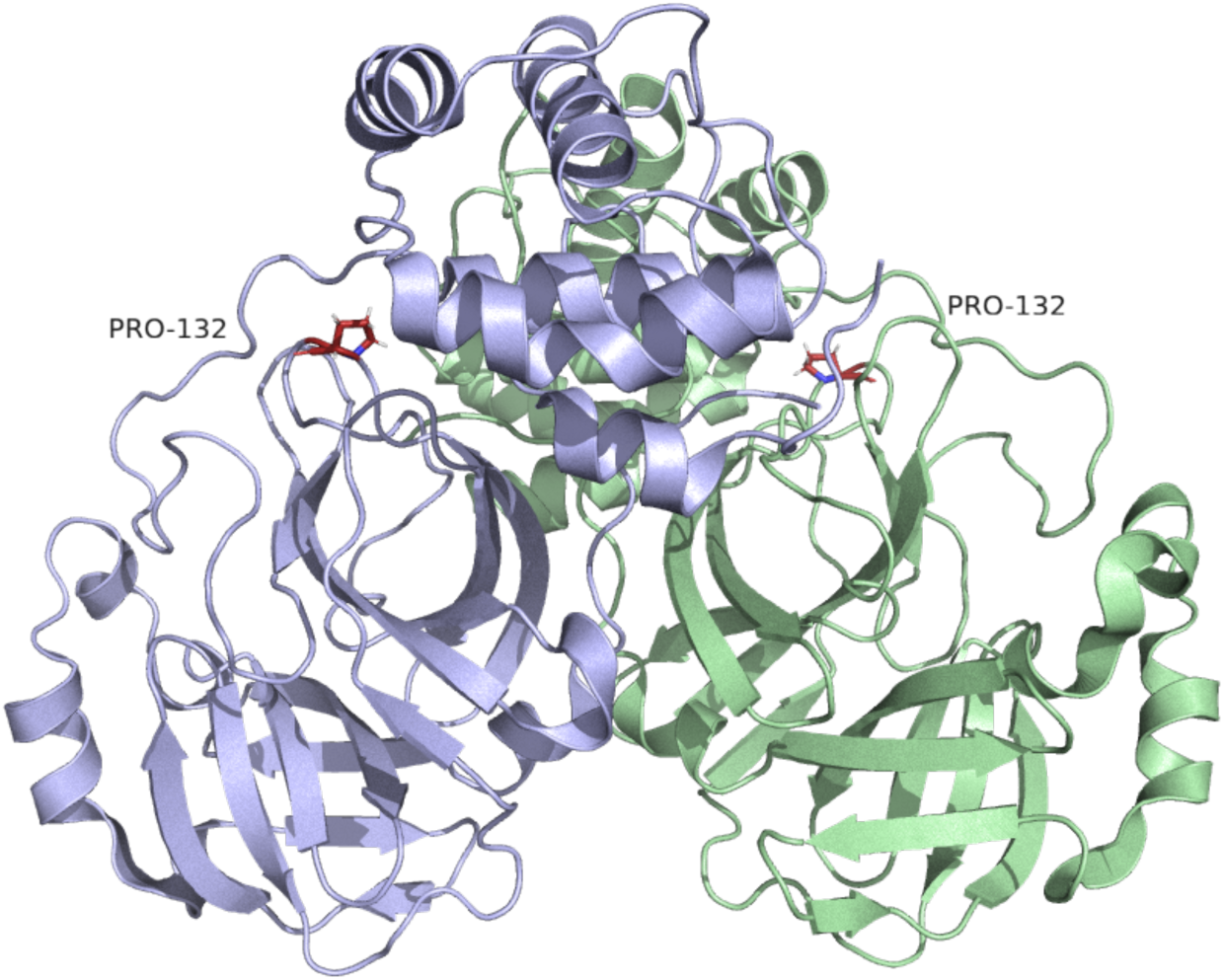
3CLpro assembled as a homodimer. Cartoon representation of 3CLpro. Protomers are coloured light blue and light green. Pro-132 is highlighted in red. Source PDB structure 7bb2.

**Figure 6.**
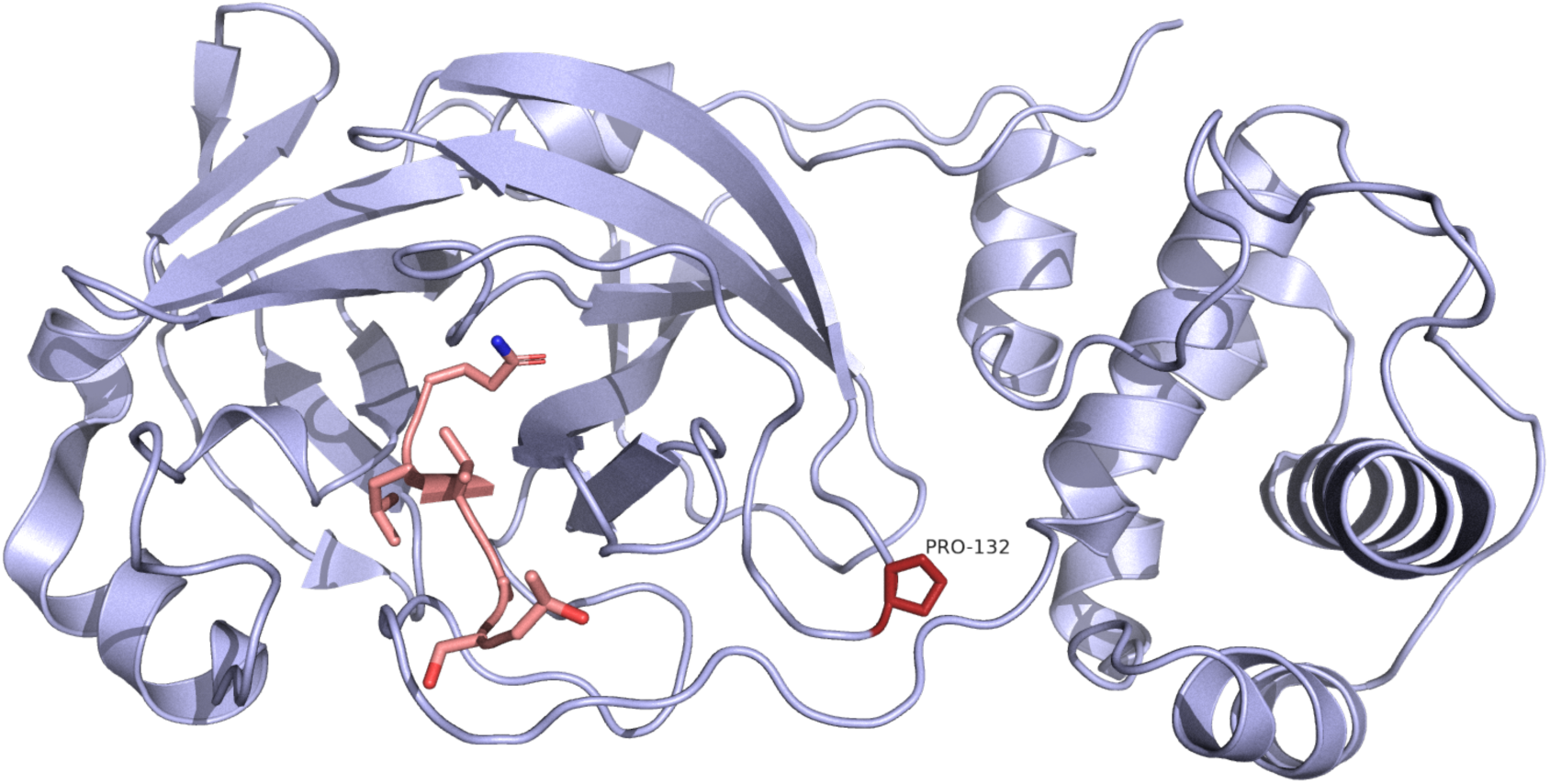
3CLpro in association with the C-terminal of nsp4. Cartoon representation of 3CLpro in monomeric form. The C-terminal of nsp4 is highlighted in pink, whilst Pro-132 is highlighted in red. The binding site for nsp4 appears to also be the catalytic site of 3CLpro. Pro-132 is some distance from this site. Source PDB structure 7lmc.

**Figure 7.**
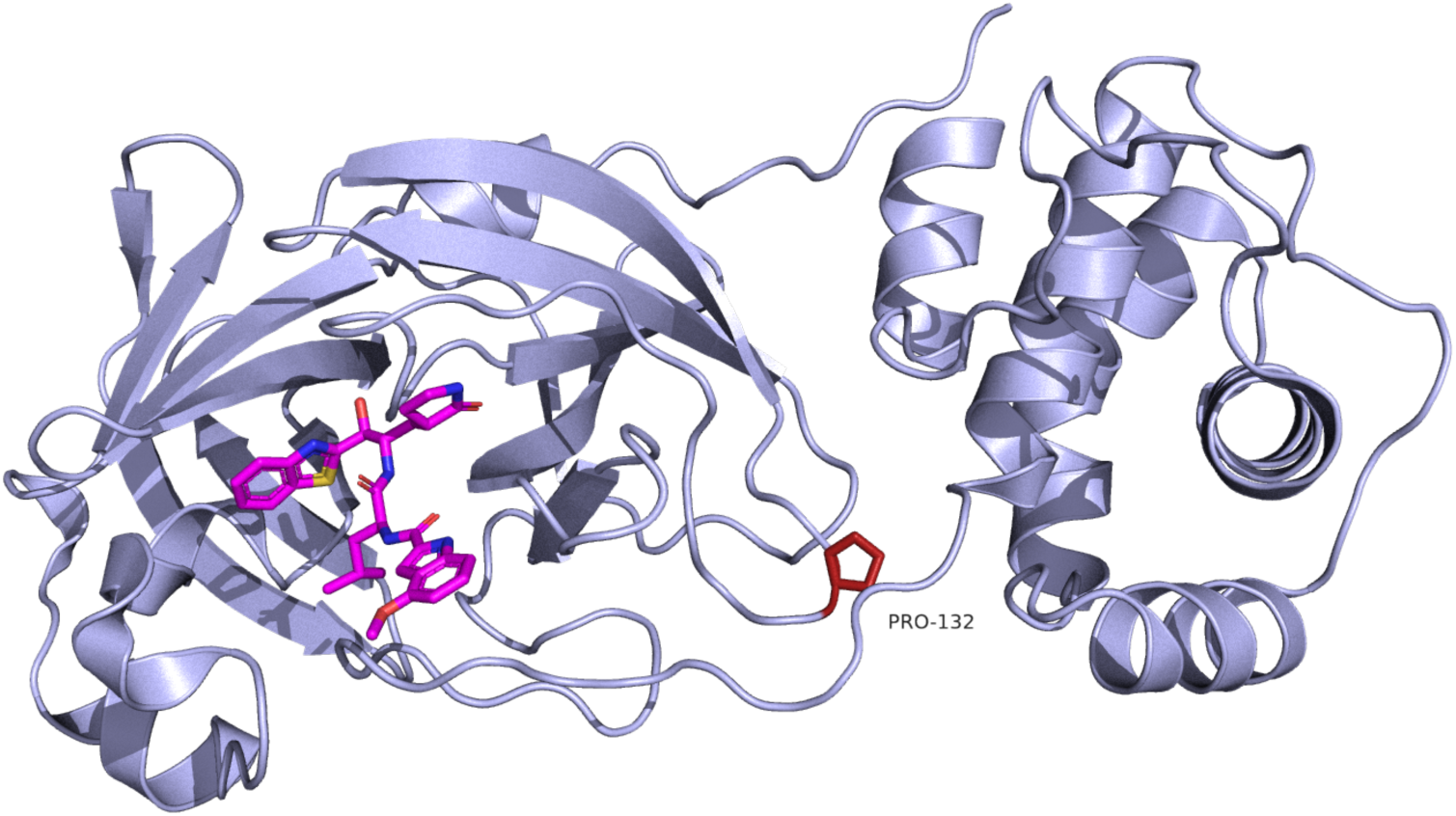
3CLpro in association with small molecule inhibitor GRL2420. Cartoon representation of 3CLpro. The small molecule inhibitor GRL2420 is highlighted in magenta. This binding pocket appears constant for multiple small molecule inhibitors and the C-terminal of nsp4. This area is associated with conserved residues across all coronaviruses, essential to its enzymatic activity (6). Source PDB structure 7jkv.

**Figure 8.**
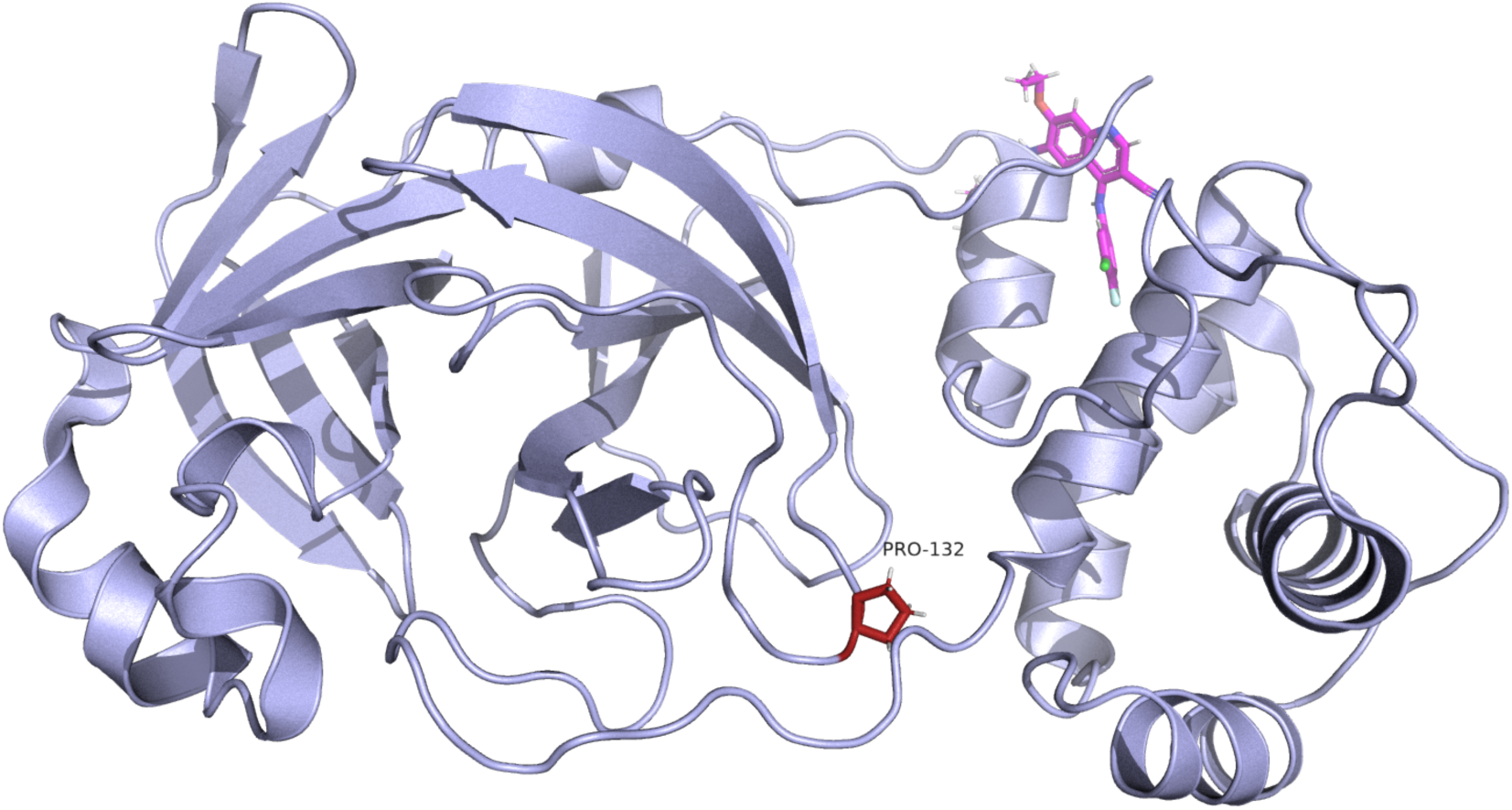
3CLpro in association with Pelitinib, which binds to the allosteric dimerisation domain. Cartoon representation of 3CLpro. Pelitinib is highlighted in pink. This small molecule inhibitor binds to a hydrophobic patch thought to be a dimerisation domain (7). This site appears to be distant from Pro-132. Source PDB structure 7axm.

*Nsp12: RNA-dependent RNA polymerase.*

PDB structure 7b3b.

BA.1/BA.2/Delta Mutation. P323L. Proline to Leucine.

Blosum score of -3. Special case to hydrophobic.

ConSurf score of 1. Highly variable residue.

ΔΔG^stability^ mCSM: -0.29 kcal/mol^-1^ (Destabilising).

This mutation is present in 99.0% of all Delta, 98.7% of all BA.1 and 99.4% of all BA.2 sequences.

Delta Mutation. G671S: Glycine to Serine.

Blosum score of 0. Special case to polar uncharged.

Consurf score of 8. Highly conserved residue.

ΔΔG^stability^ mCSM: -0.97 kcal/mol^-1^ (Destabilising).

This mutation is present in 97.5% of all Delta sequences.

This structure is often modelled with nsp7 and nsp8, and on occasion with nsp13. Here, monomeric form is shown with RNA and remdesivir bound to show active sites.

Nsp12 has a Nidovirus RdRp-associated nucleotidyltransferase (NiRNA) domain (1-249), and a right hand RdRp domain (365-932), connected by an interface domain (249-365).

The catalytic residues in the RdRp domain are SDD (759-761), and the GDD motif (823-825) which is catalytic in other viral RdRps (8).

Importantly, there are mixed reports on nsp12’s IFN antagonistic activity.

It has been reported that nsp12 inhibits nuclear translocation of IRF3, and that this is not dependent on either the RdRp enzymatic activity or the NiRNA domain, nor is the inhibitory activity modulated by nsp12’s association with both nsp7 and nsp8 (8). However, a subsequent study reports that their own observations of IFN inhibitory activity was a result of HA-tags on viral proteins in a luciferase assay (9).

Pro-323 is located on a bend in a loop between an alpha helix and the start of a beta sheet in the interface domain. This change results in a reduction in residue size, and the residue is located near the surface of the structure, where nsp8 is closely associated (Fig 10). The change may have potential to affect the association of nsp8 with nsp12 by affecting local structure, but when taking residue variability into account it seems that the protein is able to accommodate changes at this point and there are no direct bonds from this residue. Moreover, this sequence variant is shared between all three isolates and, thus, unlikely to account for differences between them.

Gly-671 is located in the RdRp domain, but does not form part of the catalytic site. The replacement of gly-671 by serine does not appear to affect polar connections in Pymol, although prediction in 3D-Covid increases polar connections from 3 to 5 to residues in adjacent loops. Gly-671 is located on the surface of the protein, but some distance from the binding sites of nsp 7, nsp8 and the RNA binding groove (Fig 11).

**Figure 9.**
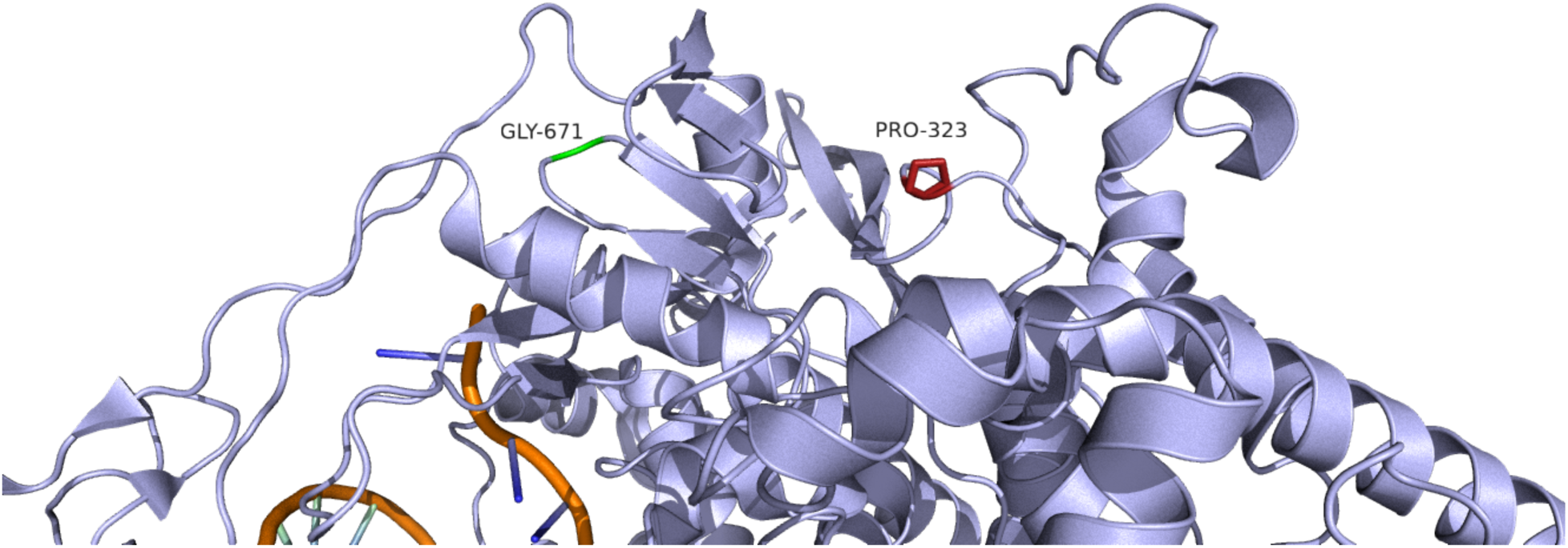
Close view of Nsp12. Cartoon representation of nsp12, showing position of residues of interest in relation to RNA. Gly-671 is highlighted in green, whilst Pro- 323 is highlighted in Red. Source PDB structure 7b3b.

**Figure 10.**
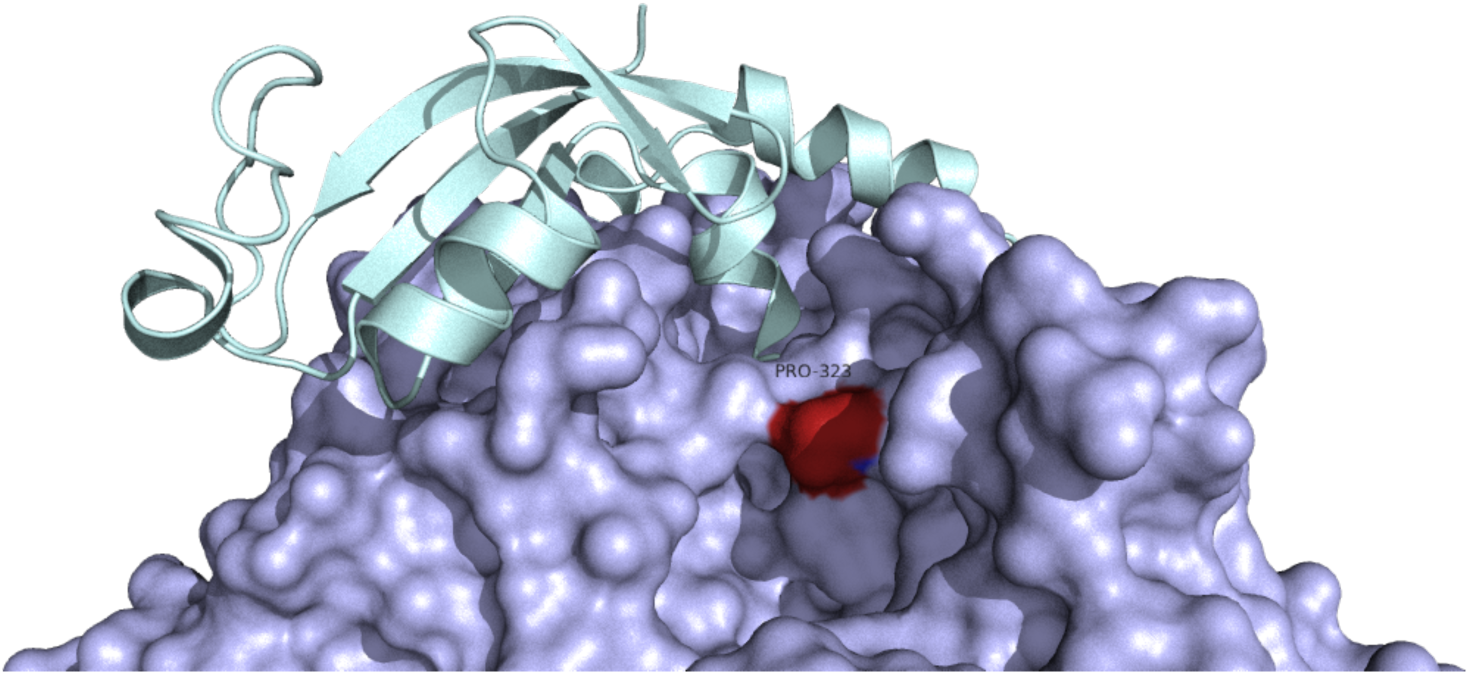
Nsp12 in association with Nsp8. Surface representation of Nsp12. The position of Pro-323 is highlighted in red. Cartoon representation of Nsp 8 is shown in close association, coloured in light green. Source PDB structure 7b3b.

**Figure 11.**
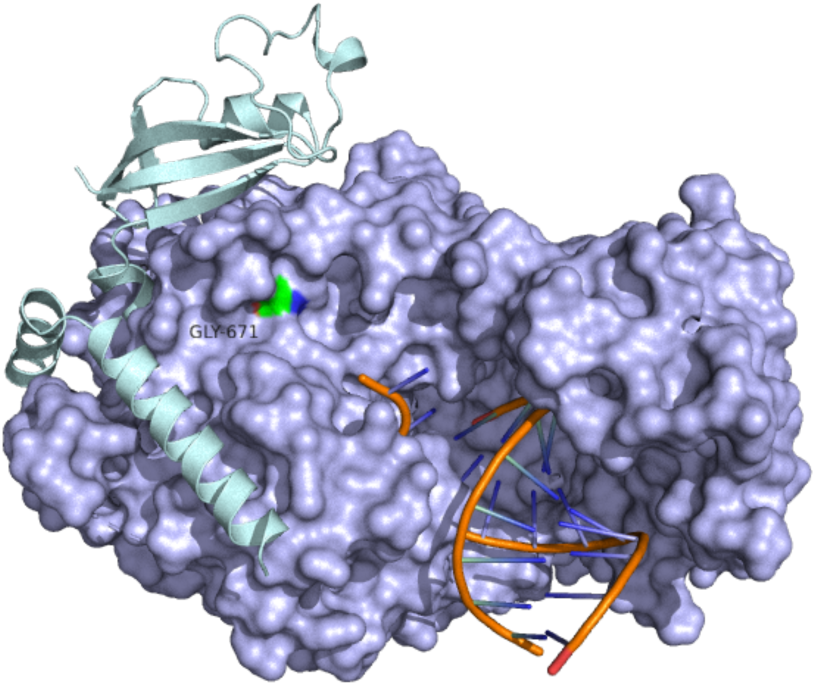
Nsp12 in association with Nsp8 and viral RNA. Surface representation of Nsp12. The position of Gly-671 is highlighted in green. Cartoon representation of Nsp 8 is shown in association, coloured in light green. Source PDB structure 7b3b.

*Nsp13: Helicase.*

PDB structure 7re2.

Delta Mutation. P77L. Proline to Leucine.

Blosum score of -3. Special case to hydrophobic.

ConSurf score of 8. Highly conserved residue.

ΔΔG^stability^ mCSM: -0.5 kcal/mol^-1^ (Destabilising).

This mutation is present in 98.8% of all Delta sequences.

BA.2 Mutation. R392C. Arginine to Cysteine.

Blosum score of -3. Positively charged to special case.

ConSurf score of 1. Highly variable residue.

ΔΔG^stability^ mCSM: -0.3 kcal/mol^-1^ (Destabilising).

This mutation is present in 99.06% of all BA.2 sequences.

Nsp13 consists of five domains arranged in a triangular shape. 1A, 2A and 1B form the base, which are connected via a stalk domain to the N-terminal zinc-binding domain (ZBD) at the apex (Fig 12). Nsp13 has been shown in complex with the SARS-CoV-2 replication complex (Nsp7, Nsp8 and Nsp12), with the implication that these interactions have potential implications for helicase activity.

Pro-77 is located at a bend in a loop in the ZBD. When in a complex, the ZBD has close association with the N-terminal helical extension of nsp8 (Fig13), although in this PDB structure there are no bonds between the area. Given the consurf score of this residue, there may be structural importance to this area of which we are unaware, although the domain as a whole appears to be flexible (10).

Arg-392 is located in the 1A domain at the bottom of the triangular structure. It does not form part of any helicase motifs, but in this particular PDB structure it makes several polar bonds both within the surrounding beta sheets, and with a residue on nearby nsp7 (Figs 14-15). This connection appears significant in terms of the overall replication and transcription complex. There are not many other locations on nsp13 where there is close contact between these proteins. On mutation to cysteine this polar bond is lost (Fig 16).

**Figure 12.**
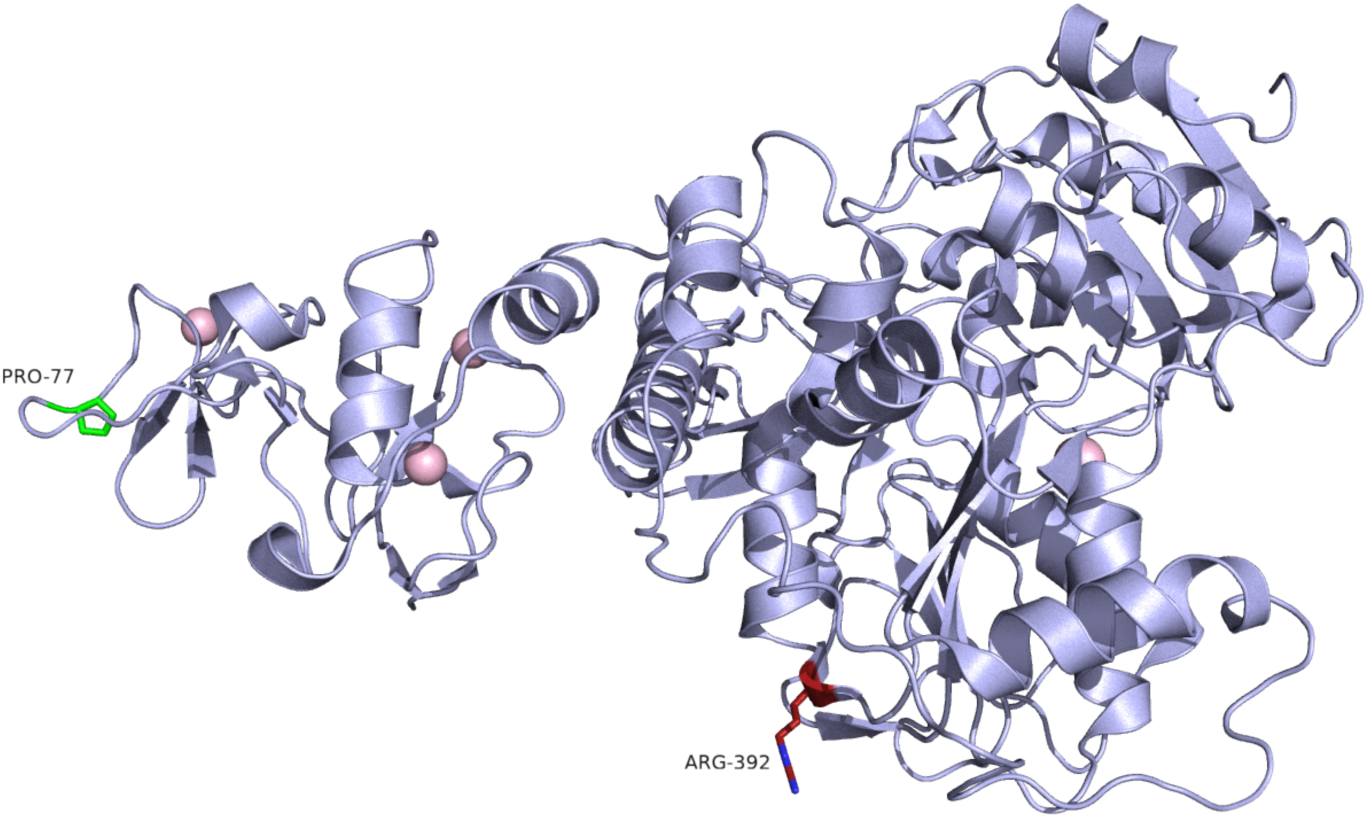
Nsp 13. Cartoon representation of Nsp 13. The Zinc binding domain is positioned on the left side of the image, where Pro-77 is highlighted in green. Arg-392 is highlighted in red within the 1A domain at the base of the triangle. Source PDB structure 7re2.

**Figure 13.**
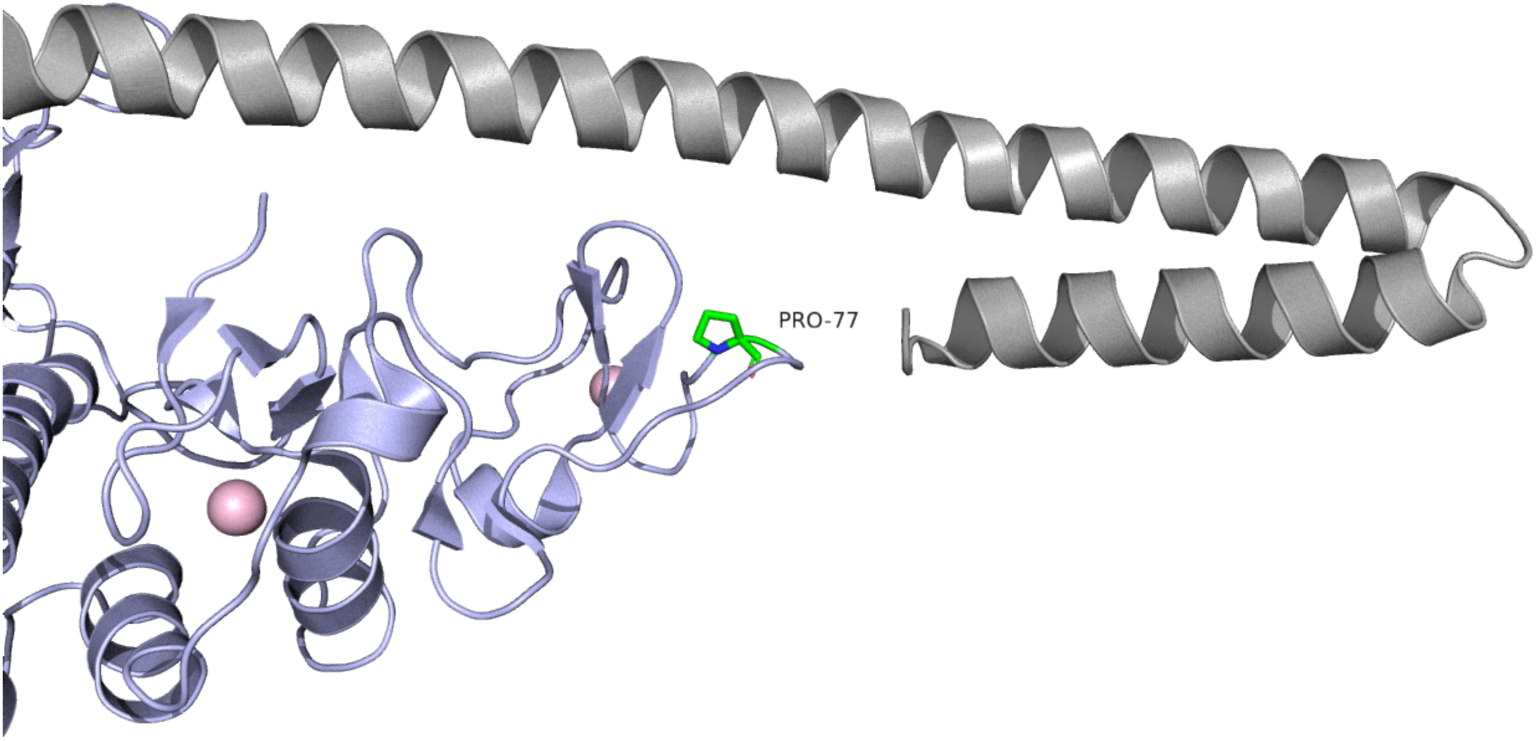
The Zinc-binding domain of Nsp13. Cartoon representation of the ZBD of Nsp13. Pro-77 is highlighted in green. The N-terminal helicase extension of nsp 8 is shown in gray. Source PDB structure 7re2.

**Figure 14.**
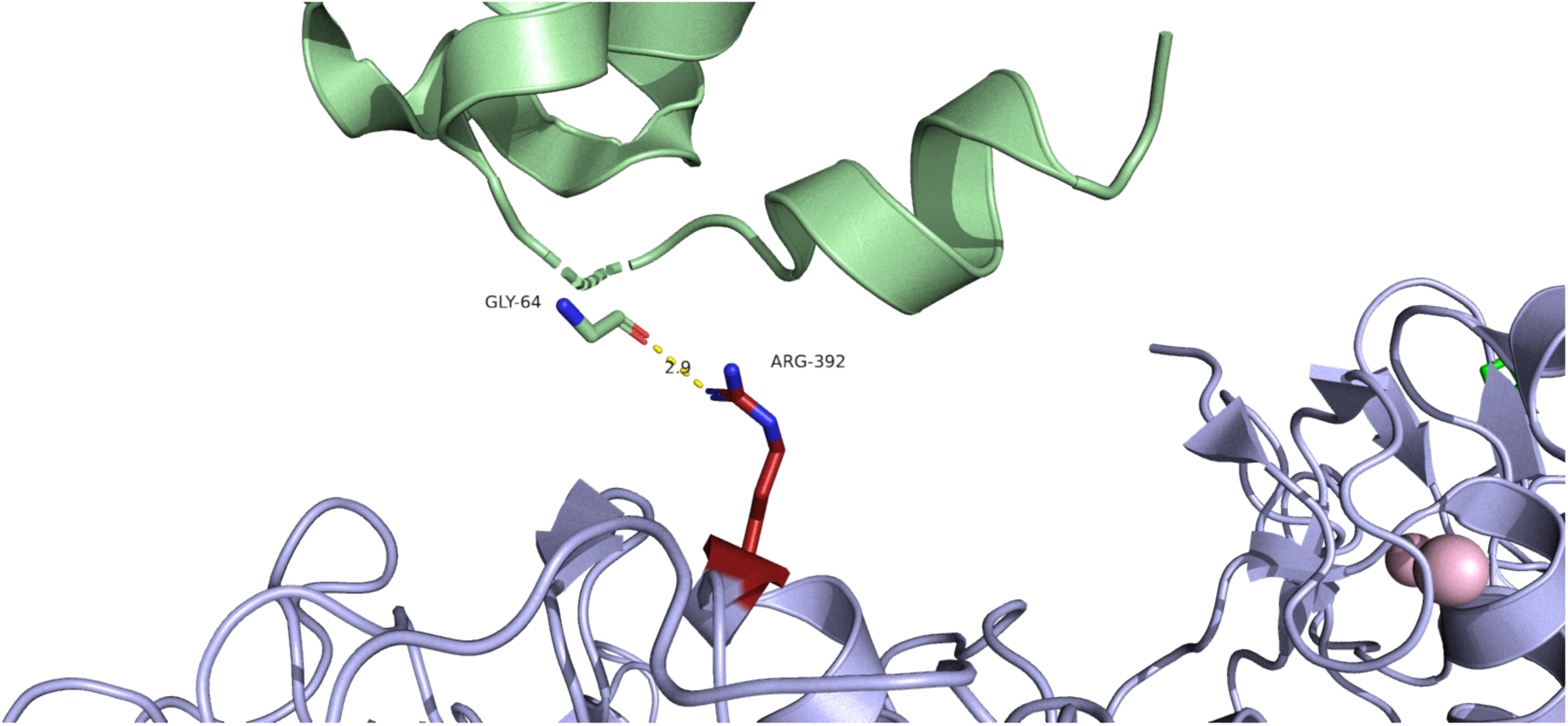
Nsp13 1A domain in association with Nsp7. Cartoon representation of Nsp13 (light blue) and nsp7 (light green): The polar bond between Arg-392 of nsp13 and Gly-64 of nsp7 is measured at 2.9 Å in Pymol. Source PDB structure 7re2.

**Figure 15.**
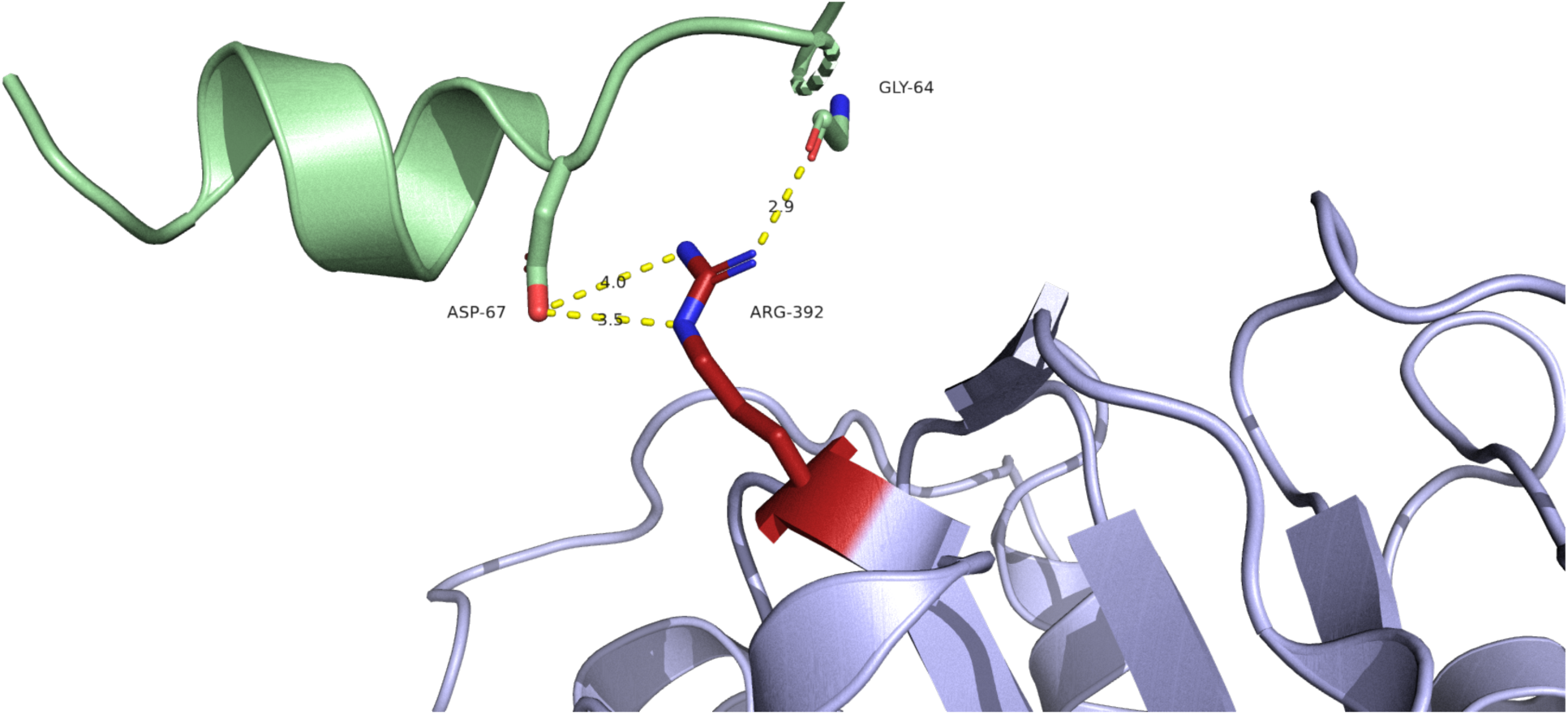
Nsp13 1A domain in association with Nsp7. Using mCSM-PPI2 to predict binding affinity within this PDB structure, we can see that wild type nsp13 forms not just a polar bond with GLY-64, but also two ionic bonds with ASP-67. There are no bonds of any type predicted when Arg-392 mutates to Cys-392. Source PDB structure 7re2.

**Figure 16.**
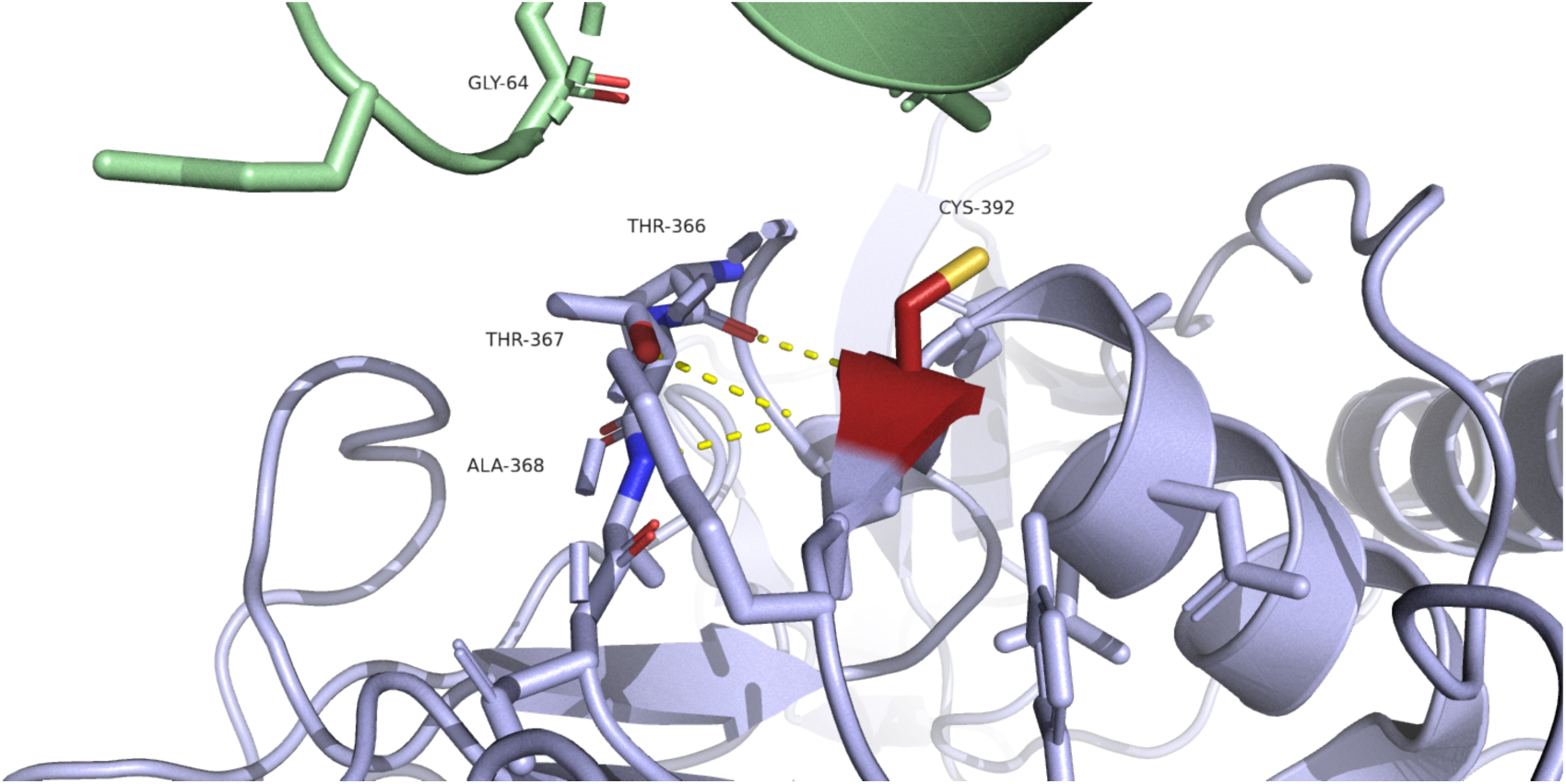
Nsp13 1A domain in association with Nsp7, with Cys-392 mutation. Cartoon representation of Nsp13 (light blue) and nsp7 (light green). Mutation to Cys-392 results in loss of polar bond to Gly-64. Source PDB structure 7re2.

Nsp13 interacts with TBK1, disrupting its association with MAVS. The Pro-77 mutation has been modelled in a molecular docking study to investigate how this might affect the binding of nsp13 with TBK1, with the conclusion that Pro-77 may result in greater affinity between nsp13 and TBK1, possibly enhancing inhibitory effects (Fig 17) (11).

**Figure 17.**
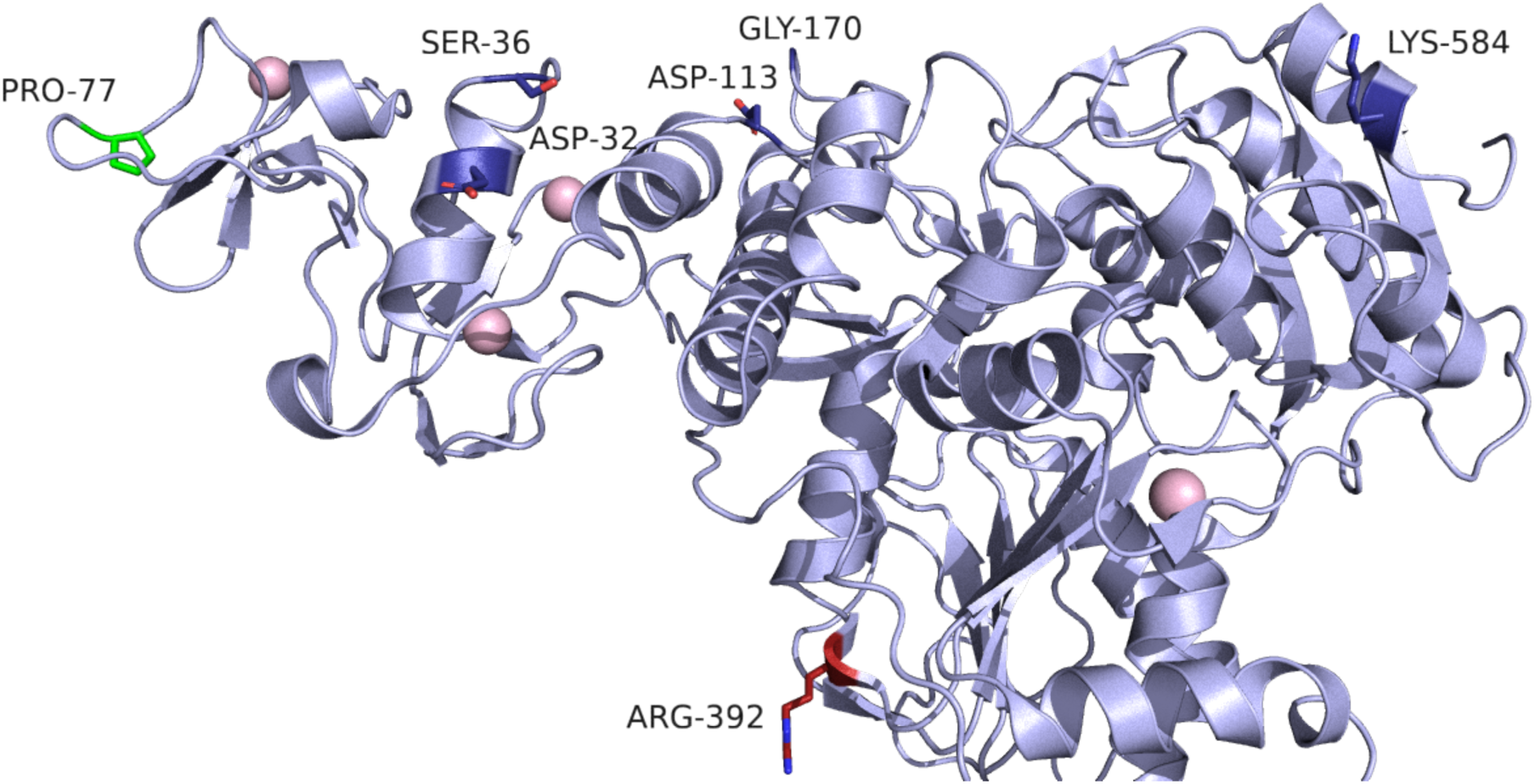
Key residues of Nsp13 thought to mediate association with TBK1. Cartoon representation of nsp13 in which key residues are coloured deep blue, at the top of the protein. Although pro-77 is located a little distance from these residues, it is on the same plane and the P77L mutation is predicted to enhance inhibitory action. Source PDB structure 7re2.

*Nsp14: Exonuclease*

PDB structure 7n0c.

BA.1/BA.2 Mutation. I42V. Isoleucine to Valine.

Blosum score of 3. Both hydrophobic.

ConSurf score of 5. Middle of range.

ΔΔG^stability^ mCSM: -1.06 kcal/mol^-1^ (Destabilising).

This mutation is present in 95% of all BA.1 sequences and 97.6% of all BA.2 sequences.

Nsp14 contains a C-terminal domain, which carries S-adenosyl methionine (SAM)-dependent N7-MTase activity and plays a role in viral RNA 5’ capping, facilitating viral mRNA stability and translation and preventing detection by innate antiviral responses.

It also contains an N-terminal ExoN domain (3’ to 5’ exoribonuclease activity). Nsp10 (a zinc binding protein) associates with nsp14 enhancing ExoN but not N7-MTase activity, through increased structural stability.

Mutations that abolish the nsp14-nsp10 interaction result in a lethal phenotype in SARS-CoronaVirus (i.e. the virus doesn’t survive/replicate).

Overexpression of nsp14 in an in vitro model reduced host cell translation and inhibited IFN-dependent induction of ISGs. Both domains were necessary for this inhibition, and association with nsp10 enhanced the inhibitory effect (12).

Ile-42 is located in the ExoN domain, although it is distant from the active site (Figs 18-19).

Specifically, it is located in a long flexible region which mediates association with nsp10 (Figs 20-21).

**Figure 18.**
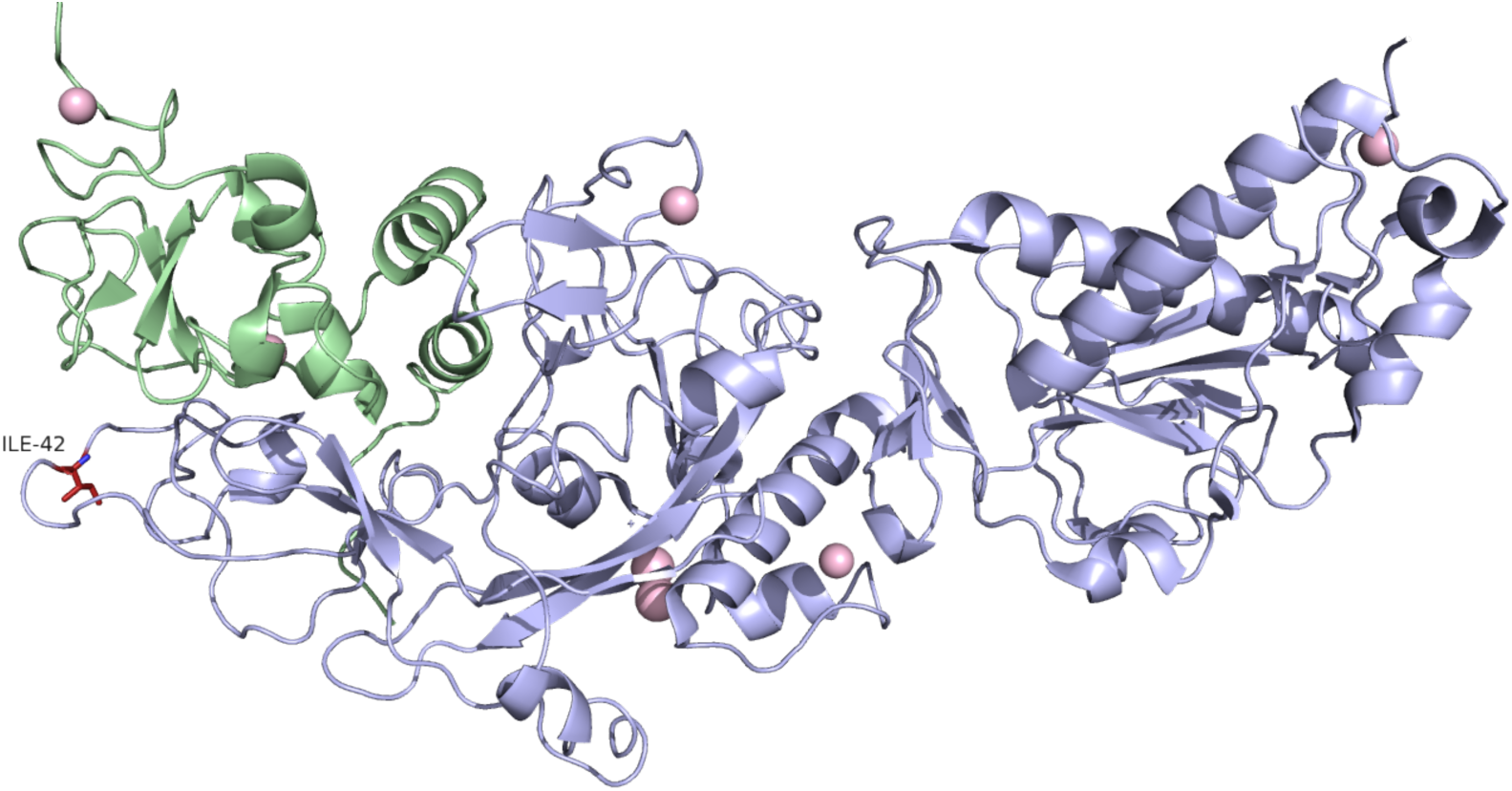
Nsp14 in association with Nsp10. Cartoon representation of nsp14 (light blue chain). The ExoN domain is on the left side of the image, N7-MTase is on the right. Ile-42 is highlighted in red, and forms part of the region that associates with nsp10 (light green). Source PDB structure 7n0c.

**Figure 19.**
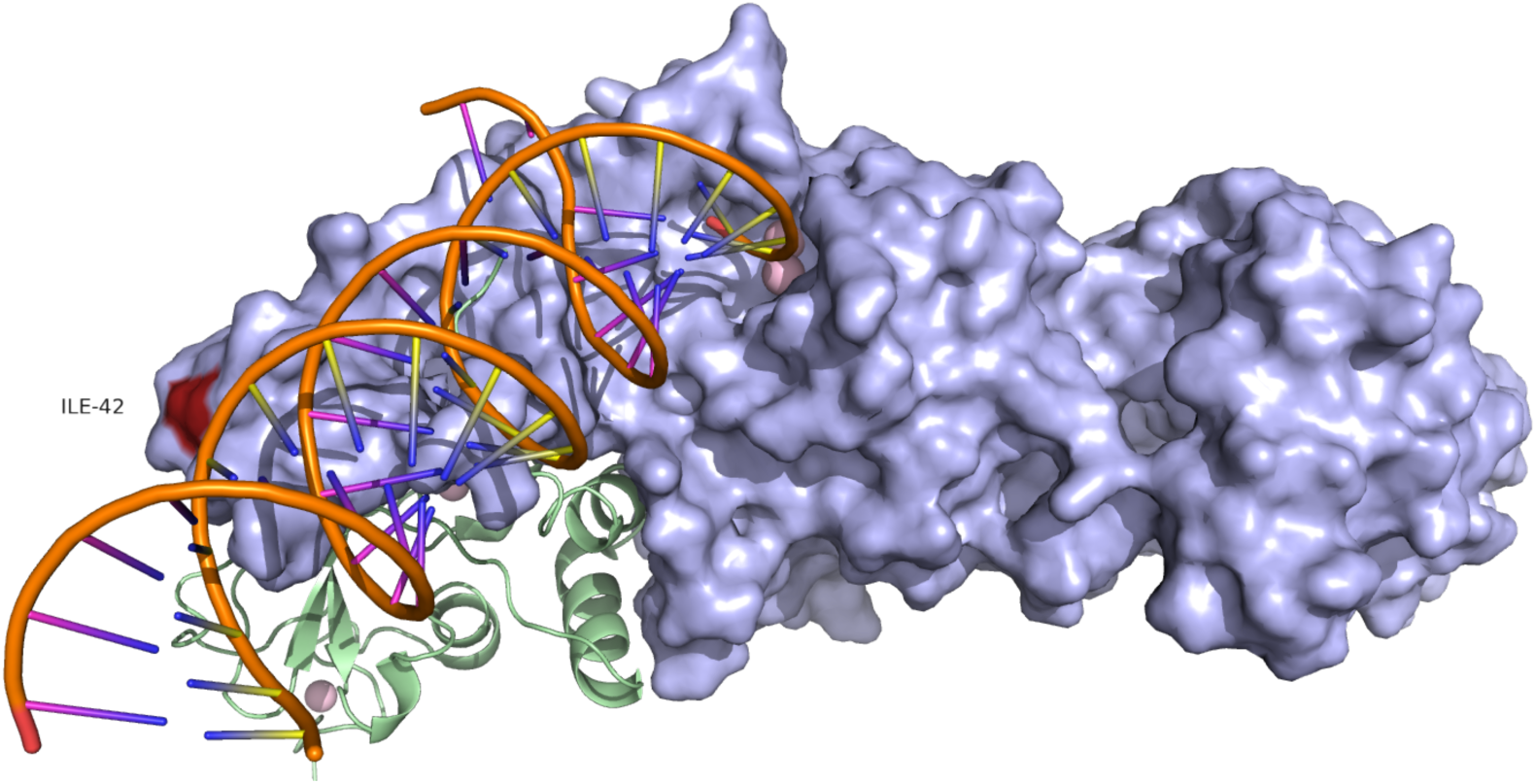
Nsp14 surface model. Surface model illustrates the RNA binding groove within the ExoN domain and distance from Ile-42 (highlighted in red). Source PDB structure 7n0c.

**Figure 20.**
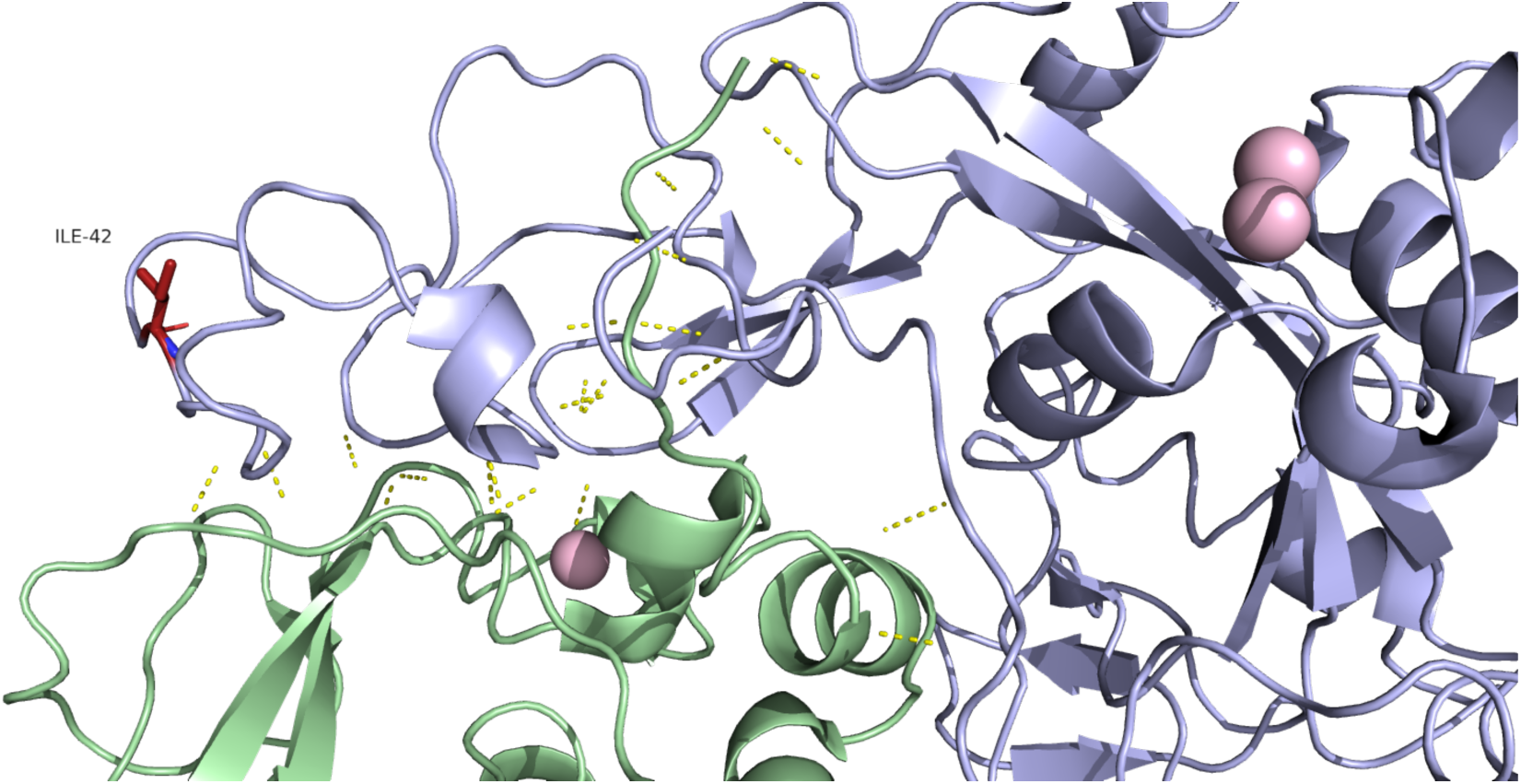
Nsp14 ExoN domain in association with Nsp10. Cartoon representation of Nsp14 (light blue) showing polar bonds between nsp10 (light green). Ile-42 is not among the residues involved. Source PDB structure 7n0c.

**Figure 21.**
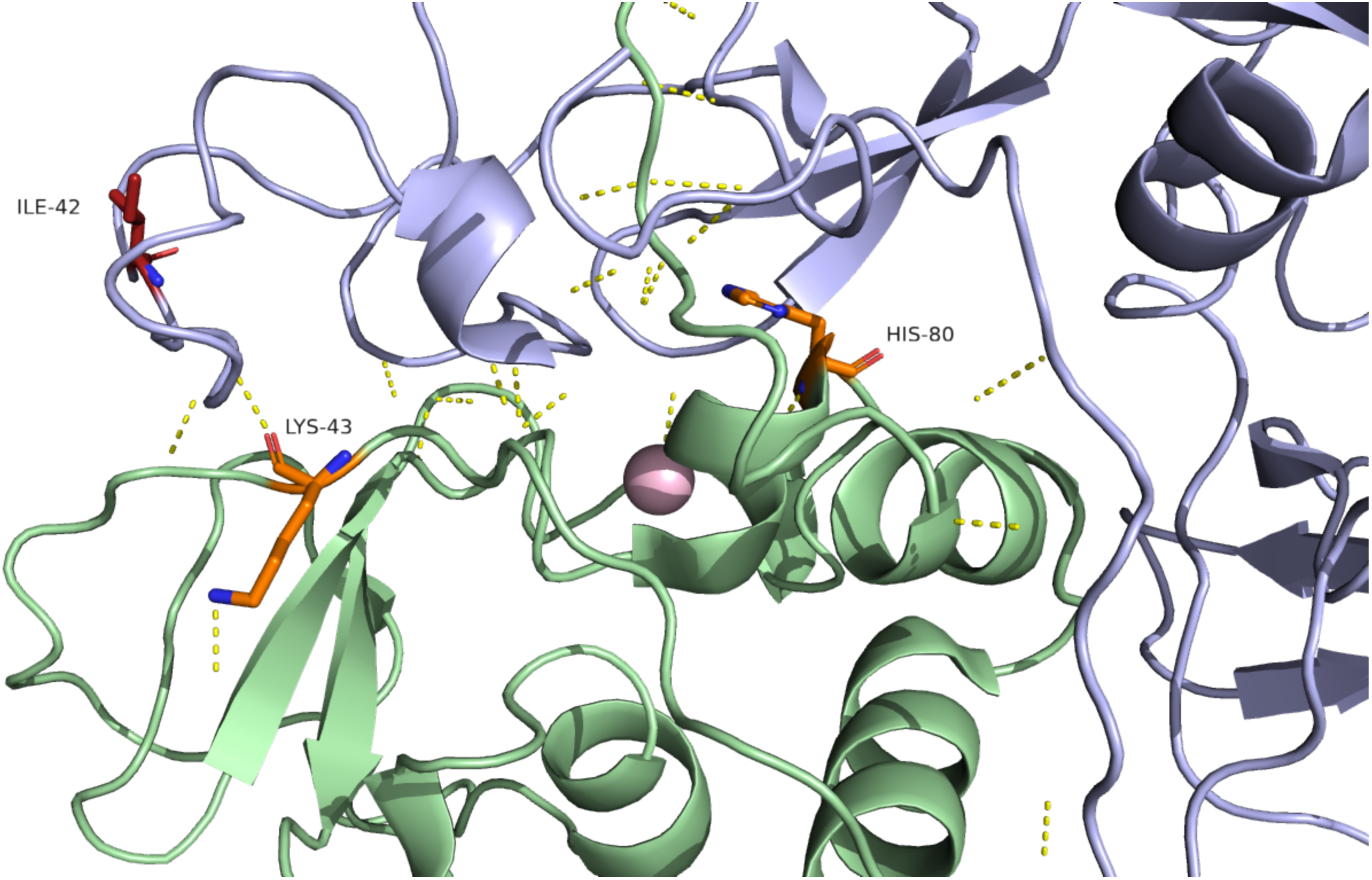
Nsp14 ExoN domain in association with Nsp10. Residues reported to be key to the interaction between Nsp10 and Nsp14 are Lys-43 and His-80 of Nsp10 (highlighted in orange above). There is no association between these residues and Ile-42. Source PDB structure 7n0c.

Running this mutation through mCSM-PPI2 shows that there is a very small increase in affinity (0.059 kcal/mol) predicted in terms of Ile-42 and surrounding residues in nsp14. This residue already makes multiple hydrophobic connections with surrounding residues, so this part of the protein appears structurally stable. There are no bonds with nsp10 involved. Therefore, this mutation appears to be of minimal consequence.

*Nsp15: EndoU/Endoribonuclease*

PDB structure 7tqv

BA.2 Mutation. T113I. Threonine to Isoleucine.

Blosum score of -1. Polar uncharged to hydrophobic.

ConSurf score of 7. Moderately conserved.

ΔΔG^stability^ mCSM: -0.054 kcal/mol^-1^ (Destabilising).

This mutation is present in 96% of all BA.2 sequences.

Structures in PDB are commonly arranged as a hexamer, a dimer of nsp15 trimers. There are three domains - an N terminal domain important for oligomerization, a variable middle domain, and an endonuclease domain. Nsp15 preferentially cleaves RNA substrates 3’ of uridines. It may regulate the length of polyuridines found at the 5’ end of negative strand viral RNA to evade activation of host innate immune responses (13)

Thr-113 is located in the variable middle domain. This part of the protein is not involved in nuclease activity or oligomerization. Thr-113 does not form any polar bonds with other residues in this chain or other chains within the proposed hexamer arrangement and this does not change upon mutation to Ile-113. The mutation therefore appears unlikely to have a substantial impact..

**Figure 22.**
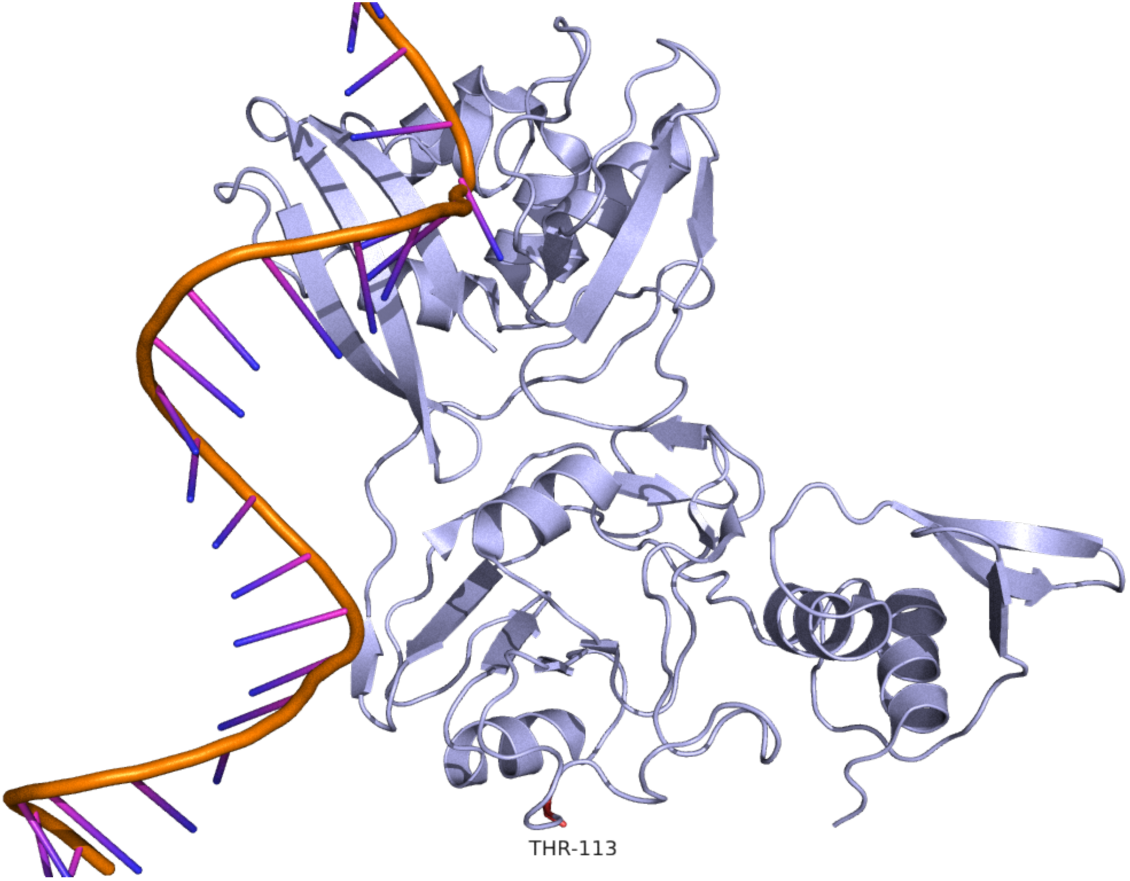
Cartoon of Nsp15. Cartoon representation of Nsp15 with viral RNA in association. Thr-113 is highlighted in red.

### ORF3a

PDB structure 7kjr.

Delta Mutation. S26L. Serine to Leucine.

Blosum score of -2. Polar uncharged to hydrophobic.

ConSurf score of 1. Highly variable.

ΔΔG^stability^ mCSM: -0.27 kcal/mol^-1^ (Destabilising).

This mutation is present in 99.2% of all Delta sequences.

BA.2 Mutation. T223I. Threonine to Isoleucine.

Blosum score of -1. Polar uncharged to hydrophobic.

ConSurf score of 9. Highly conserved.

ΔΔG^stability^ mCSM: -0.23 kcal/mol^-1^ (Destabilising).

This mutation is present in 99.3% of all BA.2 sequences.

As a transmembrane protein, ORF3a has two domains, a tall transmembrane domain and a cytosolic domain. The cytosolic domain is mainly composed of beta sheets, and the BA.2 mutation is located at the bottom edge of the last of these. According to modelling in Pymol the mutation appears to reduce the number of polar bonds this residue forms with neighbouring residues in the beta-sheet from 3 to 1, although somewhat contradicting this the 3-D Covid site predicts an increase in hydrophobic connections and hydrogen bonds on mutation. Thr-223 is not close to an ion channel, but could potentially cause some disruption to dimer formation since it is located at a point of interaction between the two chains (Fig 24).

Ser-26 is located at the C-terminal and is not included in any PDB structures to date. It appears to be a highly variable region where mutation will be of little consequence.

ORF3a is thought to upregulate SOCS1, a negative regulator of cytokine signalling, and through this action, inhibit JAK/STAT signalling. A series of experiments using truncated versions of ORF3a showed that the residues essential to IFN inhibitory activity are 70-130, which form two of the three alpha-helices. None of the mutations listed above fall within this range (14).

**Figure 23.**
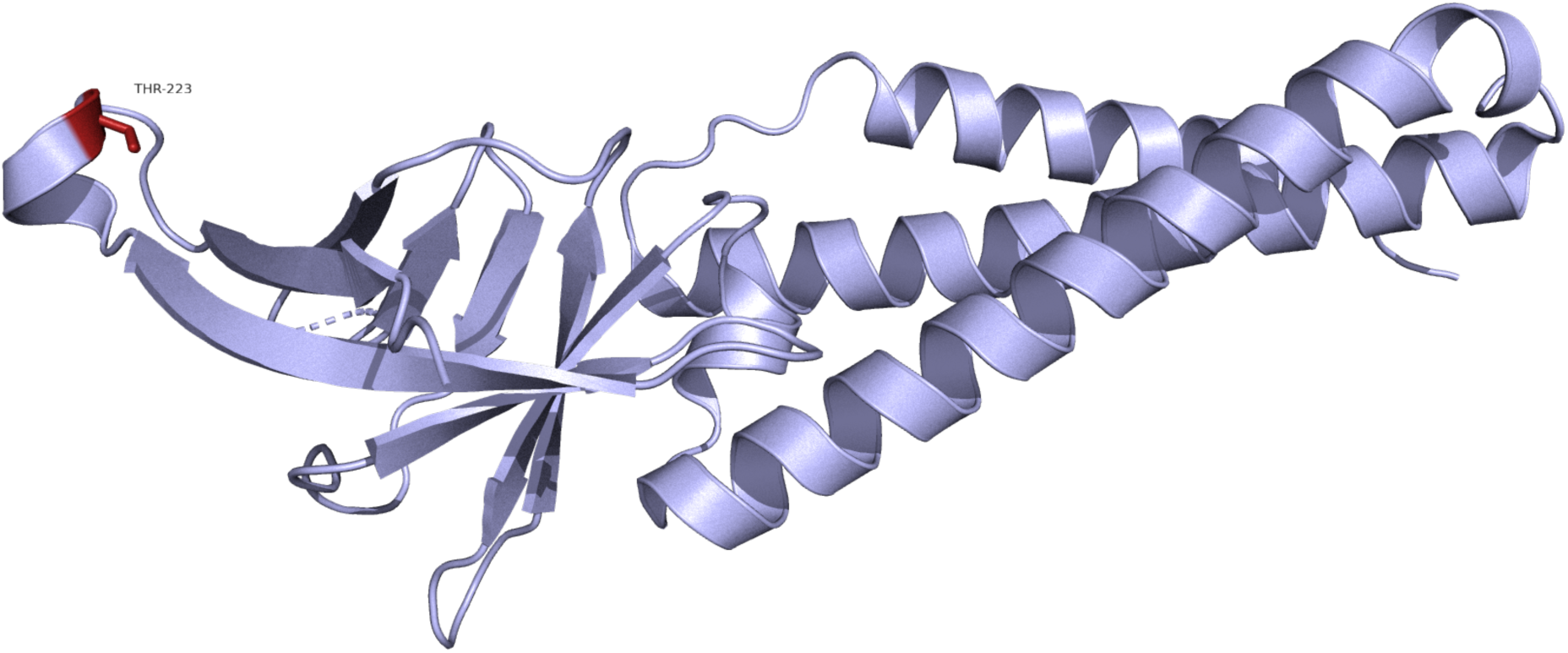
ORF3a protomer. Cartoon of ORF3a monomeric form. The cytosolic domain on the left of the image is composed of beta sheets, whilst the transmembrane domain is formed of three long alpha helices. Thr-223 is highlighted in red. Source PDB structure 7kjr.

**Figure 24.**
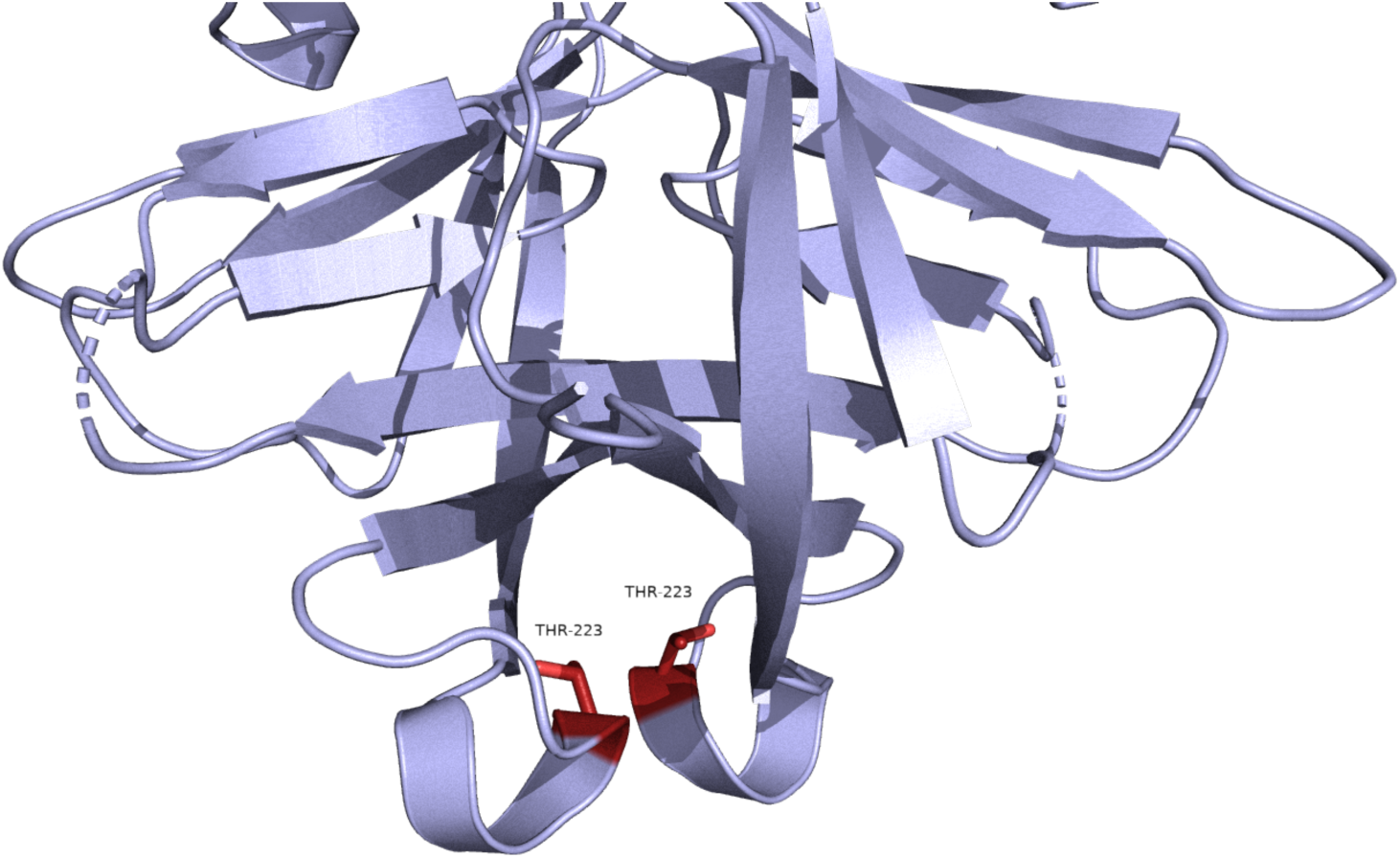
Cytosolic domain of the ORF3a homodimer. Cartoon representation of ORF3a cytosolic domain, arranged as a homodimer. Thr-223 is highlighted in red, showing the close association of this region of the protomer. Source PDB structure 7kjr.

### M protein

Structure from AlphaFold.

AlphaFold model shows transmembrane protein structure similar to protein 3a, there is a tall transmembrane domain composed of alpha-helices with a shorter cytosolic domain composed of beta sheets.

BA.1/BA.2 Mutation. Q19E. Glutamine to Glutamic acid.

Blosum score of 2. Uncharged to negative charge.

ConSurf score of 6. Moderately conserved.

ΔΔG^stability^ mCSM: -0.81 kcal/mol^-1^ (Destabilising).

This mutation is present in 96.0% of all BA.1 and 95.3% of all BA.2 sequences.

BA.1/BA.2 Mutation. A63T. Alanine to Threonine.

Blosum score of 0. Hydrophobic to polar uncharged.

ConSurf score of 7. Moderately conserved.

ΔΔG^stability^ mCSM: -1.42 kcal/mol^-1^ (Destabilising).

This mutation is present in 97.2% of all BA.1 and 98.5% of all BA.2 sequences.

Delta Mutation. I82T. Isoleucine to Threonine.

Blosum score of -1. Hydrophobic to polar uncharged.

ConSurf score of 8. Highly conserved.

ΔΔG^stability^ mCSM: -2.9 kcal/mol^-1^ (Destabilising).

This mutation is present in 98.9% of all Delta sequences.

All of these mutations are located within the transmembrane domain, each on separate alpha-helices. They are all located towards the top of the domain, each corresponding with a transmembrane motif. The transmembrane domain is thought to interact with RIG-1, ultimately inhibiting production of type I and type III IFN production induced by RIG-I/MDA5 (15).

From Pymol modelling, Iso-82 increases polar bonds from 3 to 4 on mutation, whereas Glu-19 and Ala 63 decrease theirs, from 3 to 2 and 4 to 2 respectively. 3-D Covid predictions are largely in concordance with this except the prediction for Iso-82 is a large reduction in stability due to loss of hydrophobic connections and the introduction of several clashes. Overall this suggests that all three mutations might decrease stability in these motif areas. So, despite blosum scores indicating that these are mutations of low impact, their positioning may lead them to having greater significance.

**Figure 25.**
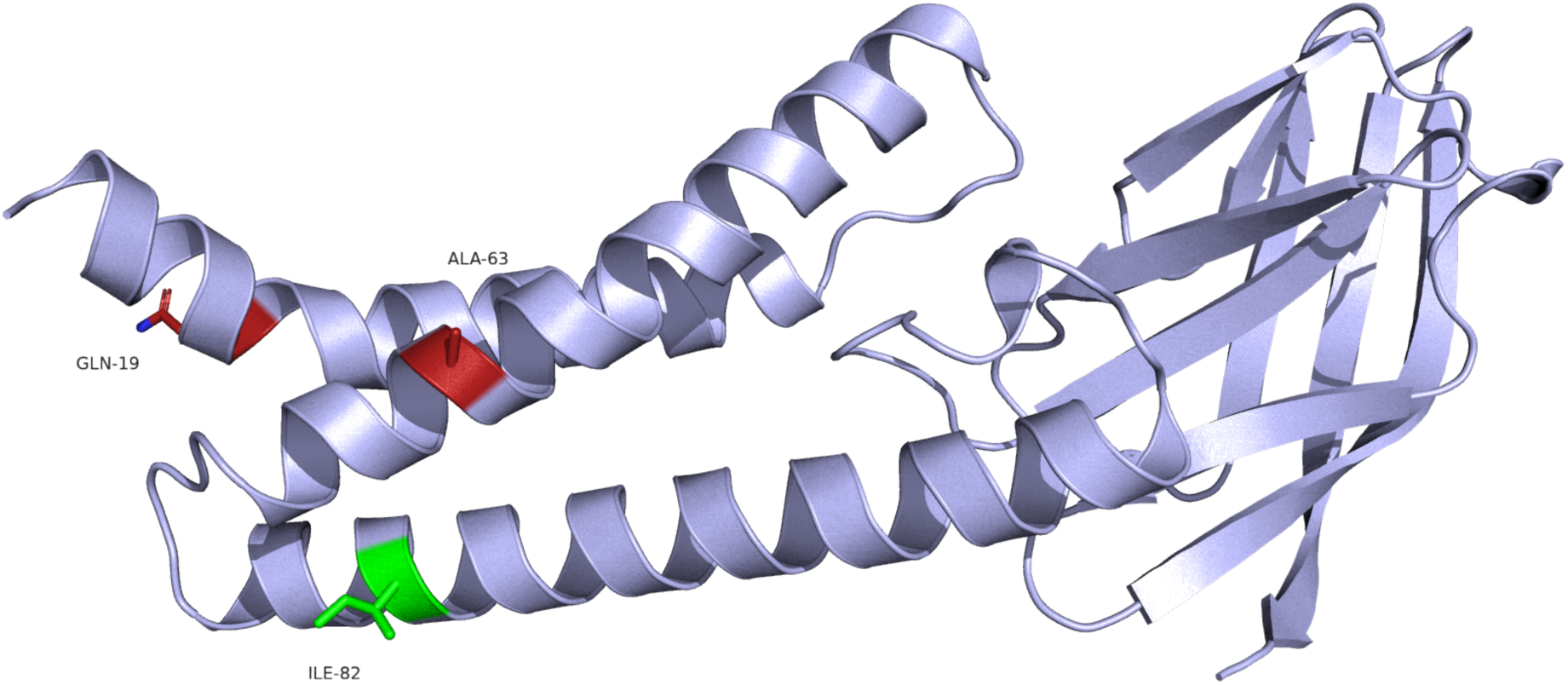
M protein. Cartoon representation of M protein, showing residues of interest positioned on three separate alpha helices, relating to three transmembrane motifs. BA.1/BA.2 residues are highlighted in red, whilst Delta residue Ile-82 is highlighted in green. Source structure from AlphaFold.

**Figure 26.**
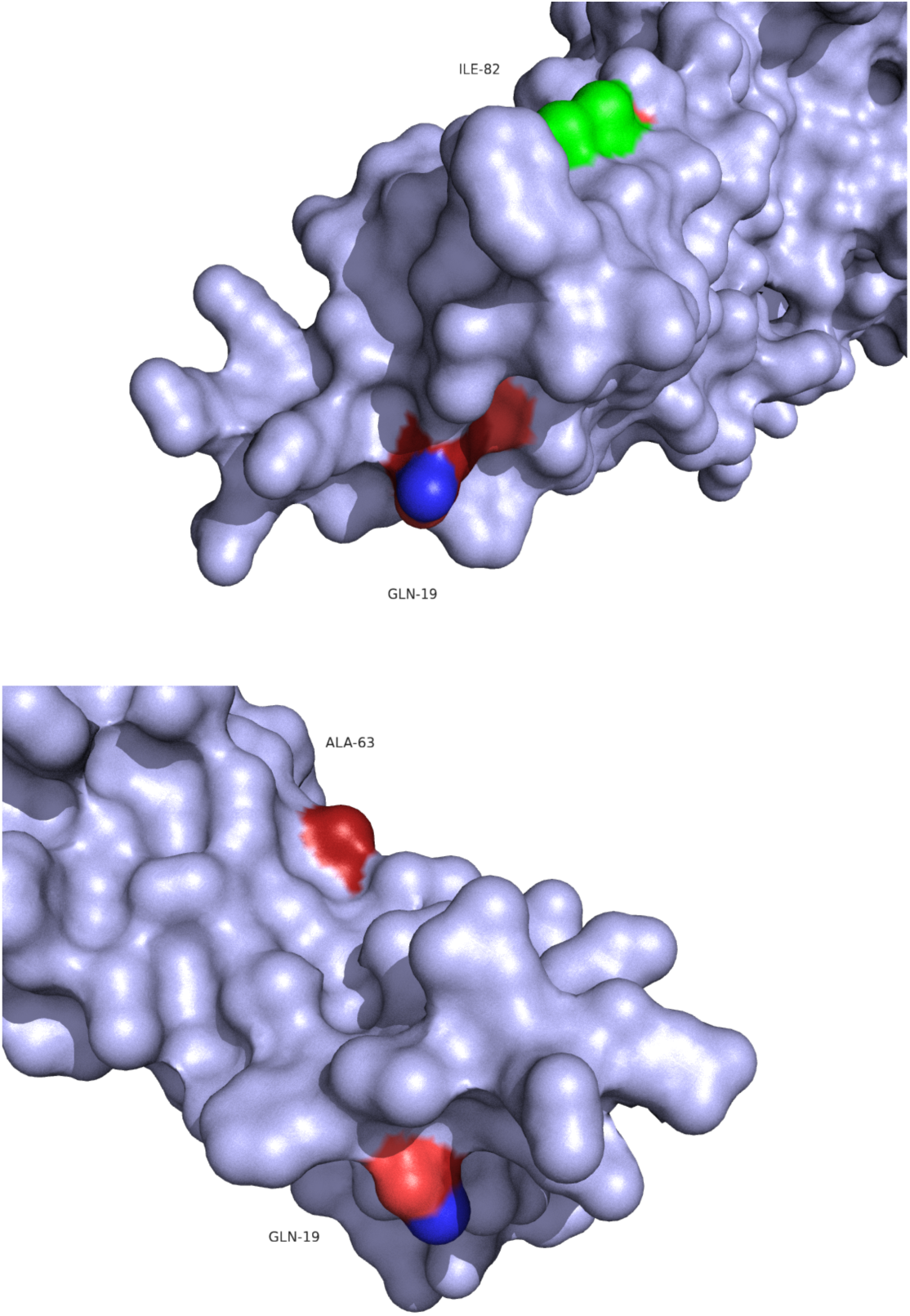
M protein transmembrane domain. Surface models of transmembrane domain of M protein. Delta residue is highlighted in green, whilst BA.1/BA.2 residues are highlighted in red. All three residues are located on transmembrane motifs.

### ORF6

PDB structure 7vph.

BA.2 Mutation. D61L. Aspartic acid to Leucine.

Blosum score of -4. Negatively charged to hydrophobic.

ConSurf score of 9. Highly conserved.

ΔΔG^stability^ mCSM: 0.73 kcal/mol^-1^ (Stabilising).

This mutation is present in 97.6% of all BA.2 sequences.

There is only one model of this structure in PDB covering the C-terminal (residues 53-61), and here it is shown with a ribonucleic acid export 1 (Rae1)– nucleoporin 98 (Nup98) complex. It is thought that ORF6 antagonises host interferon signalling through association with this complex, tight binding competitively inhibits binding of host RNA and subsequent export (16,17)

This change has the potential to be significant. The C-terminal of Orf6 binds to the RNA binding pocket of the Rae1-Nup98 complex. There are polar bonds along the length of this C-terminal domain, but the area of greatest binding is with Asp-61, which forms 5 polar bonds with 3 residues on Rae1 (Fig 28). In contrast Leucine retains just 1 bond with one of the residues (Fig 30).

**Figure 27.**
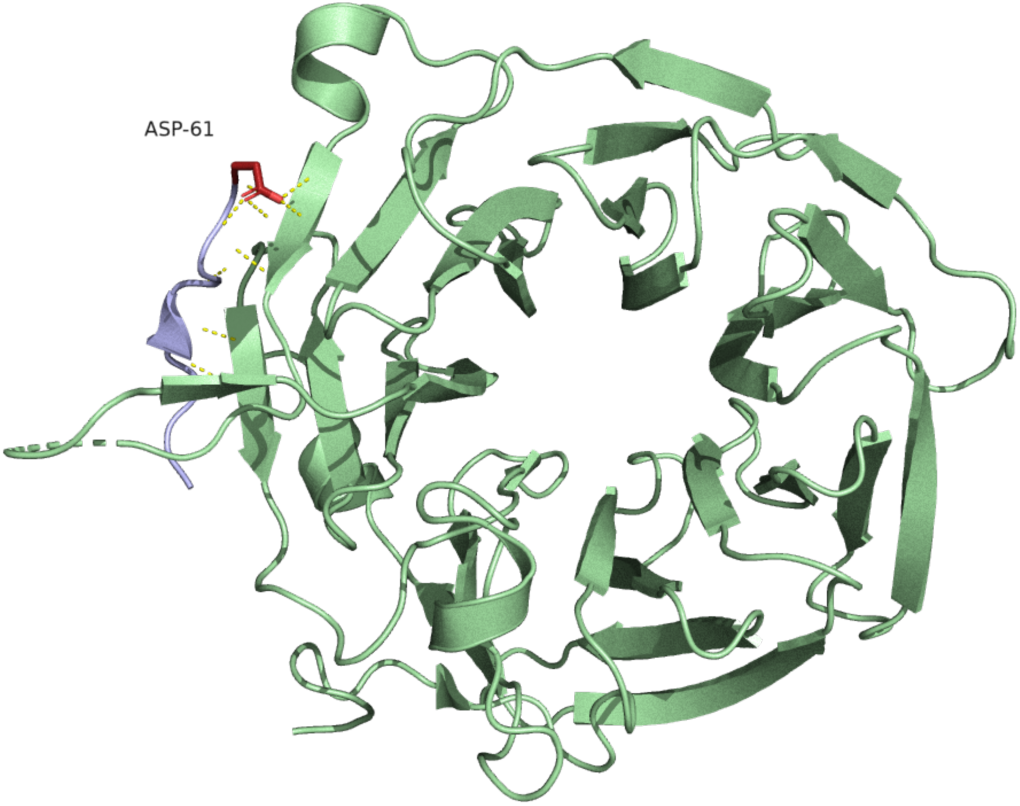
C-terminal of ORF6 bound to Rae-1. Cartoon representation of Rae-1 (light green) bound to the C-terminal domain of ORF6 (light blue). Asp-61 is highlighted in red. Source PDB structure 7vph.

**Figure 28:**
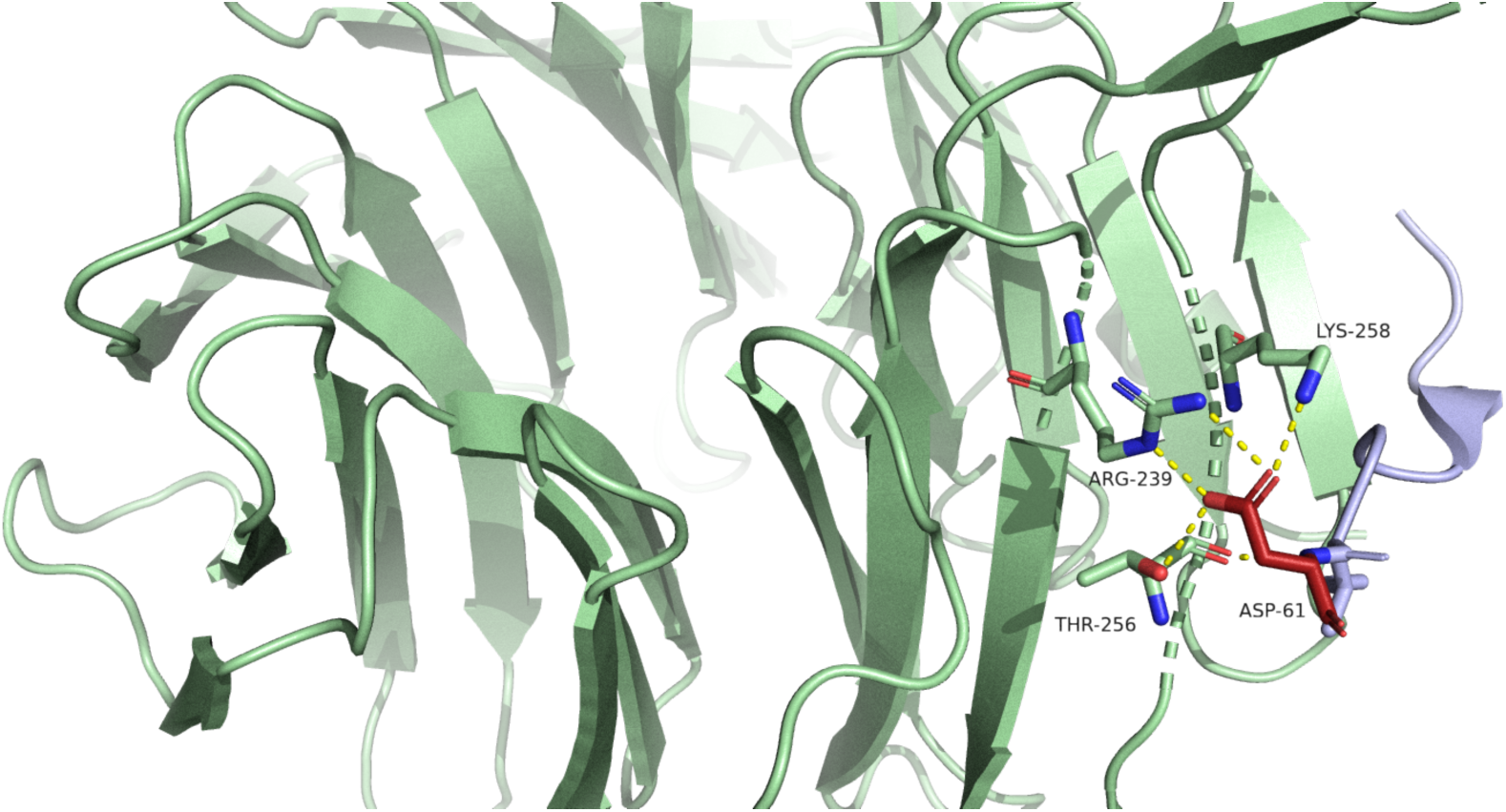
Close view of C-terminal of ORF6: Asp-61 polar bonds with Rae-1. Cartoon representation of ORF6 bound to Rae-1. Asp-61 makes four polar bonds from, with Arg-239, Thr-256, and Lys-258 on Rae-1. Source PDB structure 7vph.

**Figure 29.**
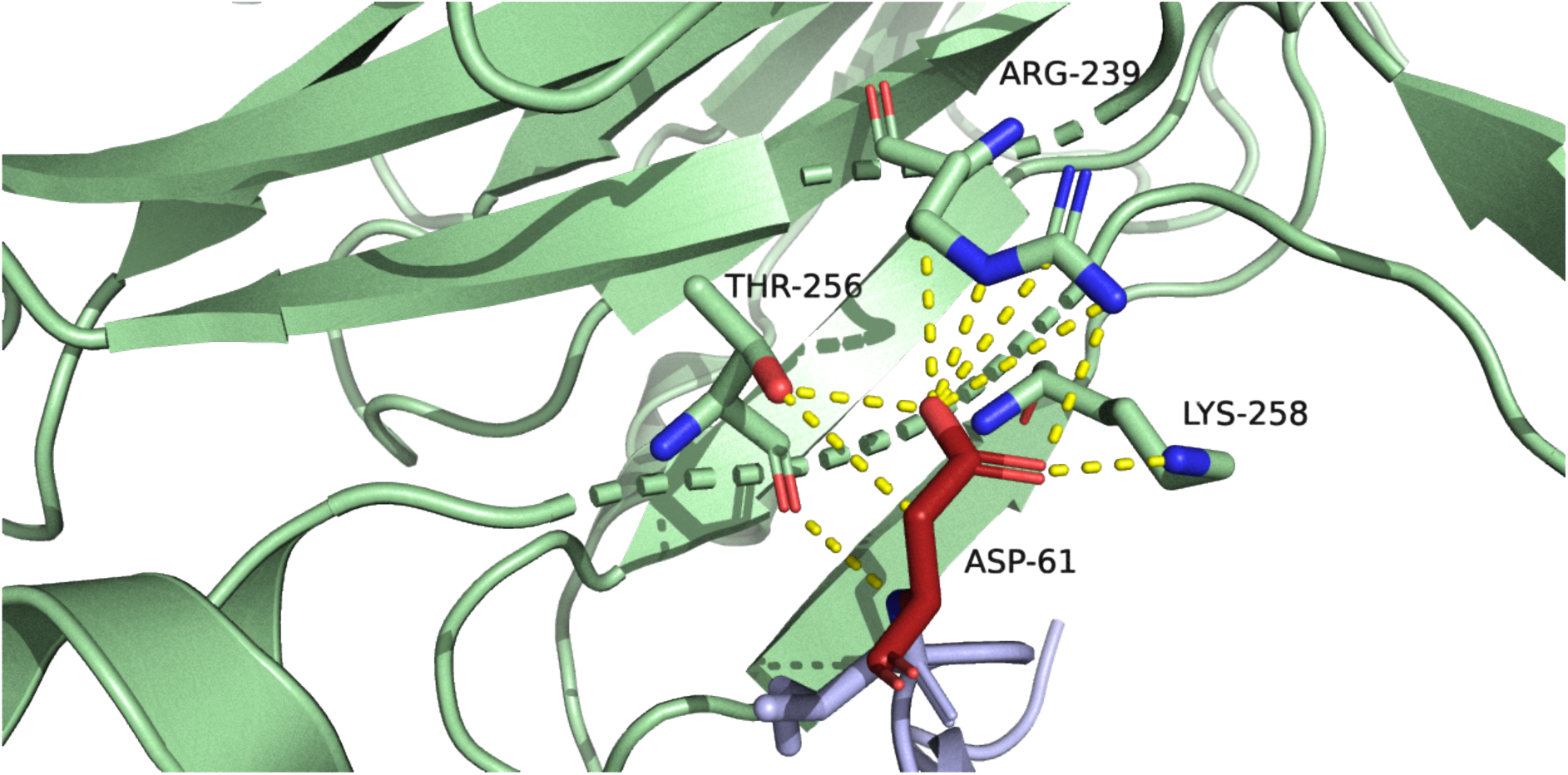
Close view of C-terminal of ORF6: Asp-61 additional bonds with Rae-1. Cartoon representation of ORF6 bound to Rae-1. mCSM-PPI2, predicts a total of seven polar bonds: three between Asp-61 and Thr-256, three between Asp-61 and Arg-239, (and two ionic bonds from nitrogen and carbon atoms of Arg-239 to an oxygen of Asp-61), and finally one between Asp-61 and Lys-258. Source PDB structure 7vph.

**Figure 30.**
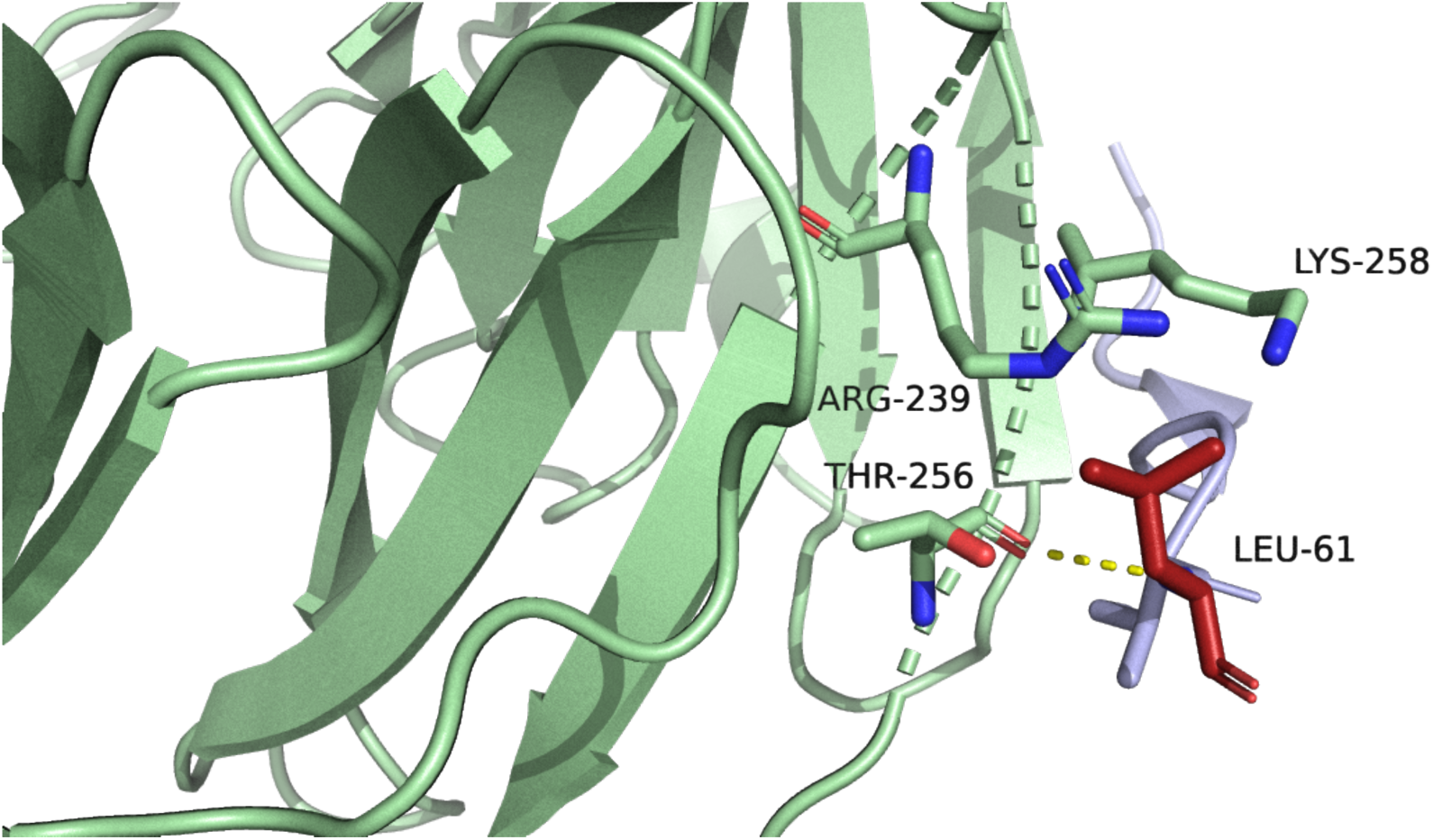
Close view of C-terminal of ORF6: Mutation to Leu-61 leads to reduced binding affinity with Rae-1. Cartoon representation of ORF6 bound to Rae-1 showing reductions to polar bonds when Asp-61 is substituted with Leu-61. mCSM-PPI2 predicts two hydrogen bonds in addition to the polar bond shown above connecting Leu-61 to Thr-256, and a total reduction in affinity between these proteins by -1.031 kcal/mol. Source PDB structure 7vph.

### ORF7a

PDB structure 7ci3.

Delta Mutation. V82A. Valine to Alanine.

Blosum score of 0. Both hydrophobic.

ConSurf score of 1. Highly variable.

ΔΔG^stability^ mCSM: -0.5 kcal/mol^-1^ (Destabilising).

This mutation is present in 93.3% of all Delta sequences.

This PDB structure shows the Ig-like ectodomain of the transmembrane protein 7a. This domain is bracketed by an N-terminal signalling region and a hydrophobic transmembrane domain containing a short ER retention motif, but no models are available of these domains.

Val-82 is located on a loop extended from a sequence of beta sheets, before connecting to the transmembrane domain. ORF7a is thought to block STAT2 phosphorylation, although details on which domains are implicated in this are currently unclear (18). The Ig-like ectodomain of ORF7a may directly interact with monocytes and modulate their antigen presenting ability (19). Key residues within this domain are located on and around the beta sheets, and Val-82 falls outside of this area. Its mutation does not alter the polar bonding it has with Arg-80. 3-D Covid predicts hydrophobic associations with Leu-88 (not shown in this model), also unaffected upon mutation. Therefore, this mutation appears to be of little significance.

**Figure 31.**
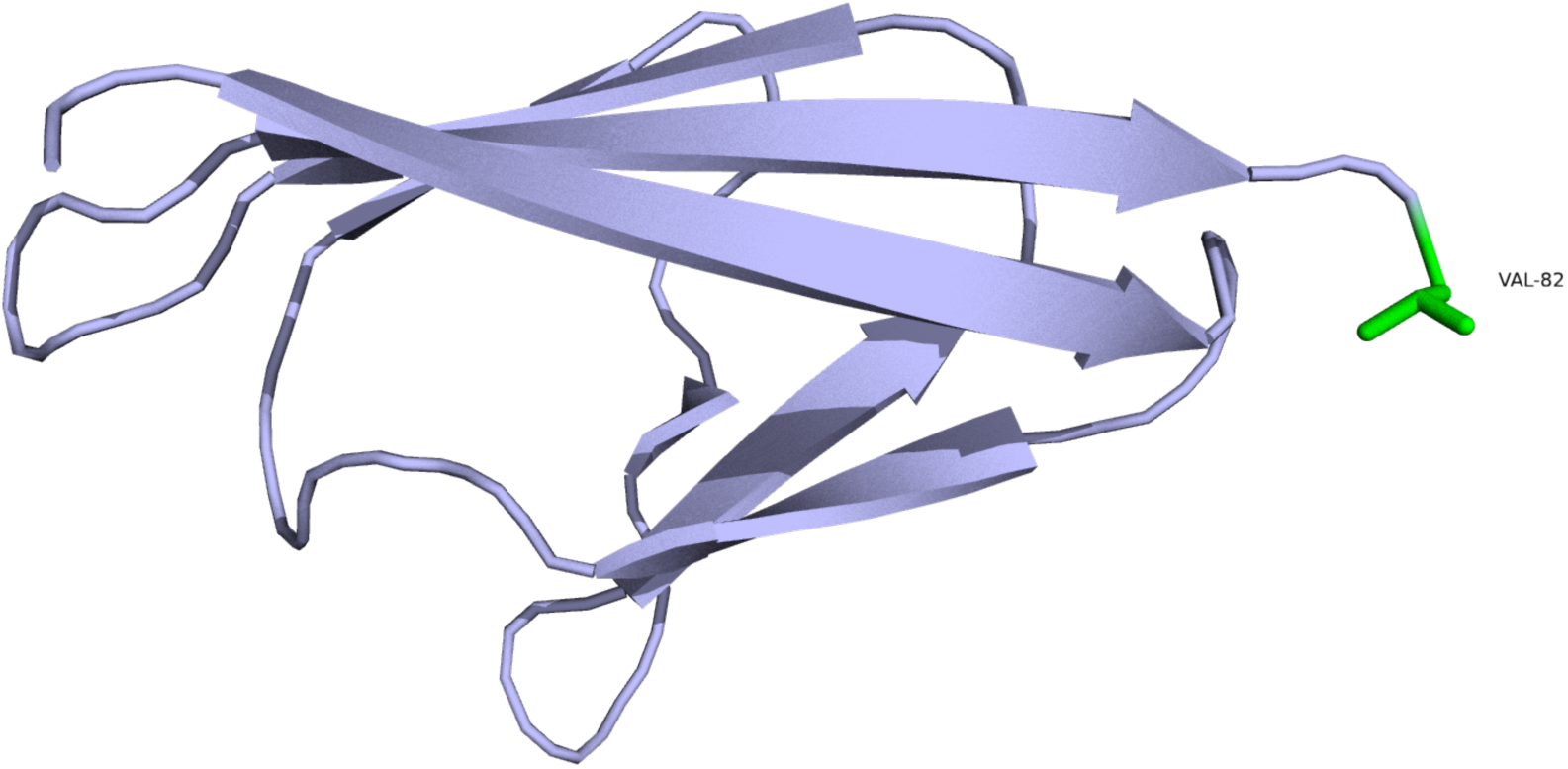
The Ig-like ectodomain of ORF7a. Cartoon representation of the Ig-like ectodomain of ORF7a. Val-82 is highlighted in green. Source PDB structure 7ci3.

### Nucleocapsid protein

*N-terminal binding domain.*

PDB structure 7act.

Delta Mutation. D63G. Aspartic acid to Glycine.

Blosum score of -1. Negative charge to special case.

ConSurf score of 1. Highly variable.

ΔΔG^stability^ mCSM: 0.44 kcal/mol^-1^ (Stabilising).

This mutation is present in 96.9% of all Delta sequences.

There are 8 other mutated residues in Supplementary Table 1, but none of the existing models cover these residues.

Asp-63 is located on a loop, seemingly away from main structural elements of this part of the protein, yet a proposed RNA binding groove lies in this domain over a nearby loop and Asp-63 is positioned at one end of this groove. The N-terminal domain has also been shown to be sufficient to suppress the activation of ISRE promoter, and reduce levels of p-STAT1 (20). There is a large reduction in residue size with this mutation, and there is potential for a change in the stability of RNA binding, but ConSurf results suggest residue variability at this site is common suggesting that the impact of this mutation may be minimal.

**Figure 33.**
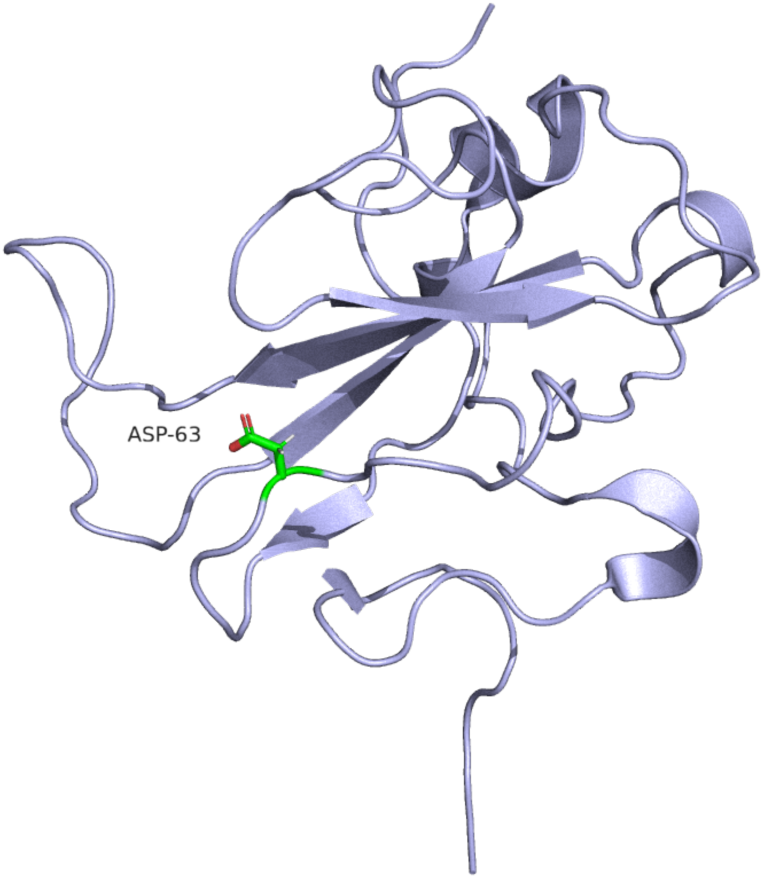
The N-terminal domain of N protein. Cartoon representation of the N-terminal binding domain of N-protein. Asp-63 is highlighted in green. Source PDB structure 7act.

**Figure 34.**
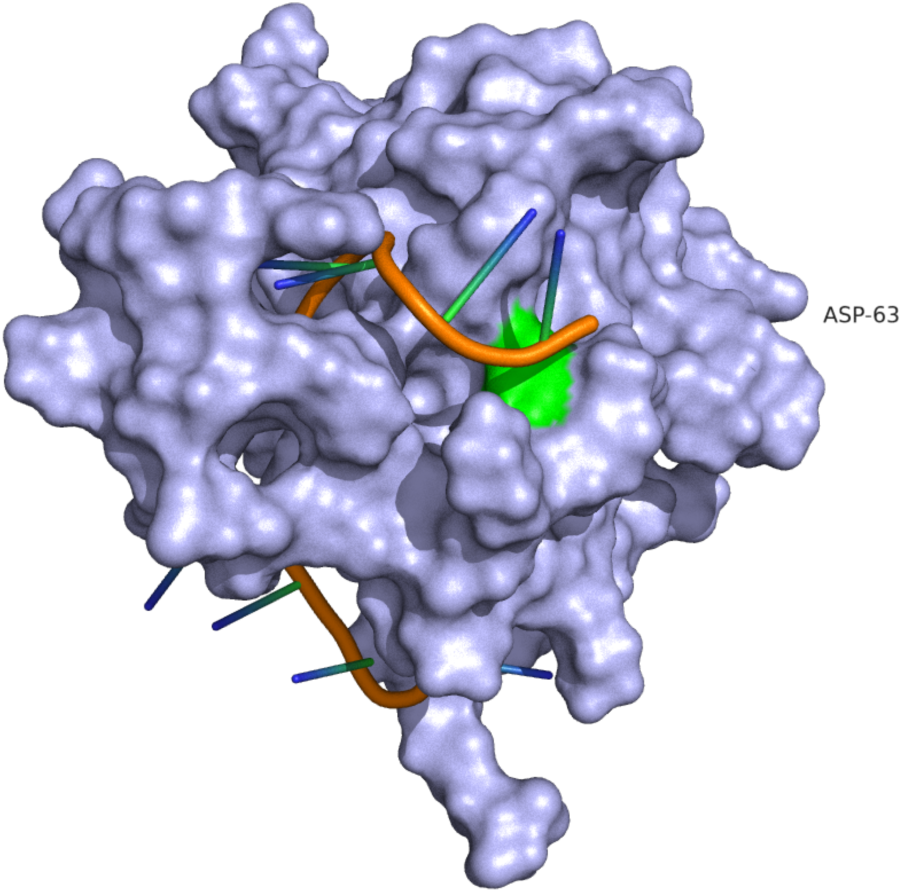
The N-terminal domain of N protein. Surface model of N-terminal binding domain of N protein, with RNA located in binding groove. Asp-63 is highlighted in green.

### Spike Protein

PDB Structures 6vxx and 7fg3.

S assembles as a homotrimer when acting as transmembrane proteins on virions. Two subunits S1 and S2 are separated by a cleavage site S1/S2. S1 contains a receptor binding domain which mediates ACE2 binding, whilst S2 contains segments that mediate fusion with host cell membranes. S^2^ represents a second cleavage site with S2.

The S1 region of S protein is highly variable, being most exposed to neutralising antibodies, and S as a whole is subject to intense research scrutiny. Consequently, there are over 800 structures available on PDB. 6vxx was selected to model mutations of interest as this cryo-EM structure was determined from S protein expression generated using the reference strain sequence. 7fg3 was also selected as it contained one of the most complete structures available.

S protein may influence host innate interferon responses. Cell fusion and syncytia formation appears to induce activation of cyclic GMP-AMP synthase (cGAS), and its downstream effector (STING), stimulating expression of IFN-beta. The S2’ cleavage site appears to be essential to syncytia formation, with S1/S2 cleavage site enhancing this activity but not being essential to it. Removal of S2’ sequence results in absence of cGAS-STING pathway activation (21).

However, there are no mutations located in or around the S2’ region in any of our isolates.

*S1/S2 cleavage site*

BA.1/BA.2 Mutation. N679K: Asparagine to Lysine.

Blosum score of -1. Uncharged to positively charged.

ConSurf score of 2. Highly variable.

ΔΔG^stability^ mCSM: 0.04 kcal/mol^-1^ (Neutral).

This mutation is present in 98.5% of all BA.1 and 99.8% of all BA.2 sequences.

BA.1/BA.2 Mutation. P681H: Proline to Histidine.

Blosum score of -2. Special case to positively charged.

ConSurf score of 1. Highly variable.

ΔΔG^stability^ mCSM: -0.34 kcal/mol^-1^ (Destabilising).

This mutation is present in 98.0% of all BA.1 and 99.5% of all BA.2 sequences.

Delta Mutation. P681R: Proline to Arginine.

Blosum score of -2. Special case to positively charged.

ConSurf score of 1. Highly variable.

ΔΔG^stability^ mCSM: 0.06 kcal/mol^-1^ (Neutral).

This mutation is present in 99.1% of all Delta sequences.

Whilst these residues do not appear to have much impact on structural stability they lie on a surface-exposed loop and P681 forms part of a furin motif (_681_PRRXR_685_) that can be processed by multiple proteases including TMPRSS2. There are reports that both P681H and P681R can lead to increased cleavage efficiency at S1/S2, possibly by providing greater accessibility to proteases, which results in increased fusion and syncytia formation (22–25).

**Figure 35.**
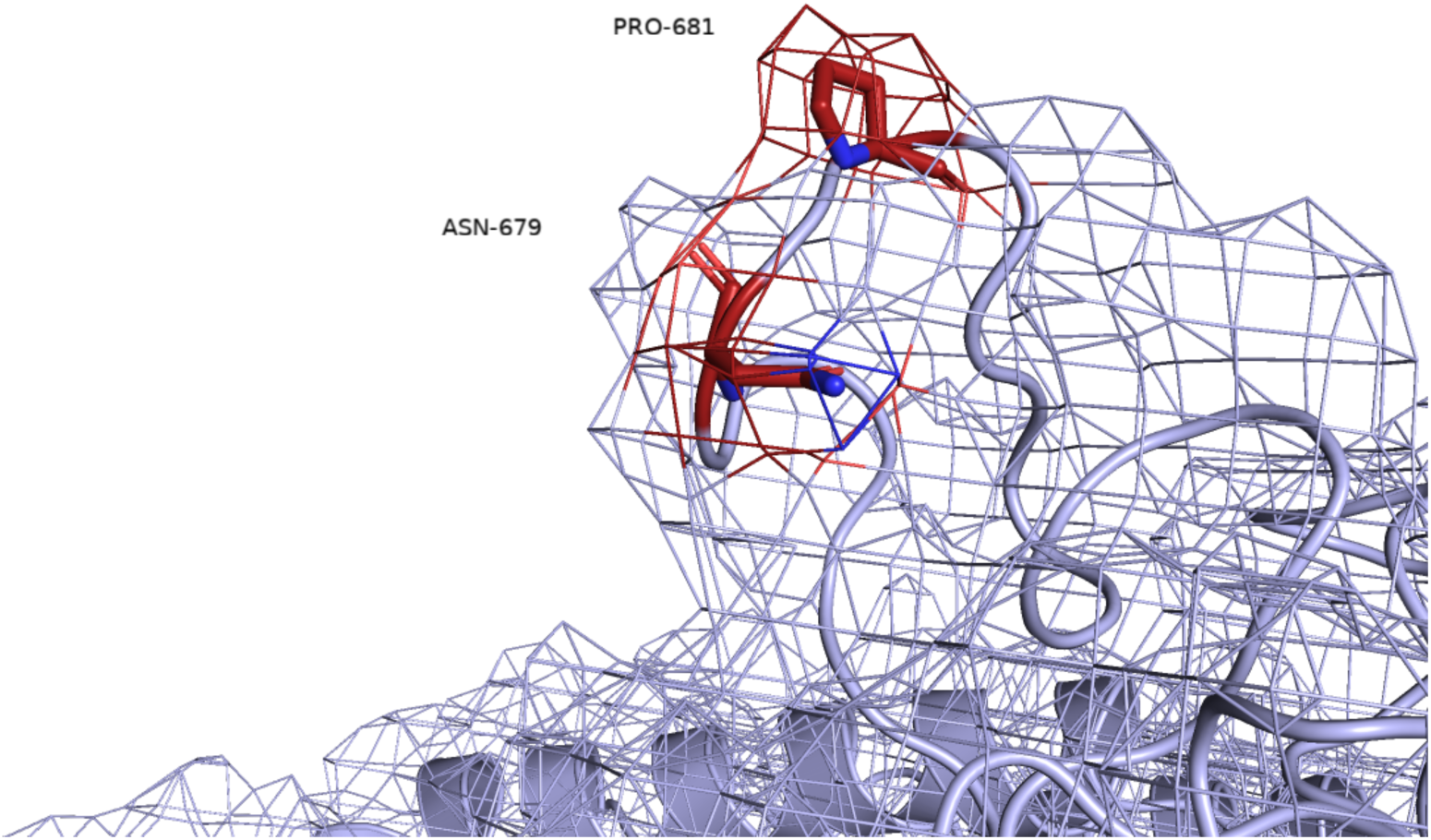
The S1/S2 cleavage site of S Protein. Close view of the S1/S2 cleavage site of S protein represented as a cartoon with the surface rendered as a mesh. Pro-681 and Asn-679 are highlighted in red. Source PDB structure 7fg3.

**Figure 36.**
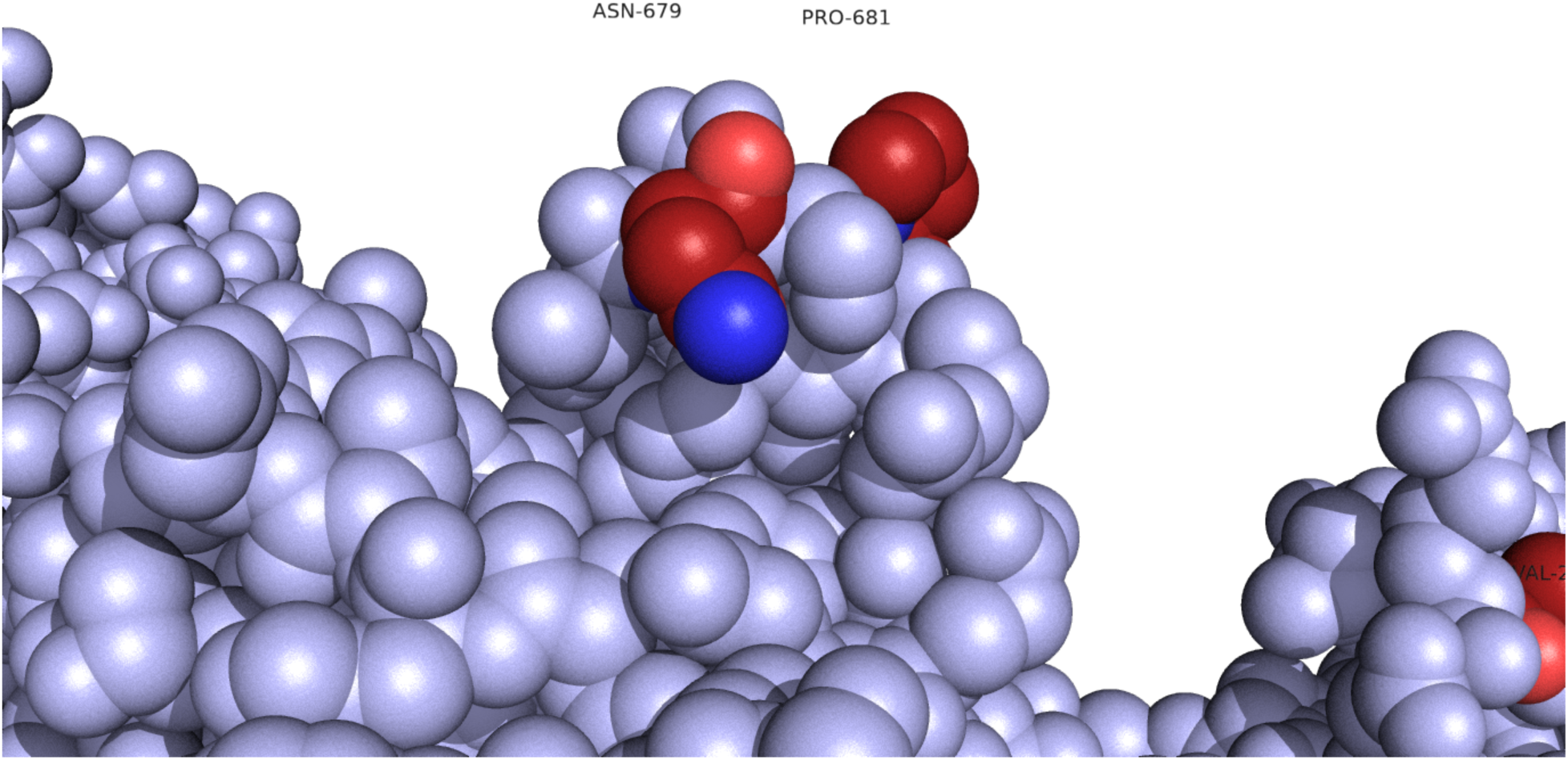
The S1/S2 cleavage site of S Protein. Close view of the S1/S2 cleavage site of S protein, represented as spheres. Asn-679 and Pro-681 are highlighted in red. Source PDB structure 7fg3.

S protein may interact directly with the JAK1-STAT1 pathway. There is some evidence that the S1 subunit interacts directly with STAT1, inhibiting ISRE promoter activation (26). There is no information available as of yet, as to the structural basis of this interaction with STAT1, or if it involves the NTD or RBD of S1, therefore we have considered both domains.

*S1-NTD*

BA.2 Mutation. L24S, always followed by Del25/27, sometimes recorded as A27S. The whole sequence of residues has a ConSurf score of 1. Highly variable. These mutations are present in 94.5% of all BA.2 sequences.

BA.1 Mutation. A67V. Alanine to Valine.

Blosum score of 0. Both hydrophobic.

ConSurf score of 1. Highly variable.

ΔΔG^stability^ mCSM: -0.09 kcal/mol^-1^ (Neutral).

This mutation is present in 95.8% of BA.1 sequences.

BA.1 Mutation. Del 69/70.

ConSurf scores of 1. Highly variable.

These deletions are present in 94.5% of all BA.1 sequences.

BA.1/Delta Mutation. T95I. Threonine to Isoleucine.

Blosum score of -1. Polar uncharged to hydrophobic.

ConSurf score of 4. Moderately variable.

ΔΔG^stability^ mCSM: -0.41 kcal/mol^-1^ (Destabilising).

This mutation is present in 37.9% of all Delta and 93.1% of all BA.1 sequences.

Delta/BA.1/BA.2 Mutation. G142D. Glycine to Aspartic Acid.

Blosum score of -1. Special case to negatively charged.

ConSurf score of 1. Highly variable.

ΔΔG^stability^ mCSM: -0.71 kcal/mol^-1^ (Destabilising).

This mutation is present in 66% of all Delta, 93.2% of all BA.1 and 97.9% of all BA.2 sequences.

BA.1 Mutation. Del 143/145.

ConSurf scores of 1 and 2. Highly variable.

These deletions are present in 93% of all BA.1 sequences.

BA.1 Mutation N211I. Asparagine to Isoleucine, typically followed by Del212.

Consurf scores of 1. Highly variable.

These mutations are present in 84.7% and 85.3% of all BA.1 sequences.

BA.2 Mutation. V213G. Valine to Glycine.

Blosum score of -3. Hydrophobic to special case.

ConSurf score of 1. Highly variable.

ΔΔG^stability^ mCSM: -0.57 kcal/mol^-1^ (Destabilising).

This mutation is present in 99.0% of all BA.2 sequences.

Delta Mutation. A222V. Alanine to Valine.

Blosum score of 0. Both hydrophobic.

ConSurf score of 1. Highly variable.

ΔΔG^stability^ mCSM: 0.26 kcal/mol^-1^ (Stabilising).

This mutation is only present in 10% of all Delta sequences.

The majority of these mutations occur on surface loops of the NTD within regions observed to be highly variable, presumably due to exposure to neutralising antibodies. There are also several deletion sequences, again located on loops which appear to be able to accommodate these changes. The one exception to this trend is the BA.1/Delta mutation T95I, which is part of a beta sheet strand, deeper within the domain. The destabilising effect of this mutation appears to be the result of clashes formed with surrounding residues.

However, our limited knowledge of STAT1 interactions with S1 makes it hard to draw firm conclusions on the significance of any of these mutations.

**Figure 37.**
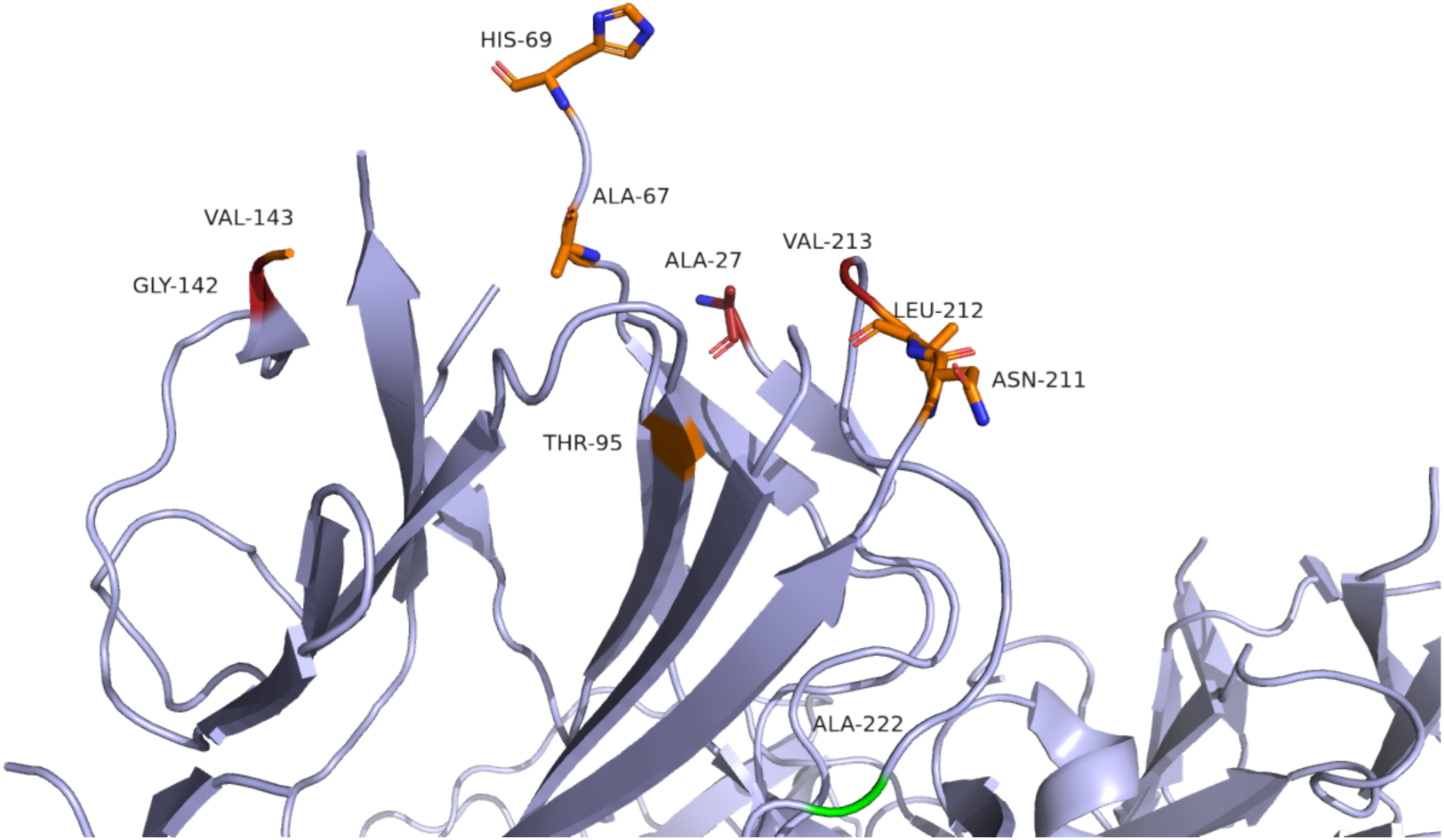
The N-terminal domain (NTD) of the S1 subunit of S protein. Cartoon representation of the S1-NTD. Residues of interest are highlighted in orange (BA.1) green (Delta) or red (BA.2). Source PDB structure 6vxx.

*S1-RBD*

BA.1/BA.2 Mutation. G339D. Glycine to Aspartic Acid.

Blosum score of -1. Special case to negatively charged.

ConSurf score of 1. Highly variable.

ΔΔG^stability^ mCSM: -0.49 kcal/mol^-1^ (Destabilising).

This mutation is present in 91.0% of all BA.1 and 97.1% of all BA.2 sequences.

BA.1 Mutation. S371L. Serine to Leucine.

Blosum score of -2. Polar uncharged to hydrophobic.

ConSurf score of 7. Moderately conserved.

ΔΔG^stability^ mCSM: -0.21 kcal/mol^-1^ (Destabilising).

This mutation is present in 82.2% of all BA.1 sequences.

BA.1 Mutation. S371F. Serine to Phenylalanine.

Blosum score of -2. Polar uncharged to hydrophobic.

ConSurf score of 7. Moderately conserved.

ΔΔG^stability^ mCSM: -1.00 kcal/mol^-1^ (Destabilising).

This mutation is present in 95.3% of all BA.2 sequences.

BA.1/BA.2 Mutation. S373P. Serine to Proline.

Blosum score of -1. Polar uncharged to special case.

ConSurf score of 4. Moderately variable.

ΔΔG^stability^ mCSM: -0.34 kcal/mol^-1^ (Destabilising).

This mutation is present in 83.1% of all BA.1 and 97.2% of all BA.2 sequences.

BA.1/BA.2 Mutation. S375F. Serine to Phenylalanine.

Blosum score of -2. Polar uncharged to hydrophobic.

ConSurf score of 8. Highly conserved.

ΔΔG^stability^ mCSM: -0.89 kcal/mol^-1^ (Destabilising).

This mutation is present in 82.9% of all BA.1 and 97.1% of all BA.2 sequences.

BA.2 Mutation. D405N. Aspartic Acid to Asparagine.

Blosum Score of 1. Negatively charged to polar uncharged.

ConSurf score of 2. Highly variable.

ΔΔG^stability^ mCSM: -0.82 kcal/mol^-1^ (Destabilising).

This mutation is present in 97.8% of all BA.2 sequences.

BA.2 Mutation. R408S. Arginine to Serine.

Blosum score of -1. Positively charged to polar uncharged.

ConSurf score of 6. Moderately conserved.

ΔΔG^stability^ mCSM: -0.08 kcal/mol^-1^ (Destabilising).

This mutation is present in 93.6% of all BA.2 sequences.

BA.2 (and BA.1?) Mutation. K417N. Lysine to Asparagine.

Blosum score of -1. Positively charged to polar uncharged.

ConSurf score of 9. Highly conserved.

ΔΔG^stability^ mCSM: 0.47 kcal/mol^-1^ (Destabilising).

This mutation is present in 94.5% of all BA.2 sequences (and 61.9% of all BA.1 sequences - unconfirmed if present in our BA.1 isolate).

BA.2 Mutation. N440K: Asparagine to Lysine.

Blosum score of -1. Polar uncharged to positively charged.

ConSurf score of 1. Highly variable.

ΔΔG^stability^ mCSM: 0.22 kcal/mol^-1^ (Stabilising).

This mutation is present in 87.3% of all BA.2 sequences (and 63.7% of all BA.1 sequences - unconfirmed if present in our BA.1 sequence).

Delta Mutation. L452R. Leucine to Arginine.

Blosum score of -2. Hydrophobic to positively charged.

ConSurf score of 1. Highly variable.

ΔΔG^stability^ mCSM: -0.92 kcal/mol^-1^ (Destabilising).

This mutation is present in 96.8% of all Delta sequences.

BA.1/BA.2 Mutation. Q493R. Glutamine to Arginine.

Blosum score of 0. Polar uncharged to positively charged.

Consurf score of 1. Highly variable.

ΔΔG^stability^ mCSM: -0.34 kcal/mol^-1^ (Destabilising).

This mutation is present in 84.3% of all BA.1 and 93.9% of all BA.2 sequences.

BA.1 Mutation. G496S. Glycine to Serine.

Blosum score of 0. Special case to polar uncharged.

Consurf score of 1. Highly variable.

ΔΔG^stability^ mCSM: -0.59 kcal/mol^-1^ (Destabilising).

This mutation is present in 80.6% of all BA.1 sequences.

BA.1/BA.2 Mutation. Q498R. Glutamine to Arginine.

Blosum score of 0. Polar uncharged to positively charged.

Consurf score of 1. Highly variable.

ΔΔG^stability^ mCSM: 0.17 kcal/mol^-1^ (Stabilising).

This mutation is present in 80.3% of all BA.1 and 92.4% of all BA.2 sequences.

BA.1/BA.2 Mutation. N501Y. Asparagine to Tyrosine. Bonding changes are significant.

Blosum score of -3. Polar uncharged to hydrophobic.

Consurf score of 1. Highly variable.

ΔΔG^stability^ mCSM: -0.37 kcal/mol^-1^ (Destabilising).

This mutation is present in 80.9% of all BA.1 and 92.6% of all BA.2 sequences.

BA.1/BA.2 Mutation. Y505H. Tyrosine to Histidine.

Blosum score of 2. Hydrophobic to positively charged.

Consurf score of 1. Highly variable.

ΔΔG^stability^ mCSM: -0.61 kcal/mol^-1^ (Destabilising).

This mutation is present in 81.3% of all BA.1 and 92.3% of all BA.2 sequences.

BA.1 Mutation. T547K. Threonine to Lysine.

Blosum score of -1. Polar uncharged to positively charged.

Consurf score of 4. Moderately variable.

ΔΔG^stability^ mCSM: -0.51 kcal/mol^-1^ (Destabilising).

This mutation is present in 98.2% of all BA.1 sequences.

Delta/BA.1/BA.2 Mutation. D614G. Aspartic Acid to Glycine.

Blosum score of -1. Negatively charged to special case.

Consurf score of 5. Middle of range.

ΔΔG^stability^ mCSM: -0.28 kcal/mol^-1^ (Destabilising).

This mutation is present in 99.3% of all Delta, 98.8% of all BA.1, and 99.9% of all BA.2 sequences.

BA.1/BA.2 Mutation. H655Y. Histidine to Tyrosine.

Blosum score of 2. Positively charged to hydrophobic.

Consurf score of 6. Moderately conserved.

ΔΔG^stability^ mCSM: 1.21 kcal/mol^-1^ (Stabilising).

This mutation is present in 98.5% of all BA.1, and 99.9% of all BA.2 sequences.

The majority of the mutations in the S1 subunit RBD are located on surface loops of the receptor binding motif (437-508), several of which have been noted as mutations of interest (L452R, S477N, N501Y) due to their modulation of binding affinity between S-protein and ACE-2 receptors, whilst K417N appears to reduce the neutralising capacity of sera (27). As with the NTD of S-protein there is no information about any specific interactions with STAT1 available to inform the structure models here.

**Figure 38.**
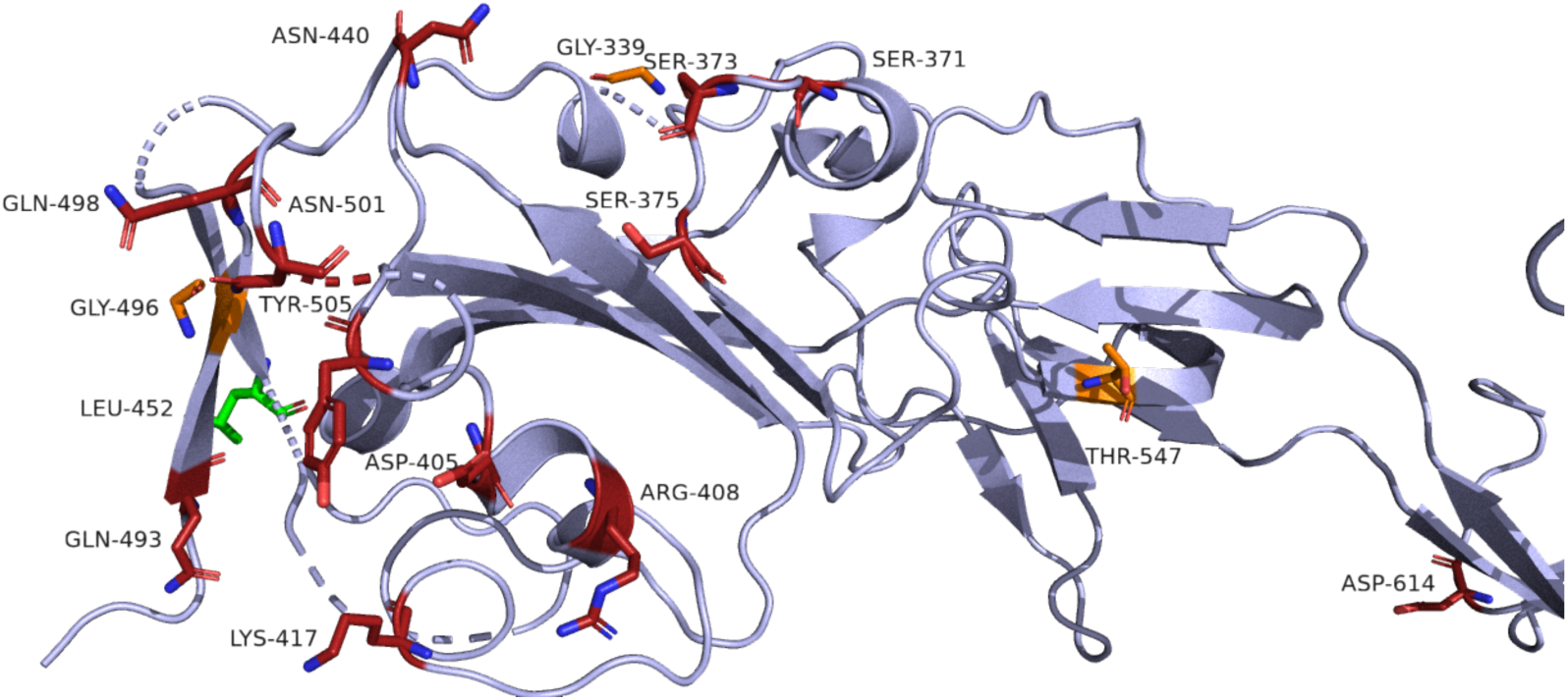
The Receptor Binding Domain of S-protein. Cartoon representation of the RBD of S-protein. BA.2 mutations are highlighted in red, BA.1 in orange, and Delta in green. Source PDB structure 6xvv.

